# Comparative connectomics of the descending and ascending neurons of the *Drosophila* nervous system: stereotypy and sexual dimorphism

**DOI:** 10.1101/2024.06.04.596633

**Authors:** Tomke Stürner, Paul Brooks, Laia Serratosa Capdevila, Billy J. Morris, Alexandre Javier, Siqi Fang, Marina Gkantia, Sebastian Cachero, Isabella R. Beckett, Andrew S. Champion, Ilina Moitra, Alana Richards, Finja Klemm, Leonie Kugel, Shigehiro Namiki, Han S.J. Cheong, Julie Kovalyak, Emily Tenshaw, Ruchi Parekh, Philipp Schlegel, Jasper S. Phelps, Brandon Mark, Sven Dorkenwald, Alexander S. Bates, Arie Matsliah, Szi-chieh Yu, Claire E. McKellar, Amy Sterling, Sebastian Seung, Mala Murthy, John Tuthill, Wei-Chung A. Lee, Gwyneth M. Card, Marta Costa, Gregory S.X.E. Jefferis, Katharina Eichler

## Abstract

In most complex nervous systems there is a clear anatomical separation between the nerve cord, which contains most of the final motor outputs necessary for behaviour, and the brain. In insects, the neck connective is both a physical and information bottleneck connecting the brain and the ventral nerve cord (VNC, spinal cord analogue) and comprises diverse populations of descending (DN), ascending (AN) and sensory ascending neurons, which are crucial for sensorimotor signalling and control.

Integrating three separate EM datasets, we now provide a complete connectomic description of the ascending and descending neurons of the female nervous system of *Drosophila* and compare them with neurons of the male nerve cord. Proofread neuronal reconstructions have been matched across hemispheres, datasets and sexes. Crucially, we have also matched 51% of DN cell types to light level data defining specific driver lines as well as classifying all ascending populations.

We use these results to reveal the general architecture, tracts, neuropil innervation and connectivity of neck connective neurons. We observe connected chains of descending and ascending neurons spanning the neck, which may subserve motor sequences. We provide a complete description of sexually dimorphic DN and AN populations, with detailed analysis of circuits implicated in sex-related behaviours, including female ovipositor extrusion (DNp13), male courtship (DNa12/aSP22) and song production (AN hemilineage 08B). Our work represents the first EM-level circuit analyses spanning the entire central nervous system of an adult animal.

## Introduction

For the body to respond to the higher processing commands of the brain, motor and sensory information must be transferred between the brain and nerve cord. In insects, there are 4 principal classes of neurons that traverse the neck. The three most numerous are: ascending neurons (ANs), which have their soma and dendrites in the VNC and feedback information to the central brain; descending neurons (DNs), which have the soma and dendrites in the brain and send commands via axons to the ventral nerve cord (VNC); and sensory ascending neurons (SAs), which have their soma outside of the VNC and send some sensory information directly from the periphery to the brain. Finally, a small number of motor neurons (MNs) exit the neck connective before reaching the nerve cord, directly targeting neck muscles in the periphery (Strausfeld et al., 1987).

Previous light microscopy (LM) and genetic studies in *Drosophila* have demonstrated that specific behaviours can sometimes be mapped to individual neurons and circuits. For the DN population, a range of behaviours are linked to individual or small groups of DNs: aDN, DNg11 and DNg12 for anterior grooming sequences (Guo et al., 2022; Hampel et al., 2015), DNa02 for turning (Rayshubskiy et al., 2020), DNp50/MDN for backwards walking (Bidaye et al., 2014), DNp01/GF for escape (Lima & Miesenböck, 2005), DNp07 and DNp10 for landing (Ache et al., 2019), DNp15/DNHS1, DNp20/DNOVS1, and DNp22/DNOVS2 for flight and neck control (Suver et al., 2016), and others. However, our understanding remains incomplete – only a few studies have examined larger groups of DNs by morphology (Namiki et al., 2018) or behaviour (Aymanns et al., 2022; Cande et al., 2018), and even less is known about ANs (Chen et al., 2023).

Until the advent of the male adult nerve cord (MANC) dataset, we could only estimate how many neurons connect the brain and the VNC by the available LM lines. 150 years after the first Golgi stainings (Golgi, 1873), the neurons of the neck connective as a complete population were described and typed for the first time, revealing 1328 DNs, 1865 ANs and 535 SAs in adult *Drosophila* (H. S. J. *. Cheong et al., 2024; Marin et al., 2023; Takemura et al., 2023). For comparison, Winding et al. (2023) have identified 182 DNs in the *Drosophila* larva. However, connectomic datasets are still scarce and to reveal the degree of variation in *Drosophila* neuronal circuits, we need to compare EM datasets across individuals, sexes and developmental stages, as done previously in *Caenorhabditis elegans* (Cook et al., 2019; Witvliet et al., 2021). The first whole brain comparative connectomics study in adult *Drosophila* (Schlegel et al., 2023) used two female datasets, FAFB-Flywire (Dorkenwald et al., 2023; Schlegel et al., 2023; Zheng et al., 2018) and the truncated hemibrain dataset (Scheffer et al., 2020), to obtain initial insights into which connections between neurons are conserved across datasets and specimens, estimating the extent of biological variation in connectomes. Here, we conduct a comparative analysis of EM datasets across sexes.

Male and female *Drosophila* exhibit sexual dimorphism in their behaviour, mediated by dimorphic neuronal circuits in both the brain and the VNC. Differences exist both in the connections made by neurons that are shared between the sexes and in the presence of neurons that appear to be specific to one sex. The sex of each *Drosophila* neuron is determined genetically, primarily through the expression of the transcription factor genes double sex (*dsx*) and fruitless (*fru*). Studies on *fru* and *dsx* expressing neurons and dimorphic behaviours have revealed several sexually dimorphic neurons and small circuits in the brain and VNC (Auer & Benton, 2016; Pavlou & Goodwin, 2013). Females, for example, require oviDNs for egg laying (Ache et al., 2019; F. Wang et al., 2020a) and vpoDN to open their vaginal plate when accepting a male (K. Wang et al., 2021). In contrast male specific P1 central brain neurons control both intermale aggression and courtship steps such as wing extension (Hoopfer et al., 2015; von Philipsborn et al., 2011) while a set of DNs (pIP10 and pMP2) and VNC neurons (TN1a, dMS2, vPR9, dPR1, dMS9, and vMS12) act to coordinate time and shape of sine and/or pulse song (Lillvis et al., 2024; Shiozaki et al., 2023; von Philipsborn et al., 2011). However, there has not been any systematic EM level comparison of dimorphic neurons between males and females.

We now describe all the neck connective neurons of the adult female fly brain (FAFB-Flywire) (Chen et al., 2023; Dorkenwald et al., 2023; Schlegel et al., 2023) and the female adult nerve cord (FANC) (Azevedo et al., 2022; Phelps et al., 2021), and compare them to the MANC dataset (H. S. J. *. Cheong et al., 2024; Marin et al., 2023; Takemura et al., 2023). We present the strategies developed to bridge physically disconnected datasets (brain and VNC) and compare datasets of different sexes. Our work represents the first atlas of DNs, ANs and SAs based on EM connectome data from the brain and the VNC. We then illustrate the utility of this complete and comprehensively annotated resource by addressing three scientific questions: first, we investigate the types of sensory information processed by DNs in the brain and the connections between ANs and DNs in the brain and nerve cord. Second, we explore stereotypy across the three datasets at the level of morphology and connectivity. Last, we examine the implications of unmatched neurons across datasets, particularly in the context of sexual dimorphism.

## Results

### Matching neurons across three datasets

We reconstructed all of the neurons that traverse the neck connective in the female adult fly brain (FAFB-Flywire) and the female adult nerve cord (FANC) datasets; we then compared them to the previously published male adult nerve cord (MANC) dataset (H. S. J. *. Cheong et al., 2024; Marin et al., 2023; Takemura et al., 2023) (Fig. 1, Supplemental file 1 for FAFB and FANC seed planes). We find that in the three datasets there are between 1315 and 1347 DNs that transmit motor commands and other information from the brain to the VNC; between 1733 and 1865 ANs that report processed sensory and motor state information from the VNC back to the brain; and between 535 and 611 SAs that convey sensory information directly from the periphery to the brain (Fig. 1b). The position of these neurons in the neck connective is morphologically segregated, with DNs more dorsal, ANs more ventral, and the SAs localised in two main and two smaller bundles on each side (Fig. 1c, see black arrows and Extended Data Fig. 1). DNs and ANs were matched across the two sides into pairs or groups in all datasets, and matched between the male and female VNC by their morphology and connectivity (Fig. 1d,e).

**Fig. 1:**
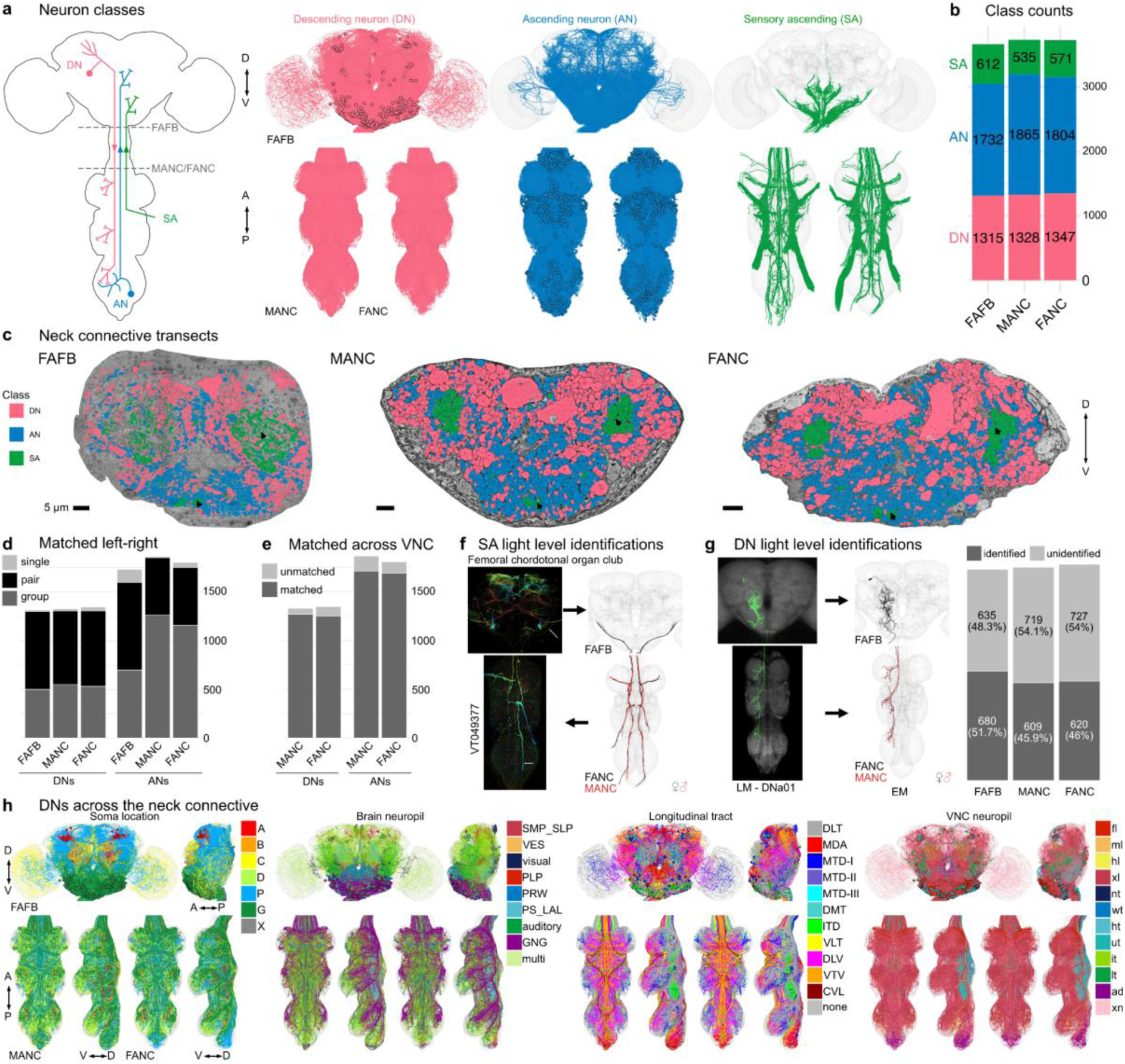
Reconstruction and identification of three neuronal classes across three datasets. **a**, Schematic of the central nervous system with the three neuronal classes that pass through the neck connective: descending neurons (DNs), ascending neurons (ANs) and sensory ascending neurons (SAs). FANC neurons are shown in MANC space here and in all following figures. **b**, Number of neurons in each class and dataset. **c**, Transects through the neck of the three datasets: Female Adult Fly Brain (FAFB), Male Adult Nerve Cord (MANC) and Female Adult Nerve Cord (FANC). These neck connective transects were used as seedplanes to find and reconstruct the three classes of neurons shown in different colours. **d**, Number of DNs and ANs that have been left-right matched into pairs or groups in the three datasets. **e**, Number of DNs and ANs that have a match across the two VNC datasets. **f**, SAs were assigned modalities by matching to light microscopy (LM) images. Left, an example of a LM image of Femoral chordotonal organ club. Right, the EM reconstructions that were matched to the image. **g**, DNs were identified in all three EM datasets by matching the EM reconstructions to LM level descriptions (mainly (Namiki et al. 2018), see supplemental file 2 - DN_identification). Left, an example of a LM image of DNa01 in the brain and VNC and next to it the FAFB, FANC and MANC EM reconstructions that were matched to those images. Right, the quantification of DNs identified in all three datasets. **h**, Identified DNs that can be matched across all three datasets coloured by soma location, brain neuropil, longitudinal tract and VNC neuropil. Please see attached files for a high resolution version of this figure.

We have contributed our proofreading and annotation of these neck connective neurons to the separate online platforms hosting each of these EM datasets as well as enabling a range of programmatic use (see Methods and Supplementary file 2); for example, both DN and AN type annotations are available for the brain in the online FlyWire connectome browser at codex.flywire.ai. However, we have found that comparisons across datasets are more powerful when each dataset can be visualised simultaneously in the same virtual space with a common interface for querying and viewing annotations. We have therefore provided access to co-registered and uniformly annotated neck connective neurons using the Neuroglancer viewer (Maitin-Shepard et al., 2021). This combined 3D web atlas can be viewed by following https://tinyurl.com/NeckConnective (see Methods – Neuroglancer Link for details).

Currently available EM datasets comprise either the brain or the VNC and are therefore truncated at the neck during specimen preparation. This creates a considerable challenge for matching the brain and VNC parts of the neurons sending projections through the neck. Matching existing light-level descriptions of these neurons to their EM-reconstructed counterparts is necessary for identifying these neurons across EM datasets, bridging brain and VNC, as well as linking the morphology to behavioural data. The ANs and SAs have recently been typed in the male VNC (Marin et al., 2023), but published LM information for these neurons is currently limited. ANs will require detailed matching with future light level resources but we were able to make some specific matches (see Fig. 4). However for SAs, a smaller and much less complex population, we were able to use comparisons with available LM images together with the position of tracts within the neck connective to assign gross sensory modalities in the brain as well as VNC (Fig. 1f, Extended Data Fig. 2; supporting evidence documented in Supplementary file 2 FAFB_SA_identification).

There is a significant amount of LM data for DNs (in contrast to ANs or SAs). In parallel work, we have recently described all DN axons in the male VNC EM connectome (H. S. J. *. Cheong et al., 2024) and matched some to earlier light level data of Namiki et al. (2018). By overlaying EM morphologies on these LM images, primarily sourced from Namiki et al. (2018) and a new LM collection, Namiki et al. (2024, manuscript in preparation), we were now able to identify 51.7% of FAFB and 46% of FANC DNs as well as increasing the proportion of LM identified DNs in MANC from 29% to 45.9% (supplemental file 2 - DN_identification). By separately matching the brain and VNC portions of a given DN to the same driver line, we were able to bridge connectome datasets. We could therefore analyse input and output, as well as compare between the male and female VNC (Fig. 1g). DNs have previously been grouped by different characteristics, and we demonstrate how soma location, longitudinal tract and neuropil innervation compare across the three datasets (Fig. 1h, Extended Data Fig. 3-6) (H. S. J. *. Cheong et al., 2024). As previously reported (H. S. J. *. Cheong et al., 2024; Namiki et al., 2018), soma location does not correlate strongly with other features such as axon tract and VNC neuropil innervation; we do note that neurons of the DNa and DNb soma groups innervate a combination of leg or upper tectulum neuropils, which matches reported functions for many of these DNs in steering during walking or flight (Extended Data Fig. 3a,b). Examining brain and VNC neuropil innervation patterns together highlights some interesting correlations. For example, DNs that innervate the SMP and SLP regions in the brain (higher order processing centres for olfactory stimuli) mainly target the abdominal ganglion of the VNC where they are likely to regulate reproductive or digestive functions (Extended Data Fig. 4a, 5i). DN axon tracts also highlight interesting groups: e.g. MTD-II tract DNs target the upper tectulum in the VNC (associated with wing pre-motor circuits) and receive input primarily from brain neuropils associated with multimodal integration and steering, the posterior slope (PS) and lateral accessory lobe (LAL). Nevertheless neuropil innervation and tract assignment are still quite coarse organisational features and are only a guide to the function or sensory input for any given DN. We therefore carried out a more detailed analysis of their sensory input.

### Sensory input onto descending neurons

By matching our EM reconstructions to LM data and linking them to previously published genetic or electrophysiological studies, we can make new predictions about DN functions and their circuits. DNs have diverse morphologies in the brain and VNC that can be uniquely identified in different EM datasets and LM lines (Fig. 2, see Methods – Light microscopy identification). Of the 223 DN types identified by LM data, just 2 could not be found in any of our EM datasets while a third turned out to be a duplicate; for 6 LM types we could identify a matching type in the brain but were unsure in the VNC (Fig. 2, Extended Data Fig. 7, supplementary file 2 DN_identification).

**Fig. 2:**
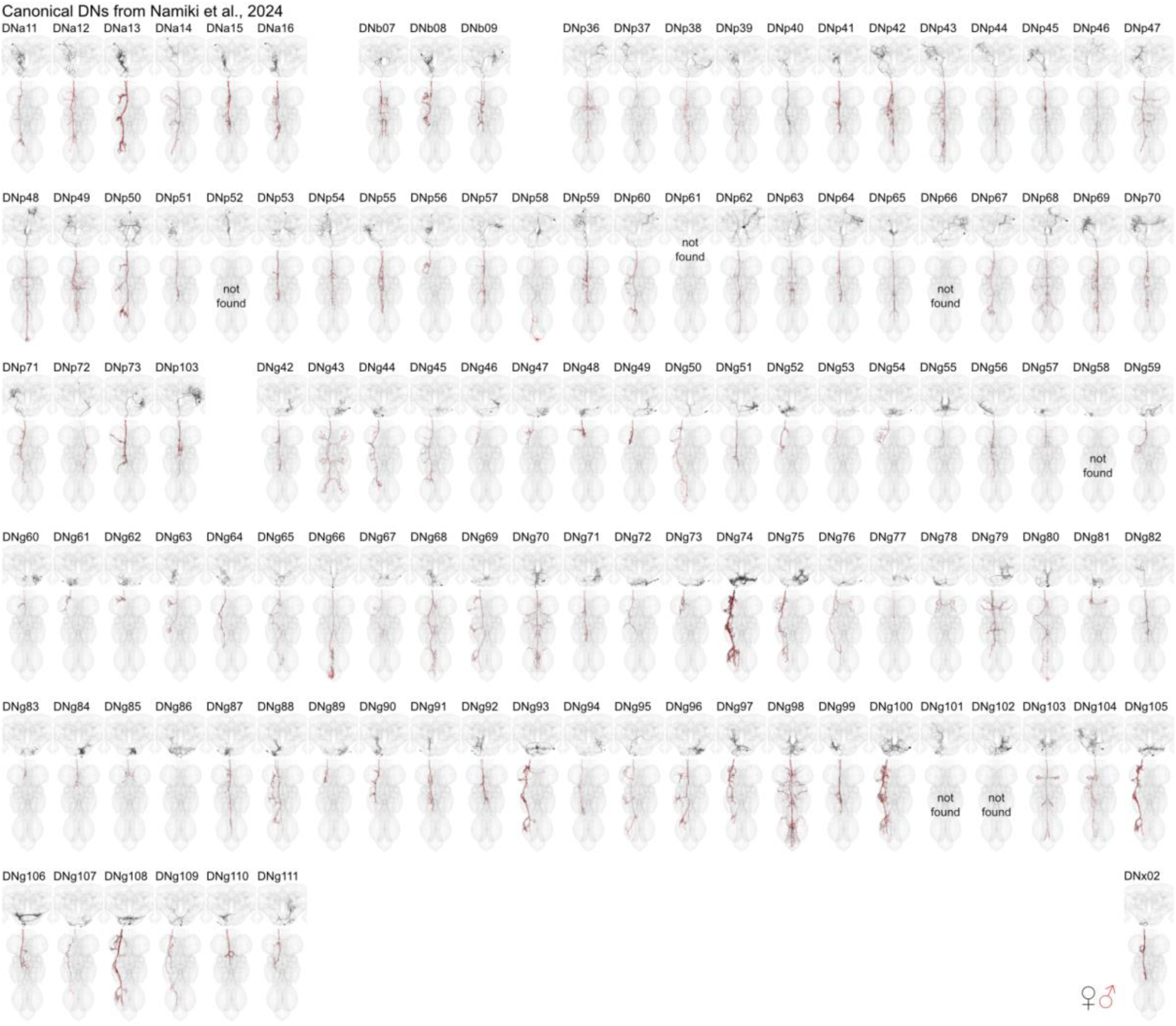
DN matching to Namiki et al. 2024 in prep. Morphology of identified DNs across all three datasets with nomenclature as described in Namiki et al. 2024 in prep. One DN type could not be found as it was duplicated (DNp61) and five could only be found in the brain (DNp52, DNp66, DNg58, DNg101, DNg102). See Extended Data Fig. 7 for matching to DNs previously characterised at light level by Namiki et al. (2018). See supplementary file 2 DN_identification for details. DN morphologies from the female datasets (FAFB, FANC) are in black, male dataset (MANC) is in red. This figure is also provided in high resolution and DNs can be viewed in 3D at https://tinyurl.com/NeckConnective. Please see attached files for a high resolution version of this figure.

Many DNs identified at light level have been associated with specific behaviours, sensory stimuli and evoked motor programs (Simpson, 2024). We have previously described the wide range of motor circuits targeted by the DN population in the male VNC . By bridging the neck connective, we can now analyse at scale the sensory information received by DNs in the brain and how that relates to the circuits targeted by their axons in the VNC. We began by analysing connectivity within the brain (Fig. 3). Summarising the input and output partners by neuronal class already revealed interesting patterns (Fig. 3a). Input is dominated by brain interneurons (class=central and visual projection). However, there are also strong inputs (∼10%) from at least two other sources: ANs and from DNs themselves. AN inputs are likely to convey a mix of processed sensory and motor state information from the VNC and there are long-standing hypotheses that these connections may be important for motor coordination (e.g. Chen et al., 2023). DN-DN connections were more unexpected. Although it has previously been reported that most DNs make axon collaterals (Namiki et al., 2018), commonly in the GNG area as they leave the brain, the large number of output connections in the brain (413,458 output synapses, Fig. 3a) was surprising since their principal axonal arbours are considered to be in the VNC. DN connections onto other DNs account for 42% of their total output in the brain but only 2% of their total output in the VNC (H. S. J. *. Cheong et al., 2024). This extensive DN-DN interconnectivity in the brain suggests the possibility of coordinated action across DNs, an idea that has recently been investigated by combining our connectome data with elegant functional studies (Braun et al., 2024). Intriguingly, many of these DN-DN connections are axo-axonic; whether this can result in direct excitation or inhibition of the downstream neuron or rather gates the axonal output of this neuron is unclear although Braun et al. (2024) were able to show that optogenetic activation of some DNs can propagate to others. The remaining DN output in the brain primarily targets central brain interneurons (45%), with functions likely including coordinating DN activity across the two sides of the brain. Finally DNs also make 8% of their brain output directly onto motor neurons, mostly controlling the proboscis; this is very similar to the fraction of direct output onto MNs in the VNC (H. S. J. *. Cheong et al., 2024).

**Fig. 3:**
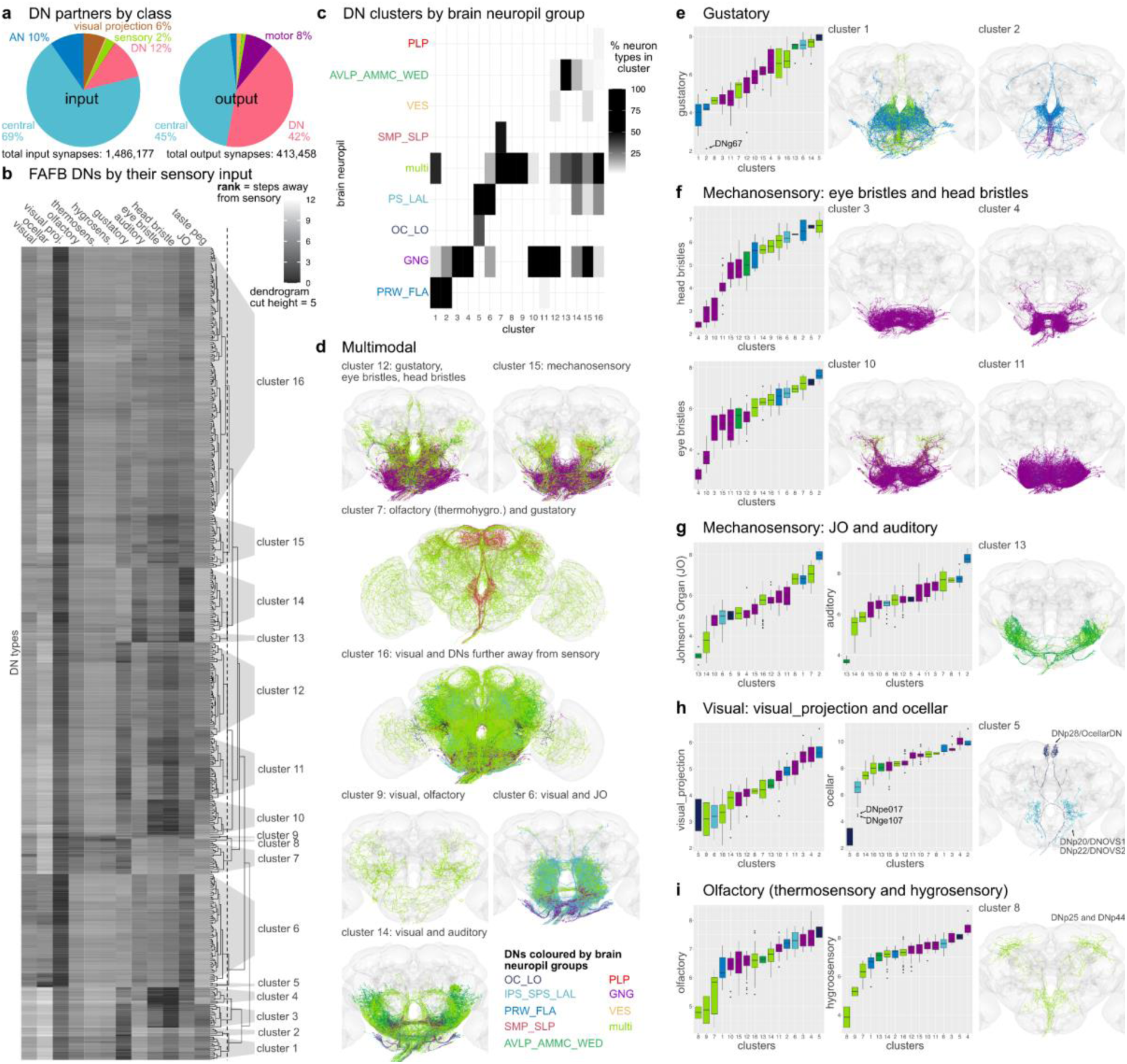
Sensory ranking of descending neurons. **a**, Pie charts show the neuron class composition of input and output partners of all DNs in the brain (corresponds to FlyWire super_class). Total synapse numbers shown below. **b**, Clustering of FAFB DNs by their sensory input rank (all apart from sensory DNs: DNx01, DNx02, LN-DNs). The ranks, ranging from 1 to 12, taken from (Dorkenwald et al. 2023), are defined as the traversal distances from a given sensory modality to each DN and then averaged by type. Low rank indicates a more direct connection from sensory modality to DN type. A cut height of 5 (dotted line in dendrogram in **b**) produces 16 clusters. **c**, Clusters shown in **b** by the brain neuropil assigned to DN types as a percentage of all types in that cluster. **d**, DN morphologies of clusters that are close in rank to several sensory modalities in the brain. **e-i**, DN morphologies of clusters that are close in rank to one particular sensory modality. Plots on the left show the average rank of the clusters defined in **b** for the different sensory modalities. Arrows point to specific DN types that stand out. DN morphologies are plotted in their brain neuropil colours.Please see attached files for a high resolution version of this figure.

DNs only receive a small fraction of their direct input from sensory neurons (2%). Therefore understanding the sensory modalities driving DNs requires more complex pathway analysis. Sensory neurons in the FAFB dataset have been annotated extensively and grouped into distinct sensory modalities: visual (photoreceptors and ocellar), olfactory, thermosensory, hygrosensory, auditory, mechanosensory eye bristles, mechanosensory head bristles, mechanosensory Johnson’s Organ (JO), and mechanosensory taste peg neurons (Schlegel et al., 2023). We looked at how far DNs are from these sensory modalities in the brain by using the information flow ranking established in Dorkenwald et al. (2023). This analysis excluded 4 DN types that are themselves sensory neurons (DNx01, DNx02, LN-DN1, LN-DN2). DNs were assigned to 16 clusters by their similarity in sensory input (Fig. 3b). DNs in each cluster typically have dendrites in the same brain neuropils (Fig. 3c, Extended Data Fig. 8 for FAFB neuropil assignments, supplemental file 2 - FAFB_DNs). For example, the two first clusters mostly arborize in the prow and flange (Ito et al., 2014), brain regions that receive taste information from the proboscis; these DNs are the closest (lowest average rank) to gustatory sensory neurons based on their modality (Fig. 3b, c, e). Seven DN clusters receive a combination of sensory modalities (Fig. 3d) and nine smaller clusters are specific to gustatory, bristle, auditory, ocellar or olfactory sensory information (Fig. 3e-i).

By looking at the DNs previously linked to specific behaviours and from what we know about different neuropils in the *Drosophila* brain, we can assess whether these clusters and the sensory modalities assigned to them are appropriate. DNs in cluster 7, for example, integrate olfactory and gustatory information (Fig. 3d); this cluster includes the oviposition-promoting oviDN neurons, which would require this type of sensory information to select a nutrient-rich food source (F. Wang et al., 2020a). Interestingly, most DNs in cluster 6, which receive a combination of visual and auditory information, innervate the posterior slope (IPS,SPS) and lateral accessory lobes (LAL), brain regions shown to be involved in higher order sensory processing and steering (Currier & Nagel, 2020; Namiki & Kanzaki, 2016; Steinbeck et al., 2020). These two sensory cues are essential for the navigation behaviour linked to DNs in this cluster, such as DNb06, allowing the fly to turn away from or towards a sound or a visual stimulus (Yang et al., 2023).

DNs close to mechanosensory inputs fall into three main groups. Four clusters (3,4,10,11) are close to eye and head bristles (Fig. 3f), two (13 and 14) are close to a combination of JO and auditory sensory neurons (Fig. 3g), and cluster 15 receives input from a combination of mechanosensory modalities. This is in agreement with the neuropil innervation that is more GNG-based for the DNs close to bristle sensory inputs and more WED, AMMC and AVLP for the DNs linked to auditory and JO information. Based on the neuropils innervated, this large group of DNs is likely to be responsible for the highly targeted grooming of the corresponding bristle locations (Eichler et al., 2024). Most DNs are close in raking to visual projection neurons, but only one small cluster (cluster 5, Fig. 3h), containing 3 DN types (DNp28/OcellarDN, DNp20/DNOVS1 and DNp22/DNOVS2), is specific for visual sensory information from the Ocelli (*Drosophila* have three Ocelli that sense light in addition to the compound eye). Both DNOVS1 and DNOVS2 additionally receive input from optic lobe output neurons that encode pitch-associated or roll-associated optic flow and are involved in fast flight and neck motor control (Suver et al., 2016). Olfactory, thermosensory and hygrosensory information converge onto the same clusters, especially onto cluster 8 (Fig. 3i). There are only 2 DN types in this cluster, DNp25 and DNp44; both were previously suggested to be close to olfactory sensory inputs in the hemibrain dataset (Schlegel et al., 2021) and DNp44 to hygrosensory inputs in FAFB (Marin et al., 2020). The lack of big DN clusters associated specifically with vision or olfaction suggests that this sensory information is more likely to be integrated with other sensory modalities, i.e. olfactory with gustatory, visual with auditory, or preprocessed in higher brain regions further away from DNs, cluster 16 (Fig. 3d). Thus, we are able to separate DN groups by their most likely sensory modalities, retrieve groups of DNs by their brain innervation, neuropil and sensory integration, and, by matching them to light level lines, follow them into the VNC, offering insights into the type of sensory information conveyed to the VNC and its potential functions.

### DN and AN interactions

To guide sequences of behaviour to given stimuli, we expect there to be feedback from the VNC ANs back onto DN circuits in the brain. Strong direct connections between DNs and ANs are uncommon in the VNC and the brain (arrows point to connections with the highest weight in Fig. 4a,b, weight = number of synapses). However, there is one exception that stands out in the VNC: DNx02 output onto AN06B025, which is strong in the number of synapses as well as percent of the total synaptic output (Fig. 4b,c). DNx02 are sensory descending neurons, two on each side, that enter the brain via the occipital nerve, a nerve that was only recently identified (Eichler et al., 2024). Analogously to DNx01, which responds to mechanosensory stimuli on the antenna and is serial to the bilateral campaniform sensillum (bCS) neurons, we predict that DNx02 would respond to mechanosensory stimuli from the eye, potentially with grooming behaviours (Aymanns et al., 2022; Namiki et al., 2018). DNx02 neurons in the brain stay within the GNG, while in the VNC they project into the neck and haltere neuropils (Fig. 4d). The effective connectivity of DNx02 shows the strong connection onto neck and haltere MNs, both ipsi- and contralateral, 2-3 layers into the VNC (Fig. 4c). We were able to find a line for AN06B025 and identify it in the brain (Fig. 4d, supplemental file 2 - AN_identification). We studied the DNx02 and AN06B025 circuit across the neck connective (Fig. 4e). This revealed that the top target of DNx02 is AN06B025 not only in the VNC but also in the brain. Neurotransmitter types have been predicted using a convolutional neural network for FAFB-Flywire and MANC trained on a set of neurons with identified neurotransmitter types (Eckstein et al., 2024; Takemura et al., 2023). AN06B025 is predicted to be GABAergic in both datasets, suggesting that it in turn shuts off DNx02, a self loop motif that was not observed in the *Drosophila* larval dataset (Winding et al., 2023) (Fig. 4e).

**Fig. 4:**
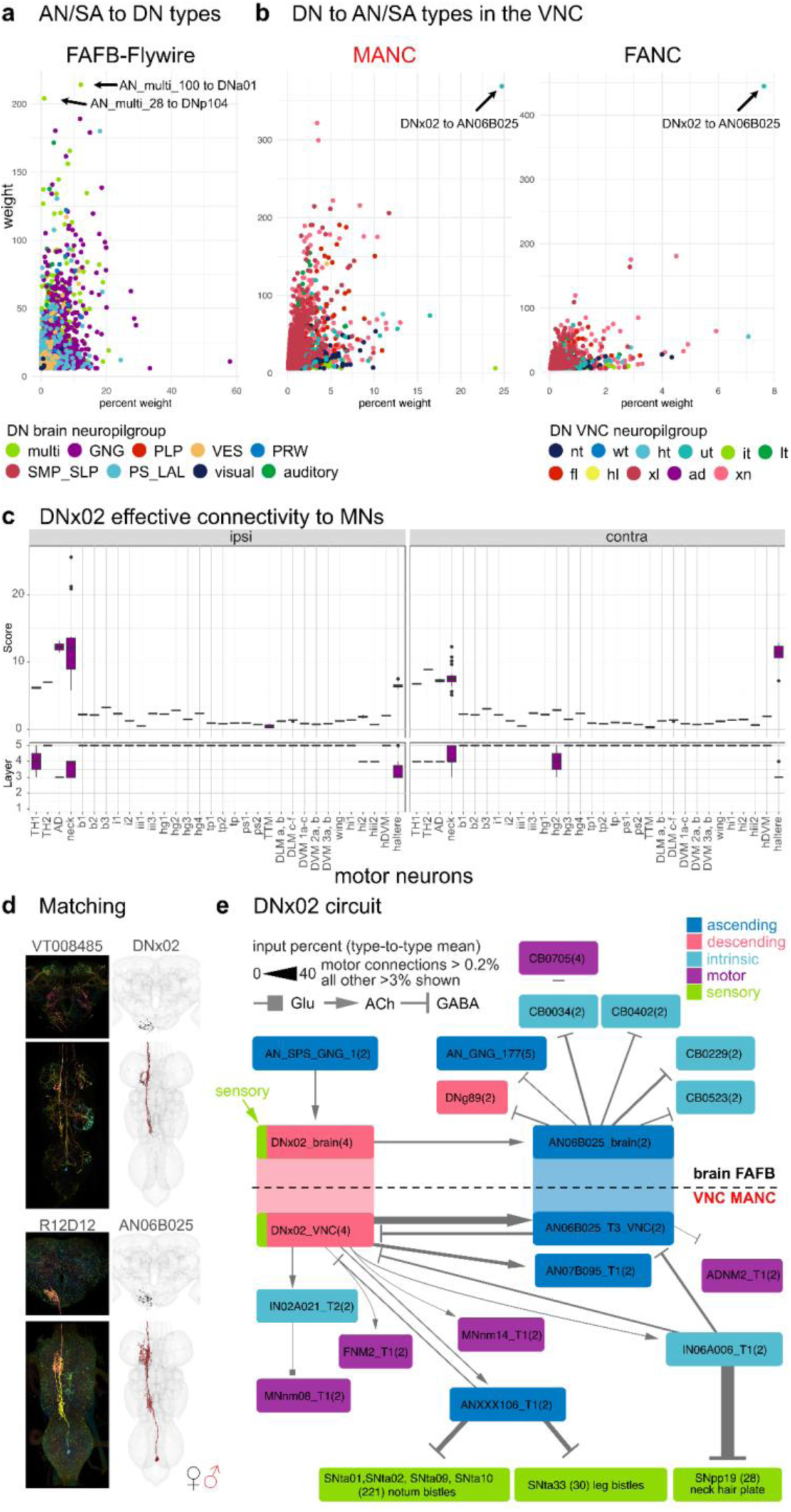
Direct descending and ascending neuron connections. Connectivity of AN/SAs to DNs and vice versa. a, Direct connectivity of AN/SAs onto DNs in the brain. DN and AN/SAs connections are averaged by type and plotted by mean weight in percent to mean weight. Arrows point to the two strongest connections in weight from AN/SAs onto DNs. b, Direct connectivity of DNs in the VNC onto ANs and SAs. Connections are averaged by type, like in a. Arrows point to the one connection that stands out in both MANC and FANC. The weight in a and b are the number of pre synapses. c, The effective connectivity to motor neuron targets ipsilateral and contralateral to the root side of DNx02. d, Morphology of DNx02 and AN06B025 in the brain and VNC. In black the EM morphology from female datasets (FAFB, FANC); in red from the male dataset (MANC). e, DNx02 circuit in the brain (FAFB-Flywire) and in the VNC (MANC). Connections in both datasets are averaged by type and shown in the percent input to the receiving neuron. Please see attached files for a high resolution version of this figure.

Additionally, DNx02 targets neck MNs directly and quite strongly via several hops. Two of the neck MNs, FNM2 and ADNM2, have been previously matched to data available from the blowfly, *Calliphora erythrocephala (H. S. J. *. Cheong et al., 2024; Strausfeld et al., 1987)*. The FNM2 in the blowfly is connected to the adductor muscle that moves the head both upwards and inwards while ADNM2 together with the cervical nerve motor neuron (CB0705) innervates the TH2 that controls yaw-movement of the head (Kauer et al., 2015; Strausfeld et al., 1987). We propose that DNx02 moves the head upwards and inwards in response to sensory stimuli. It is then inhibited by AN06B025 which inhibits the ADNM2 and disinhibits CB0705, both of which project to the TH2 muscle, potentially preparing for an additional sideways deflection of the head. Both movements could be part of a head grooming sequence. In the brain DNx02 has only one strong upstream partner, which also collects neck information from the VNC (AN_SPS_GNG_1/AN06B057, see supplemental file 2 - AN identification). Through one hop in the VNC DNx02 inhibits sensory neurons coming from bristles (leg and notum) and neck hair plate neurons via an ascending and intrinsic neuron, potentially dampening sensory information from these regions until the grooming movement is complete (Fig. 4e).

This is just one example of the kind of sensorimotor analysis made possible by our matching of brain and VNC neurons through the neck, across the entire central nervous system of *Drosophila*. Future EM datasets that contain a brain with attached VNC will make it possible to look at larger scale feedback loops and sequences of DN and AN activations required for serial behaviours such as grooming (Seeds et al., 2014).

### Stereotypy in the VNC

One important step for comparing EM datasets is to assess how stereotyped the morphology and connectivity of neuron types are across the two sides of an animal as well as between animals (Schlegel et al., 2023). From the 223 LM described DN types available to us, we matched all but 3 types in the brain and all but 6 types in both VNC datasets, supporting the commonly held observation that *Drosophila* neurons are highly stereotyped across individual animals (Fig. 2, supplemental file 2 - DN_identification). Matching across sides in the 3 datasets and across the two VNC datasets for DNs and ANs (Fig. 1d,e), even when not able to match to LM data, suggests a high degree of stereotypy in these two neuronal classes (matching in supplemental file 2). In addition, we have quantified their consistency in tract and VNC neuropil innervation (Fig. 5a,b, Extended Data Fig. 9,10). Nonetheless, there are a few cases in which the neuropil assignment does not agree across the two VNC datasets. We assume this is a combination of biological variation and the differences in the created neuropil meshes for the two different VNC datasets. For example, the DN type targeting specifically the middle leg neuropil in MANC (DNml) has a considerable amount (>5%) of its synapses in the front leg neuropil in FANC, and is therefore in the neuropil category xl (for multiple leg neuropil innervating) (Fig. 5b, highlighted green triangle). Examples of variability that we believe are due to differences in neuropil meshes between the datasets include some of the upper tectulum (ut) DNs in FANC that fall below our 80% synaptic output threshold and thus are assigned to multiple neuropil innervating (xn) (Fig. 5b, highlighted green triangle). Their morphology is, however, unique enough to match with high confidence across the two datasets (supplemental file 2 - FANC_DNs, MANC_DNs).

**Fig. 5:**
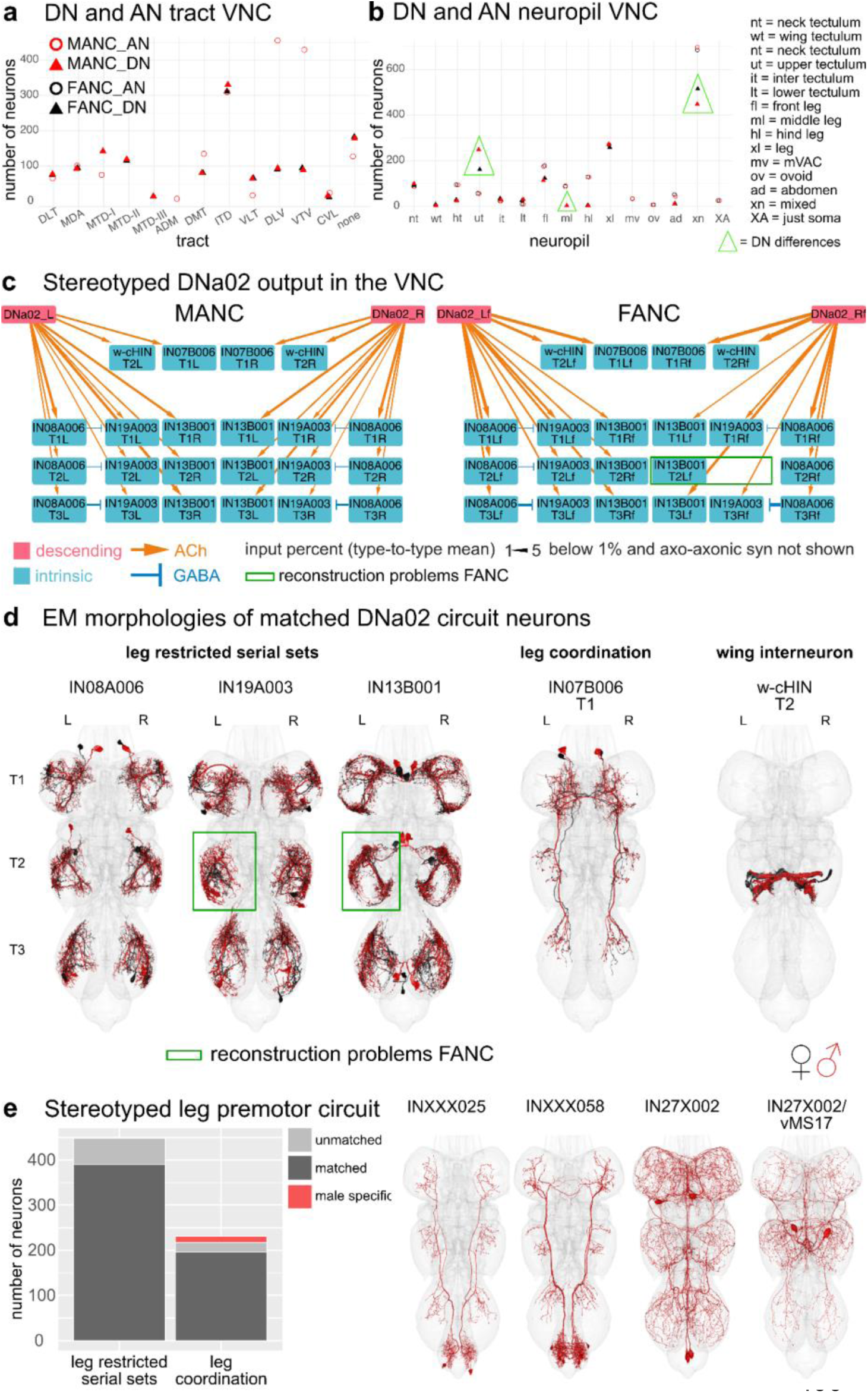
Across VNC dataset comparisons. a, Number of DNs/Ans assigned to a tract in MANC and FANC. b, Number of DNs/ANs assigned a VNC output/input neuropil in MANC and FANC. c, Example of a stereotyped circuit in the VNC. DNa02 in MANC and FANC in percent connect onto 3 sets of serial leg restricted neurons (IN08A006, IN19A003, IN13B001), the w-cHIN and a bilaterally projecting neuron (IN07B006). The types were matched across the two datasets and given the MANC type names accordingly. Downstream targets were selected by receiving more than 2% of DNa02 output. Arrow thickness corresponds to the percent input to the receiving neuron and only values above 1% are shown. d, EM morphologies of the neurons shown in the connectivity graphs in c. In black reconstructions from FANC, in red from MANC. e, MANC leg premotor circuit neurons published in (H. S. J. *. Cheong et al., 2024) matched to FANC. All leg restricted serial sets were found, although some are missing on one or the other side in FANC. All apart from 4 types of leg coordination neurons were matched to FANC. EM morphologies of those 4 unmatched types are shown on the right as potentially male specific neurons. Please see attached files for a high resolution version of this figure.

When normalising the number of synaptic connections using the percent input to the receiving neuron, we can recapitulate a previously analysed circuit of DNa02 (H. S. J. *. Cheong et al., 2024) in the FANC dataset (Fig. 5c, supplemental file 2 - other_MANC_FANC_matches), demonstrating that at a threshold of above 1% we identify the same downstream targets. The downstream partners consist of 3 serially repeated local neuron sets located in the leg neuropils, a bilaterally projecting neuron, and the w-cHIN neurons, which control wing MNs. The neurons were matched across the two datasets based on their morphology (Fig. 5d). The connection strengths across the two sides of the animal are comparable to the differences seen across the two datasets, with the exception of two neurons in FANC that are not well reconstructed (Fig. 5c,d, green box). Leg restricted serial sets are preferable when looking for stereotypy as we are able to match sets of 6 or 12 neurons per type (1 or 2 per leg neuropil), rather than individual neurons that might not be found because of reconstruction status or borderline significant differences in morphology. We concentrated on the previously published MANC leg premotor circuit (H. S. J. *. Cheong et al., 2024), as it includes 67 leg restricted serial types (448 neurons) and 75 leg interconnecting types (231 neurons) that are strongly connected to the serially repeated leg MNs. We matched these neurons by morphology and connectivity in the FANC dataset (Fig. 5e, supplemental file 2 - other_MANC_FANC_matches), and while there are many cases in which we have not yet found a match on one or the other side of FANC, we identified matches to all 67 serial sets (Fig. 5e). From the set of 75 interconnecting neurons we did not find any FANC neurons of the following 4 types: INXXX025, INXXX058, IN27X002 and IN27X002/vMS17 (Fig. 5e, morphologies on the right), suggesting that they might be male specific or sexually dimorphic in morphology to the extent that we cannot confidently match them. The first two types (INXXX025 = predicted cholinergic, INXXX058 = predicted GABAergic) project from the abdominal ganglion to the leg neuropils and would be good candidates for male specific leg movements in response to, for example, abdominal curling. The other two types are very similar to one another and have therefore been assigned to the same serial set and systematic type (IN27X002 and IN27X002/vMS17). The neuron with the T2 soma has been identified in LM as vMS17 and is reported to be involved in male courtship song (Lillvis et al., n.d.). Thus, we suggest that the other two neurons of this systematic type regulate similar male specific behaviours.

While complete matching of all VNC neurons across the female and male datasets is still in progress, the neurons we have matched so far (including DNs, ANs, SAs, and the leg premotor circuit, supplemental file 2) exhibit highly stereotyped morphology and connectivity across sides, neuromeres, and datasets. Until now the FANC (Azevedo et al., 2022; Lesser et al., 2024) and MANC (H. S. J. *. Cheong et al., 2024) datasets had only been analysed independently.

### Dimorphism

Some neurons could not be matched across male and female VNC datasets (Fig. 1e, MANC DNs = 59, FANC DNs = 97, MANC ANs = 155, FANC ANs = 115). This could be due to differences in connectome reconstruction, inter-individual biological variation or sexual dimorphism. Previous literature has already identified several sexually dimorphic (abbreviated to as sex. dimorphic in Fig. 6) and female or male specific neurons (sex-specific) (McKellar et al., 2019; F. Wang et al., 2020a, 2020b; K. Wang et al., 2021). In the following section, we will present known sexually dimorphic, and sex-specific DNs, such as the oviDNs, DNp13 and DNa12/aSP22. We defined potentially sex-specific DNs and ANs as neurons that are well reconstructed and can be confidently paired across the two sides of one VNC, but cannot be matched across the VNC datasets. We consider neurons to be potentially sexually dimorphic if they differ in morphology between the two VNC datasets, but have a morphologically consistent match across both sides of each nervous system. Details of individual neurons can be found in supplemental table 2 -dimorphic_DNs, dimorphic_ANs.

**Fig. 6:**
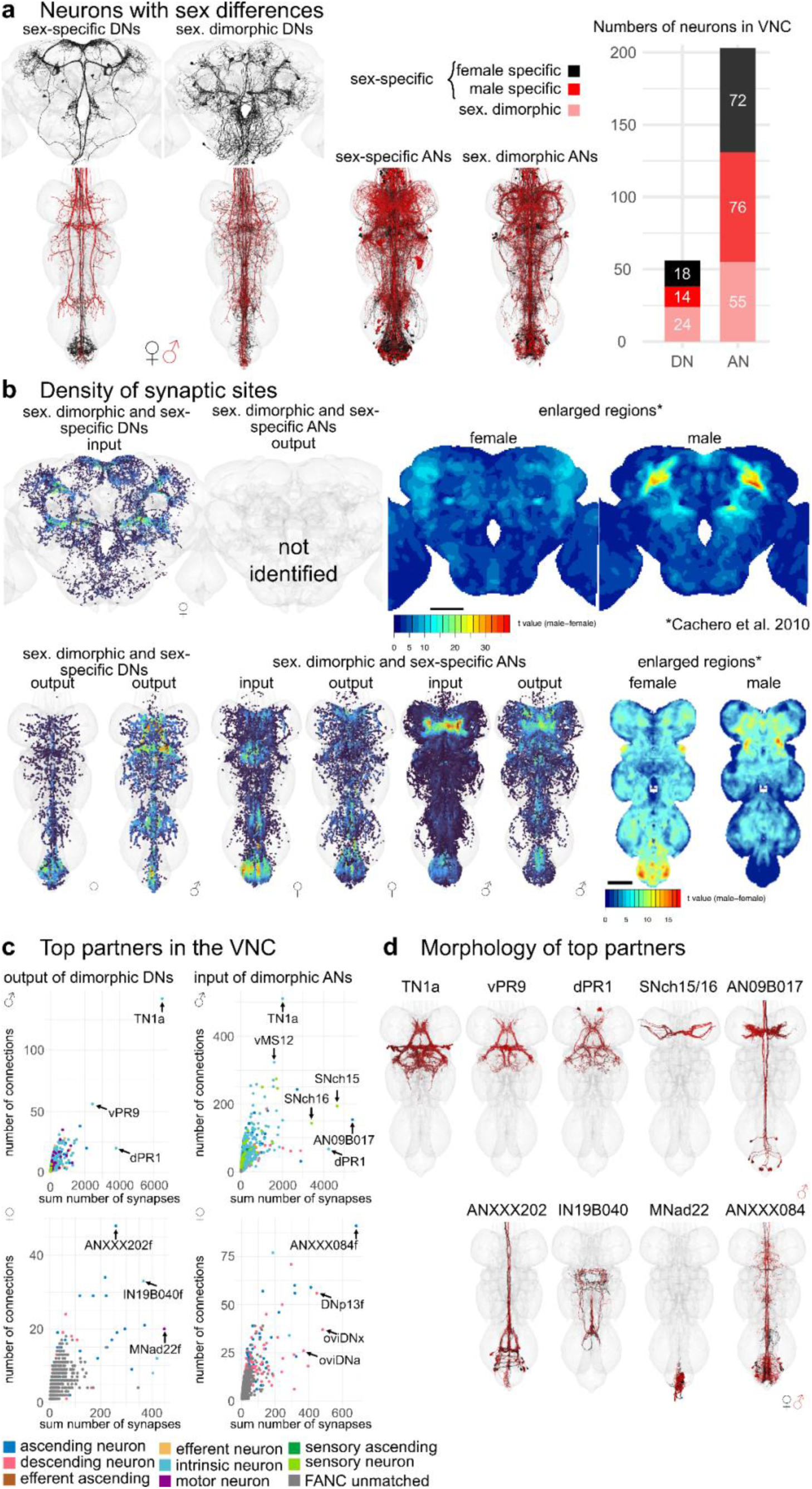
Sex-specific or sexually dimorphic (sex. dimorphic) neurons. **a**, Morphology of DNs in three datasets and ANs in the two VNC datasets that are sexually dimorphic or sexually specific (sex- specific) as described in the literature or predicted by the matching. In black the EM morphology from female datasets (FAFB, FANC), in red from the male dataset (MANC). **b**, Density of pre- or postsynapses of the DNs and ANs shown in **a** compared to previously published images of enlarged regions in the female and male central nervous system. **c**, Downstream or upstream partners of sexually dimorphic or sex-specific DNs or ANs respectively in FANC and MANC. Arrows point to the strongest partners by number of synaptic connections and number of neurons connecting onto them. **d**, Reconstructions of partner neurons in MANC (**c** top row) or in FANC that were matched to MANC neurons of that type. Please see attached files for a high resolution version of this figure.

Based on these definitions, we assigned 1% of DNs (18 female/14 male neurons) and 4% of ANs (72 female/76 male neurons) to be sex-specific, while 2% of DNs (24 neurons) and 3% of ANs (55 neurons) were sexually dimorphic in morphology (Fig. 6a). Using this information, we can identify the brain and VNC regions where sex-specific and sexually dimorphic neurons receive and send information (Fig. 6b). In FAFB-Flywire, we observed that input synapses to sex-specific DNs are present, especially in the protocerebral bridge, partially but confidently overlapping with previously published images of enlarged regions in the female fly brain (right panel Fig. 6b) (Cachero et al., 2010). The dimorphic DN output in the VNC also aligns with the enlarged abdominal ganglion in females and the distinct, dimorphic triangular region in males. In females, the input to dimorphic ANs is most pronounced within the abdominal region. Conversely, in males, the input to dimorphic ANs is concentrated in the T1 leg sensory area, as observed in Cachero et al. 2010 (Fig. 6b, right hand side).

Next, we wanted to see if dimorphic ANs/DNs tended to have the same upstream or downstream partners. We summed the number of connections to each partner type, as well as how many times a connection (above a weight of 5) occurred (Fig. 6c,d). The top output partners of dimorphic DNs are all part of the male song circuit: TN1a (silencing decreases sine song), vPR9 (silencing alters the amount of pulse and sine song) and dPR1 (silencing increases sine song) (Lillvis et al., 2024). Top input partners of dimorphic ANs include some of the song circuit neurons already mentioned, as well as two sets of sensory neurons coming from foreleg (T1) taste bristles (SNch15 and SNch16, both midline-crossing) (Marin et al., 2023; Possidente & Murphey, 1989); dimorphic ANs such as AN09B017 and AN05B035 also extensively interconnected in this T1 leg sensory area. AN09B017 are likely equivalent to the Fru positive vAB3 neurons that transmit pheromone signals from the front legs to P1 neurons in the brain (Clowney et al., 2015). This type of analysis is difficult without systematically defining cell types to define functional units of the nervous system. Our preliminary analysis of the FANC dataset found that the top output partners of dimorphic DNs may also be dimorphic. ANXXX202 has a dimorphic innervation pattern in the abdominal ganglion, IN19B040 has differences in morphology and the abdominal motor neuron MNad22 is dimorphic in connectivity. Input partners to dimorphic ANs include 3 interesting dimorphic DNs: oviDNa, oviDNx and DNp13, as well as the dimorphic ANXXX084 (see Fig. 6d for morphologies). Our analysis suggests that within the VNC male dimorphic DNs and ANs are heavily involved in the song circuit and the response to sensory information coming from the front legs, while female dimorphic DNs and ANs have important abdominal targets, and that the output activity of oviDNs in the VNC may be rapidly fed back to the brain via dimorphic ANs.

### Sexually dimorphic DNs involved in courtship and egg laying

The oviposition-promoting oviDNs are probably the best known female specific DNs. They have previously been divided into two subtypes based on LM data (F. Wang et al., 2020a). By direct comparison with EM reconstructions, we have defined six female oviDN types and one male type (Fig. 7a,b). Existing LM data were insufficient to match two of the oviDNs across the brain and VNC; EM data clearly distinguish these as two separate types (see Fig. 7a unmatched types). In addition to the oviDNs, we identified one female specific DN, vpoDN/DNp37, which has previously been described as important in female receptivity by controlling opening of the vaginal plate (K. Wang et al., 2021). We also identified 6 types of male specific DNs in the VNC. Two of these, pMP2 and pIP10 (Kohatsu et al., 2011; Shirangi et al., 2016; von Philipsborn et al., 2011), have previously reported functions and light microscopy lines that will allow identification in a future male brain dataset (Nern et al., 2024). We also identified 11 DN types which can be found in both male and female datasets but appear sexually dimorphic; for 7 of these (DNa08, aSP22, DNp13, pIP9, DNp48, LH-DN1 and -DN2), light level matches provide additional evidence for sexual dimorphism and have been matched in the two VNC datasets and in the female brain (Fig. 7f).

**Fig. 7:**
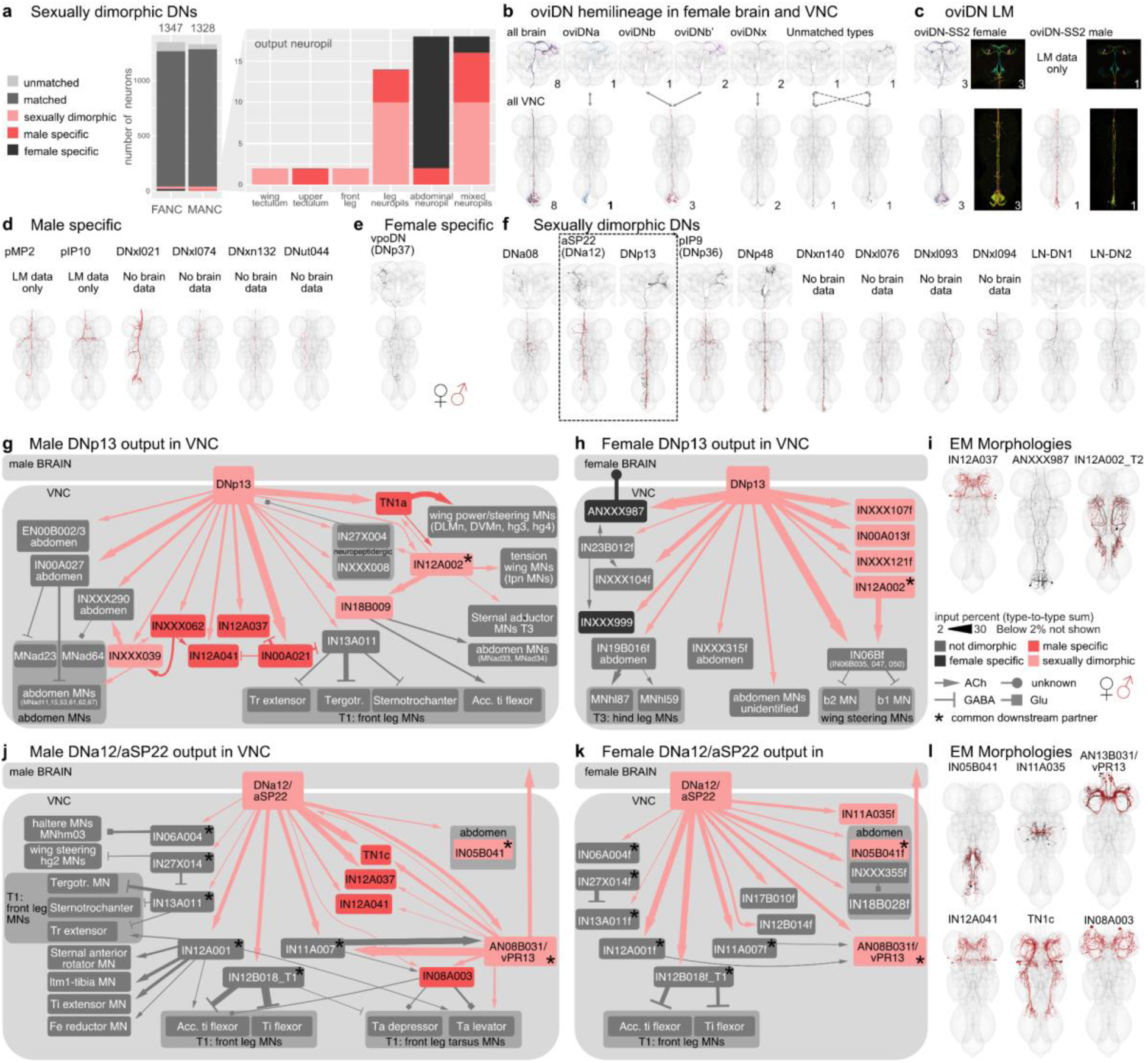
Sexually dimorphic and sex-specific descending neurons. **a**, Proportion of DNs that are sex-specific or sexually dimorphic by dataset and primary input neuropil. **b**, Morphology of DNs in the three datasets belonging to the oviDN hemilineage. **c**, EM morphologies identified within the LM images from female and male of the oviDN-SS2 line. **d**, EM morphologies of previously LM characterised male specific DNs and new potentially male specific DNs. **e**, EM morphology of the female specific DN, vpoDN (DNp37). **f**, EM morphologies of LM characterised sexually dimorphic DNs and new potentially sexually dimorphic DNs. **g**,**h**, Connectivity downstream of the sexually dimorphic DNp13 in MANC (**g**) and FANC (**h**). There is just one downstream partner in common between the two sexes, IN12A002 marked with a *. All other partners are either sex-specific (coloured black or red), are dimorphic in their connections (pink) or are not downstream of DNp13 in the other dataset (coloured grey). **i**, EM morphology of some of the top VNC targets. **j**,**k**, Connectivity downstream of the sexually dimorphic DNa12/aSP22 in MANC (**j**) and FANC (**k**). There are 8 downstream neurons in common. Only T1 leg motor neurons (MNs) have been systematically identified between the two datasets, thus other FANC Leg MNs are not shown. **l**, EM morphology of some of the top VNC targets. In black the EM morphology from female datasets (FAFB, FANC), in red from the male dataset (MANC). * indicates shared partners downstream of dimorphic DN pairs. Please see attached files for a high resolution version of this figure.

After comprehensive identification of sexually dimorphic DNs, we then carried out analysis of their downstream connectivity. We focused on dimorphic DNs making connections outside of the abdominal ganglion of the VNC due to reconstruction issues in this region of the FANC dataset. We selected DNa12/aSP22 and DNp13 for detailed study; both are fully proofread, present in both sexes, dimorphic in morphology, and have reported roles in both male and female mating behaviours (McKellar et al., 2019; Mezzera et al., 2023; F. Wang et al., 2020b). Robust comparative analysis of the downstream connections from DNs onto VNC neurons requires that these neurons have been both adequately proofread and matched across datasets. Starting from the automated segmentation (Azevedo et al., 2022), at the time of writing the FANC community has proofread just over 5000 neurons (including the 1804 ANs reported in this study) out of approximately 16,000 neurons documented in the VNC (Marin et al., 2023). Prior to this work, there has been relatively little matching of precise cell types between the FANC and MANC datasets, with the notable exception of foreleg MNs and wing MNs (Azevedo et al., 2022; H. S. J. *. Cheong et al., 2024). Through our detailed analysis of the MANC and FANC neck connective neurons, combined with new computational approaches for intrinsic neurons, we have now matched over 4000 neurons across the datasets including top targets of these DNs (supplemental file 2 - other_MANC_FANC_matches, see Methods).

Although identifiable as the same cell type across males and females, DNp13 has highly distinctive morphology and connectivity (Fig. 7f). In both sexes their downstream circuits ultimately target MNs in the wing neuropil and abdominal ganglion. However, these MNs are of distinct types and are targeted via different VNC interneurons in each sex. The top partner of DNp13 in MANC is the male specific doublesex-positive neuron TN1a, which is particularly important for sine song (Shirangi et al., 2016), however, we also find that there are several strongly connected wing MNs directly downstream (Fig. 7g). DNp13 in females has been shown to respond to courtship song, and activation of DNp13 leads to ovipositor extrusion (F. Wang et al., 2020b). Unsurprisingly, abdominal MNs are top targets of DNp13 in FANC (Fig. 7h); however, we were surprised to see that the top partners of DNp13 in FANC are three very similar looking IN06B neurons (types: IN06B035, IN06B047, IN06B050) that output strongly to b1 and b2 wing MNs (Fig. 7h,i), suggesting that there might be a female wing phenotype that has not yet been described experimentally. This is reminiscent of recent observations that vpoDN may control a wing-spreading behaviour in the evolutionarily related Drosophilid *D. santomea (Li et al., 2023)*. The only common downstream target of DNp13 in males and females, with a 2% threshold, is IN12A002 (Fig. 7g,h black star, morphology shown in 7i). IN12A002 has similar morphology in both sexes, but different connectivity between the two circuits suggests that it is also dimorphic. The remaining intrinsic neurons targets (shown in grey Fig. 7g,h), receiving input from DNp13 in one sex, are present in both datasets but not directly downstream of DNp13 in the other sex.

DNa12, also known as aSP22, shows greater similarity in connectivity and morphology between the two datasets when compared to DNp13 (Fig. 7j,k). Activation of DNa12 elicits proboscis extension, front leg extension, spontaneous posture adjustments, and abdomen movements in both sexes (McKellar et al., 2019). Interestingly, the type of abdominal movement elicited by DNa12 differs between males (abdominal bending) and females (abdominal extension). In accordance with this work, we see that with a threshold of >2% input DNa12 neurons share 8/12 MANC and 8/13 FANC downstream partners. The partners that are not shared are also not present when considering all downstream partners with a synapse weight threshold >10. To confirm this, we additionally matched all downstream partners of the MANC DNa12 to those in the FANC dataset. All partners can be matched between the datasets except for the sex-specific TN1c, IN12A037, IN12A041, and IN08A003, which are not targeted by the FANC DNa12.

The connections to the front leg Tibia extensors can be found in both MANC and FANC, in line with the foreleg lifting seen in both sexes (McKellar et al., 2019). DNa12 connects to the sexually dimorphic AN08B031 in both sexes. In turn, AN08B031 has distinct connectivity in males and females. This connection provides an example of a sexually dimorphic AN-DN pair which forms sexually diverging circuits while preserving their connection with one another (Fig. 7j,k,l). As this AN cannot be linked to LM, we cannot say if it conveys the proboscis extension that has been described in both sexes in response to DNa12/aSP22 activation. The most surprising connection from the MANC DNa12 is to TN1c, a neuron shown to modulate pulse song (Shirangi et al., 2016) as a phenotype in song has not been reported at time of publication. DNa12 in MANC both directly as well as indirectly connects to TN1c via the sex-specific AN08B043 and AN13B031 (Fig. 7j,k). Two especially interesting neurons that are only downstream of DNa12 in FANC are IN05B041 and INXXX335. They target the abdominal ganglion and we suggest they are responsible for the dimorphic abdomen extension in females (Fig. 7k).

These two examples show that together with the available literature we can now use this LM matching to understand and compare EM circuits across the two VNC datasets. This helps us explain the phenotypes observed in behavioural and genetic activation experiments and link them to yet unstudied neurons in the VNC. Moreover, it gives us the chance to make new hypotheses about the function of these neurons, principally suggesting potential roles in song or wing movement that should be looked at experimentally for both DNp13 and DNa12.

### Sex-specific ANs of the 08B hemilineage

While the literature and experimental research have explored some sexually dimorphic and sex-specific DNs, there remains a notable gap in our understanding of ANs in general, particularly with respect to dimorphism. Work on ANs as a population has started to appear, for example ANs encoding behavioural states (Chen et al., 2023; McKellar et al., 2019) but only very few genetically identifiable AN cell types with LM lines are available: LAL-PS-ANs (Fujiwara et al., 2022) and Lco2N1/Les2N1D ANs (Tsubouchi et al., 2017), moonwalker ANs (Bidaye et al., 2014; Tsubouchi et al., 2017) and PERin ANs (Mann et al., 2013). We are not currently aware of any literature reporting sex-specific or sexually dimorphic ANs, therefore all ANs we have annotated are being reported here for the first time (morphologies shown in Extended Data Fig. 11). We caution that this label is putative: a definitive classification of these ANs as sexually dimorphic or sex-specific will require systematic identification and confirmation with LM data.

We identified the hemilineage and soma neuromere for all dimorphic ANs in FANC whenever possible, and compared the number of dimorphic ANs across the two datasets (Fig. 8a). We found that the hemilineages with the highest number of dimorphic ANs are the hemilineage 08B and the ANs in the abdominal ganglion, to which we were unable to assign a hemilineage in either MANC or FANC. To avoid reconstruction issues in the abdominal ganglion, we focused our analyses on sex-specific 08B neurons in both datasets. The sex-specific ANs, both in MANC and FANC, are the only sex-specific ANs that clearly innervate the mesothoracic triangle associated with dimorphic neurons involved in male song production, which include pMP2 or vPR1 (Yu et al., 2010) (Fig. 8b,c). The male specific 08B ANs consist of 7 types (Marin et al., 2023); here, we categorised the female specific 08B ANs into 6 types based on morphology and connectivity (Fig. 8d,e). For the male specific ANs, we observe that 5 of them form an interconnected circuit, including the dimorphic DNs pMP2 and piP10, intrinsic neuron dPR1 and sexually dimorphic AN19B007/dMS9. We infer that these are involved in the male song circuit as expected by their innervation pattern. AN08B020 and AN08B059, on the other hand, only have connections from two DNs, neither of which have been associated with song production at present, but which could be involved in a different male specific behaviour based on their innervation (Fig. 8d). The upstream circuit to female sex-specific 08B ANs does not include any of the neurons upstream of the MANC sex-specific 08B ANs. This supports the idea that these neurons differ both in terms of their morphology and connectivity. Another supporting factor is that the known sexually dimorphic fru+ pIP9/DNp36 (Yu et al., 2010) inputs onto two of the AN types. The other neurons upstream in FANC all exist in MANC but do not connect to any ANs of the 08B hemilineage.

**Fig. 8:**
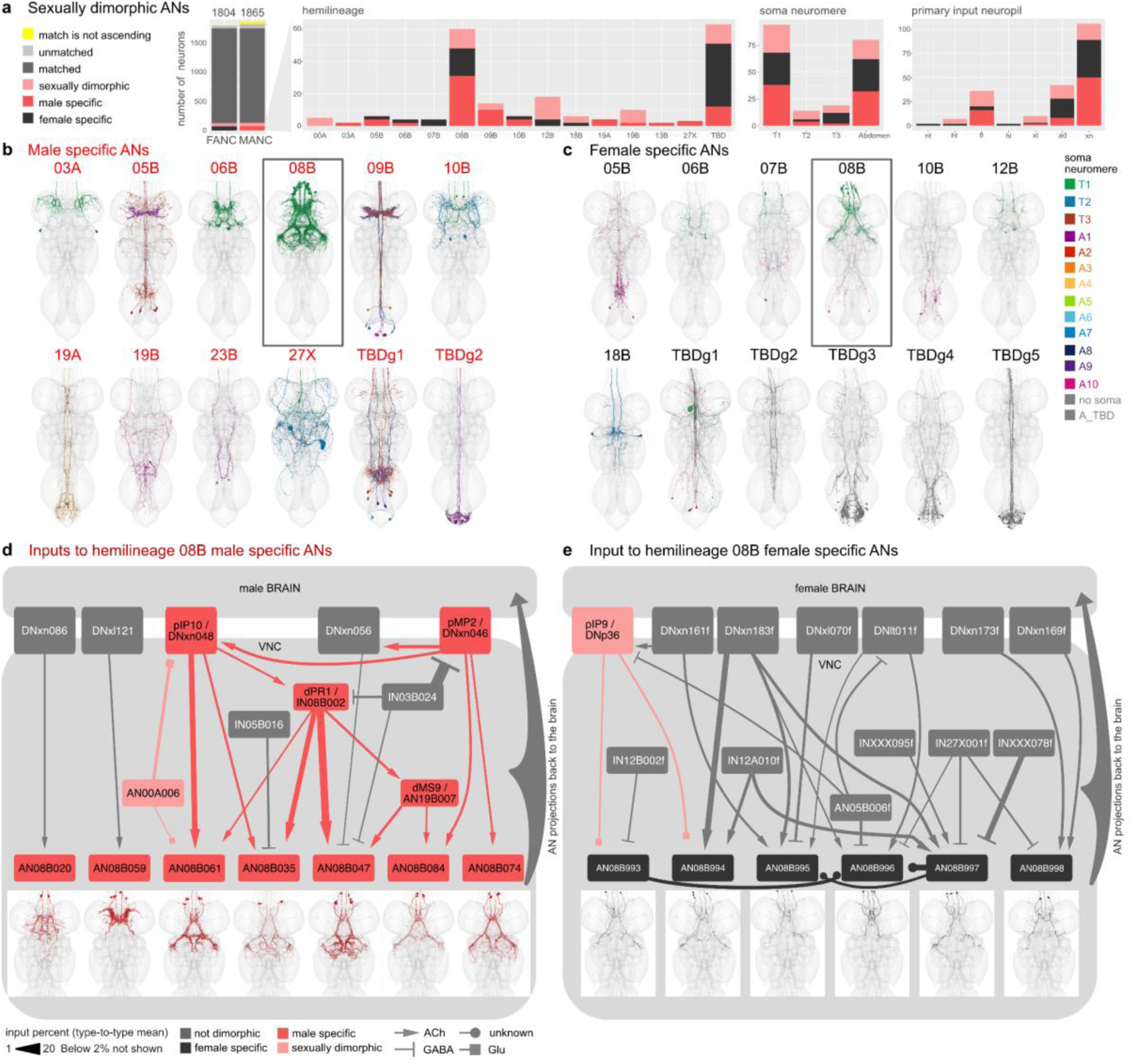
Sexual dimorphism and sex-specific ascending neurons. **a**, Proportion of ANs that are potentially sex-specific or potentially sexually dimorphic by hemilineage, soma neuromere and primary input neuropil. **b**,**c**, Morphology of ANs that are potentially sex-specific in males (**b**) and females (**c**) by hemilineage. FANC neurons were assigned hemilineages and soma neuromere if possible and given new type names. **d**,**e**, Input circuit in the VNC to potentially sex-specific AN types of hemilineage 08B with soma location in T1 (black box in **b** and **c**). Morphology of AN types underneath. All input neurons with more than 2% input onto the receiving AN are shown. FANC neurons in **e** were matched to MANC neuron types by morphology and connectivity and given the MANC names with an addition of f for female. Please see attached files for a high resolution version of this figure.

We conclude that 5 out of the 7 newly identified types of male specific 08B ANs are important for providing feedback during male song production. This may represent another example of ANs acting as a corollary discharge to suppress the auditory response to self-generated song (Poulet and Hedwig 2006; H. S. J. Cheong et al. 2024). Conversely, the female specific 08B ANs transmit feedback about sexually dimorphic information (such as sexual receptivity and post-mating state) back to the brain, a pathway which either takes a different course or does not exist in the male nervous system. Thus, we present two strategies of dimorphic circuits in the nervous system: 1) Sex-specific neurons interact with one another to produce a sex-specific behaviour, and 2) circuit elements present in both sexes interact with sex-specific neurons to establish representations which are necessary in one sex but not the other.

## Conclusion

Through detailed reconstruction of DNs, ANs, and SA neurons across three EM connectome datasets, we present the first complete set of these neuronal classes spanning the neck connective. We have categorised the neurons by sensory modality (for DNs and SAs in FAFB) and neuropil innervation across the brain and nerve cord, as well as tract and soma location. We have established a platform for systematic neuron typing based on light-level and cross-dataset identification, which we encourage the community to utilise for studying their specific circuits of interest. Early access to the proofread and annotated connectome data that we describe has already enabled and assisted a range of exciting work including studies of the circuit basis of locomotor behaviour and the organisational and functional logic of descending neurons (Braun et al., 2024; H. S. J. Cheong et al., 2024; Dallmann et al., 2024; Lee et al., 2024; Lesser et al., 2024; Sapkal et al., 2023; von Reyn et al., 2024 unpublished; Yang et al., 2023).

The sensorimotor reconstructions presented enable us to infer the circuit basis of behaviours, including sexually dimorphic patterns, allowing us to formulate new hypotheses regarding numerous circuit components. Future studies of any part of the CNS can now use this resource to link genetically-defined neurons, physiological responses or optogenetic manipulations to the connectome. While earlier studies often focused on a single neuropil of the brain, e.g. Antennal Lobe for odour processing or Mushroom Body for learning and memory, connectomics has already underlined that the sensorimotor circuits underlying behaviours are complex and brain-spanning. Future research can now adopt similar approaches across the whole CNS by integrating multiple datasets as we have now demonstrated. A recent connectome-constrained neural network of the motion pathways in the optic lobe of *Drosophila* reliably predicted measurements of neural activity from the connectivity of 64 cell types (Lappalainen et al., 2023); this approach relied on optimising the neural output of the modelled system to solve a behavioural task. Our neck connective work should enable modelling of the key output neurons of the brain and also models spanning the entire central nervous system.

Despite the common challenge of small sample sizes in connectomics, our study stands out for its use of datasets across individuals and sexes; prior work compared specific (isomorphic) circuits in the *Drosophila* larva (Gerhard et al., 2017; Valdes-Aleman et al., 2021) while recent adult work has compared two female brain connectomes (Schlegel et al., 2023). By systematically comparing DNs, ANs and SAs from both male (MANC) and female (FANC) VNC datasets, we categorised similarities and differences between the two sexes. This represents the first comprehensive comparison of *Drosophila* neuronal morphology and connectivity between sexes at EM resolution. We have identified all previously published dimorphic DNs and describe the circuits of DNa12 and DNp13 in both datasets. Moreover, we have excluded and annotated all differences that we believe are due to biological variation between individuals (variation in numbers, single, missing or additional neurons of a type) or reconstruction state in one of the two datasets. Our findings suggest potential sex-specific or sexually dimorphic DNs and ANs, with a specific focus on circuits of sex-specific ANs from the 08B hemilineage associated with male song during courtship. This now lays the groundwork for understanding circuits for sex-related behaviours all the way from the sensory periphery, through higher brain processing to motor ouput.

## Methods

### Neck connective proofreading and annotation

We defined a perpendicular plane through the neck connective posterior to the cervical nerve for the two VNC datasets (MANC and FANC), and anterior to the cervical nerve for the FAFB dataset. Supplemental file 1 includes two tables, FANC_seed_plane and FAFB_seed_plane, that list all profiles with their xyz coordinates in this plane, ids and the neuronal class. Every neuronal profile passing through these planes in FANC or FAFB was individually reviewed, reconstructed and annotated by manual proofreading of the corresponding automated segmentations. We reviewed 3874 profiles (which received a total of 100,747 edits) in FANC, and 3693 profiles (which received 131,207 edits) in FAFB. Both datasets provide open community-based proofreading platforms (see https://flywire.ai/ and https://github.com/htem/FANC_auto_recon/wiki), and some of these edits were due to general proofreading in each volume, but the majority were from our comprehensive proofreading of neck connective neuron. The first pass review of the MANC neck connective was carried out in mid 2021; for FAFB the initial review periods were late 2020/early 2021 and again in mid 2022. After initial review of ANs and SAs in the VNC datasets, neurons were assigned a putative soma side programmatically, directly or indirectly via a MANC mirroring registration (H. S. J. *. Cheong et al., 2024). Neurons were mirrored based on their soma side or their neck plane side and NBLAST clustered (Costa et al., 2016). This analysis allowed for an initial grouping of left-right homologous sets and to identify neurons with different morphologies on each side of the nervous system, triggering further proofreading (since these differences usually resulted from residual segmentation errors). The combination of comprehensive proofreading of the whole dataset followed by within dataset matching and focussed proofreading was essential to ensure high quality connectome data and annotation. Most DNs and ANs have a unique morphology and were grouped into pairs, otherwise neurons were combined into larger groups containing more than one neuron per side. This was especially the case for SA neurons in FAFB. A similar approach has recently been described for MANC (H. S. J. *. Cheong et al., 2024). Note that proofreading across the FlyWire-FAFB dataset was reported in aggregate in (Dorkenwald et al., 2023) and that a first version of the neck connective annotations was released as part of the brainwide FlyWire annotations paper (Schlegel et al., 2023).

### Light microscopy identification

DNs from LM images were identified by overlaying the EM reconstructed DNs with images of Gal4 lines, mainly from the Namiki collection ((Namiki et al., 2018) and in preparation Namiki et al. 2024), Janelia’s Gal4 and Split-Gal4 collections (Jenett et al., 2012; Meissner et al., 2024; Tirian & Dickson, 2017), or via the neuronbridge tool (Clements et al., 2022; Meissner et al., 2023) for MANC DNs. To compare the reconstructions and LM images in the same space, the latter were segmented and transformed into MANC space as described in (H. S. J. *. Cheong et al., 2024) or into FAFB space. The full list of DN types with the identifier for the LM image (slide_code) and for the type (VFB_ID) can be found in supplemental file 2 - DN_identification. A small list of ANs were also matched to LM in FAFB and MANC as they were of special interest for the circuit described in Fig. 3 (see supplemental file 2 - AN_indentification). We did not match ANs to LM images systematically, due to a lack of a catalogue describing these neurons (as is available for DNs; (Namiki et al., 2018)). SA neurons were divided into subclasses by comparing them to LM images of Janelia’s Gal4 and Split- Gal4 datasets using the neuronbridge tool (Clements et al., 2022; Meissner et al., 2023) for MANC and then manually matching their axonal continuations into the brain to FAFB neuron reconstructions. Extended Data Fig. 2 shows the LM line that SA neurons were matched to, as well as the assigned long_tract and entry_nerve that were used to give SA neurons a subclass name,aiding their identification (see also supplemental file 2 - FAFB_SA_identification).

The process of matching to LM data is not exhaustive (in part because LM data is not yet available for all neurons) and we kindly ask the *Drosophila* community to contact the authors with missing identifications which can be reviewed and integrated in this resource.

### Matching of neurons across VNC datasets

FANC DNs and ANs were transformed into MANC space using the transform_fanc2manc function from the fancr R package (https://github.com/flyconnectome/fancr). This is a one step thin plate spline transform based on 2110 landmark pairs fitted to a complex transformation sequence mapping FANC to the JRCVNC2018F template (Phelps et al., 2021) and JRCVNC2018F to MANC (Takemura et al., 2023). A combination of NBLAST (Costa et al., 2016), and connectivity analysis was used to identify candidate morphological matches. These were assessed manually and assigned MANC names if the match was of high confidence (confidences ranged from 1 to 5, high is >3, Supplemental file 2). Additionally all ANs that were not matched with high confidence were assigned hemilineage and soma location in FANC and were compared by two independent annotators within each hemilineage after thorough review of the non-matching ANs to exclude reconstruction problems as a cause. ANs were first matched between FANC and MANC as individual neurons. We then reviewed these MANC- FANC matches to ensure that they respected the groups of neurons previously defined in MANC, thus providing an additional layer of validation. Cosine similarity as well as the identity of strong upstream and downstream synapses partners was used to help resolve ambiguous cases.

### Tract identification

VNC longitudinal tracts for MANC ANs, MANC SAs and FANC DNs were identified as previously described in (H. S. J. *. Cheong et al., 2024). In brief, neurons were simplified to their longest neurite starting from the VNC entry point at the neck and subsequently NBLAST clustered (Costa et al., 2016). The clusters were manually assigned a tract by overlaying with tract meshes made for MANC (H. S. J. *. Cheong et al., 2024).

Analysis of AN tracts revealed for the first time that one cluster did not match any of the previously published DN tracts. This new tract was given the name **A**N-specific **d**orsal **m**edial tract (ADM) in accordance with the tract naming of (Court et al., 2020).

### Neuropil identification

Primary brain neuropils were assigned in the FAFB dataset using the per neuron neuropil counts of presynapses for ANs and the postsynapses for DNs in the 783 FAFB version (available for download at https://codex.flywire.ai/api/download). A single brain neuropil was assigned if 80% of all synapses were within that neuropil, two neuropils were assigned as a name (primaryneuropil_secondaryneuropil) if combined they reached the 80% threshold and each contained at least 5%. An assignment as *multi* was given to 367 DNs and 282 ANs as they collected input or gave significant output (>20%) to more than two neuropils.

Primary VNC neuropils were assigned to all DNs and ANs in the MANC/FANC datasets as previously performed for MANC DNs (H. S. J. *. Cheong et al., 2024). For MANC AN synapses, we used the neuprint synapse ROI information of the manc:v1.2. For FANC AN and DN synapses we retrieved the synapses allocated to AN and DN IDs from the synapse parquet file, retrievable via FANC CAVE and available from the FANC community upon request (provided by Stephan Gerhard).

A single neuropil abbreviation was given to a DN/AN if they innervated a VNC neuropil with >80% of their pre/post synapses. The two letter abbreviations nt, wt, hl, it, lt, fl, ml, hl, mv, ov, ad correspond to NTct, WTct, HTct, IntTct, LTct, LegNpT1, LegNpT2, LegNpT3, mVAC, Ov and ANm, respectively. Additionally DNs/ANs which innervated a combination of upper tectulum (ut) or leg neuropils (xl) with more than 80% of their pre/post synapses were given those abbreviations accordingly. Any neuron that did not fall into one of those two categories was grouped as xn, standing for multiple neuropils. ANs that only contained a soma and soma tract in the VNC were excluded from this neuropil analysis and referred to as XA as previously described (Marin et al., 2023).

If the neuropil names were inconsistent within a group or pair of neurons, we calculated the mean of the pre- or post synapses to determine the assignment.

### Information flow ranking

The information flow ranking previously reported by Dorkenwald et al. (2023) for FlyWire, was subsetted for descending neurons and averaged by DN type. The information flow analysis is based on an algorithm implemented in Schlegel et al. (2021) (https://github.com/navis-org/navis). A low rank indicates a more direct connection from sensory inputs to that DN type.

### Sexually dimorphic and sex-specific neurons

DNs previously described to be sex-specific such as the female specific oviDNs or male specific pIP1 were matched to the available light level data and referred to as **sex-specific** throughout the paper. Other DNs and ANs that we could not match between the VNC datasets (between female and male), couldn’t be matched to light level data, but were well reconstructed and had a left-right partner were considered to be potentially female/male specific, also referred to in text and figures as **sex-specific**.

DNs such as DNa08 that exist in both sexes but are known to be dimorphic in morphology were matched to light level data and referred to as **sexually dimorphic** (**sex. dimorphic** in figure). Other DNs and ANs that we could confidently match across the two VNC datasets but that were dimorphic in morphology are also referred to as **sexually dimorphic** (**sex. dimorphic** in figure). The following neurons were not considered even though they show morphological differences:

● Specifically, neurons presumed to be neuropeptidergic were not included, as big morphological differences in neuronal arbour are common even between left and right of the same animal (Marin et al., 2023).
● The ascending histaminergic neurons (AHNs), which have been shown to have a difference in morphology not related to the sex of the animal (H. S. J. Cheong et al., 2024).
● Those neurons that innervate the abdominal ganglion, where there are problems in the FANC dataset that make it impossible to distinguish between a difference in reconstruction state and potential dimorphism, noted as reconstruction issues in the supplementary tables.
● Differences in number of ANs or DNs of a type were not considered as dimorphism in this paper as they occurred in neurons that we consider populations and whose numbers differed across the two sides of one animal. A difference in number was noted in the supplementary tables as biological variation, a match that is not ascending, or a general matching problem, if one side was not found (Supplemental file 2).

### Synapse density plots

To calculate the synapse density of sexually dimorphic and sex-specific neurons in the VNC we collected all synapses of the identified neurons in each dataset (FAFB: cleft-score > 50 applied). We then tiled the space their synapses occupy into roughly isotropic voxels of 5 µm size and counted synapses in each voxel. Synapses were then colour coded by density and plotted in three-dimensional space.

### FANC neuron types

All FANC neurons that can be matched to MANC neurons are referred to by their MANC name, with an additional “f” denoting female, when presented in comparative graphs or connectivity plots. All FANC neurons identified are listed in the supplementary material (Supplemental file 2 - FANC_DNs, FANC_ANs, other_MANC_FANC_matches).

FANC ANs and DNs that were not previously identified in LM and that could not be matched between datasets were assigned new type names. For ANs and DNs, the type names were given in accordance with the previously established systematic type names (DN-target neuropil abbreviation-number or AN-hemilineage abbreviation-number). To distinguish from the previous type names in MANC, the numbering starts at 999 and goes down.

### Connectivity

For connectivity graphs we used a threshold of weight >10 and percent output >0.5% for the initial retrieval of partners of the neurons of interest. In the following step we added all MNs, SNs or SAs that connected to those with a weight >5 to adjust for known reconstruction problems in these neurons and for the fact that sensory neurons tend to make fewer synapses with their partners individually and connect as a population of the same sensory origin (reflected by their type). Once all neurons of interest had been defined we took an all-by-all connectivity adjacency matrix, in which all values were converted to input percent to the receiving neuron, averaged by type (unless otherwise indicated). The graphs shown in the figures note the additional percent thresholds that were chosen for the nodes plotted in each graph.

### Neuroglancer Resource

To help compare the neurons described in our work we created a neuroglancer environment (Maitin-Shepard et al., 2021) displaying meshes for all three datasets in a common space. This environment can be opened in any modern web browser (we use Google Chrome) by following the short URL https://tinyurl.com/NeckConnective.

We opted to use the Janelia FlyEM male CNS dataset as a single anatomically consistent target space for display based on resources provided by (Nern et al., 2024). FlyWire neurons were transformed into the space of the male CNS brain using rigid and non rigid consecutive registrations (Bates et al., 2020; Nern et al., 2024). Meshes for MANC neurons are those released by (Takemura et al., 2023); we then use neuroglancer to apply an affine registration “on-the-fly” to place them within the space of the VNC of the male CNS volume. We applied non-rigid transformations (fancr::transform_fanc2manc function described above) to put FANC neurons into MANC space and then used the same MANC to male CNS affine registration within neuroglancer to complete the transformation into male CNS space. Metadata annotations are provided for the three datasets using the format Type_Side_Class format. At present only the optic lobe portion of the male CNS EM volume has been released but having all the data transformed into male CNS space means that this neuroglancer scene can be modified with minimal effort to display the full male dataset when it becomes available.

## Supporting information

High resolution images of the figures

## Acknowledgments

We thank the following people for their contributions to this work: our colleagues in the FANC and FlyWire communities for additional contributions to the proofreading of neurons that we report as detailed in Supplementary file 3. This sheet also notes proofreaders employed by Ariadne.ai and operating on our behalf. Elizabeth C. Marin for helping us identify hemilineage and soma neuromere for sex-specific FANC ANs. Stefanie Hampel and Andrew M. Seeds for identification of aDN1 and aDN2. Gerry Rubin for sharing unpublished lines for the identification of DNp68 and DNg107. Jens Goldammer for sharing unpublished lines for the identification of DNp60, DNp66, DNp72, DNg95 and DNg100. Alice Schildkamp for final proofreading of problematic and dimorphic ANs. The FlyLight Project Team at Janelia Research Campus for the use of the Gal4 and Split-Gal4 collections. Helen Langley, Markus Pleijzier and Elizabeth C. Marin for comments on the manuscript.

H. S. Seung declares a financial interest in Zetta AI. L. Serratosa Capdevila declares a financial interest in Aelysia Ltd.

This work was supported by Wellcome Trust collaborative awards 220343/Z/20/Z and 221300/Z/20/Z to GSXEJ and GMC, GM Rubin, S Waddell and M Landgraf; core support from the MRC (MC-U105188491) to GSXEJ; a Neuronex2 award to GSXEJ (NSF 2014862, MRC MC_EX_MR/T046279/1); NIH Brain Initiative 1RF1MH120679-01 to D Bock with GSXEJ; a Boehringer Ingelheim Fonds and the Cambridge Commonwealth, European and International Trust to I.R.B.; a Walter-Benjamin-Fellowship from Deutsche Forschungsgemeinschaft to TS (STU 793/2-1).

## Author contributions

TS, PB, LSC, BJM, AJ, SF, IM, AR, FK, LK, HSJC, JK, ET, RP, ASB, GMC, MC, GSXEJ and KE were involved in the proofreading. PS, JSP, MB, SD, AM, SY, CEM, ARS, SS, MM, JT, WAL, MC, GSXEJ and KE made core contributions to the datasets. TS, PB, LSC, MG, SC, IRB, IM, AR, PS, MC and KE performed matching across sides and across VNC datasets. TS, PB, SN, HSJC, JK, ET, RP, GMC, MC, GSXEJ and KE performed LM-EM matching. KE determined tract assignments and TS determined neuropil assignments. PB, BM, AR and KE identified entry nerve assignments in FANC. SC produced the neuroglancer environment. GSXEJ developed the fancr R package. TS, PB, ASC and KE performed analyses. TS, PB and KE produced the figures with input from coauthors. TS, PB, GSXEJ and KE wrote the manuscript with input from MC, SC, IRB, LSC. KE and MC coordinated proofreading, matching and LM identification. MC supervised LSC, BJM, AJ, SF, MG, IM, AR. KE supervised FK and LK. GSXEJ managed the overall effort and acquired funding.

## Supplemental information

### Supplementary file 1 Seed planes

Two tables listing the xyz coordinates, svid, root_id, side and class of profiles passing through the FAFB or FANC seed plane.

### Supplementary file 2 Typing and matching

13 tables listing the neuronal ids and annotations used in the manuscript for: FAFB_DNs, FANC_DNs, MANC_DNs, dimorphic_DNs, DN_identification, FAFB_ANs_SAs, FANC_ANs, FANC_SAs, MANC_ANs, dimorphic_ANs, AN_identification, FAFB_SA_identification, other_MANC_FANC_matching.

### Supplementary file 3 User edits

Two tables listing the number of edits to the neck connective neurons in FANC and FAFB summarised by lab.

---

**Extended Data Fig. 1.**
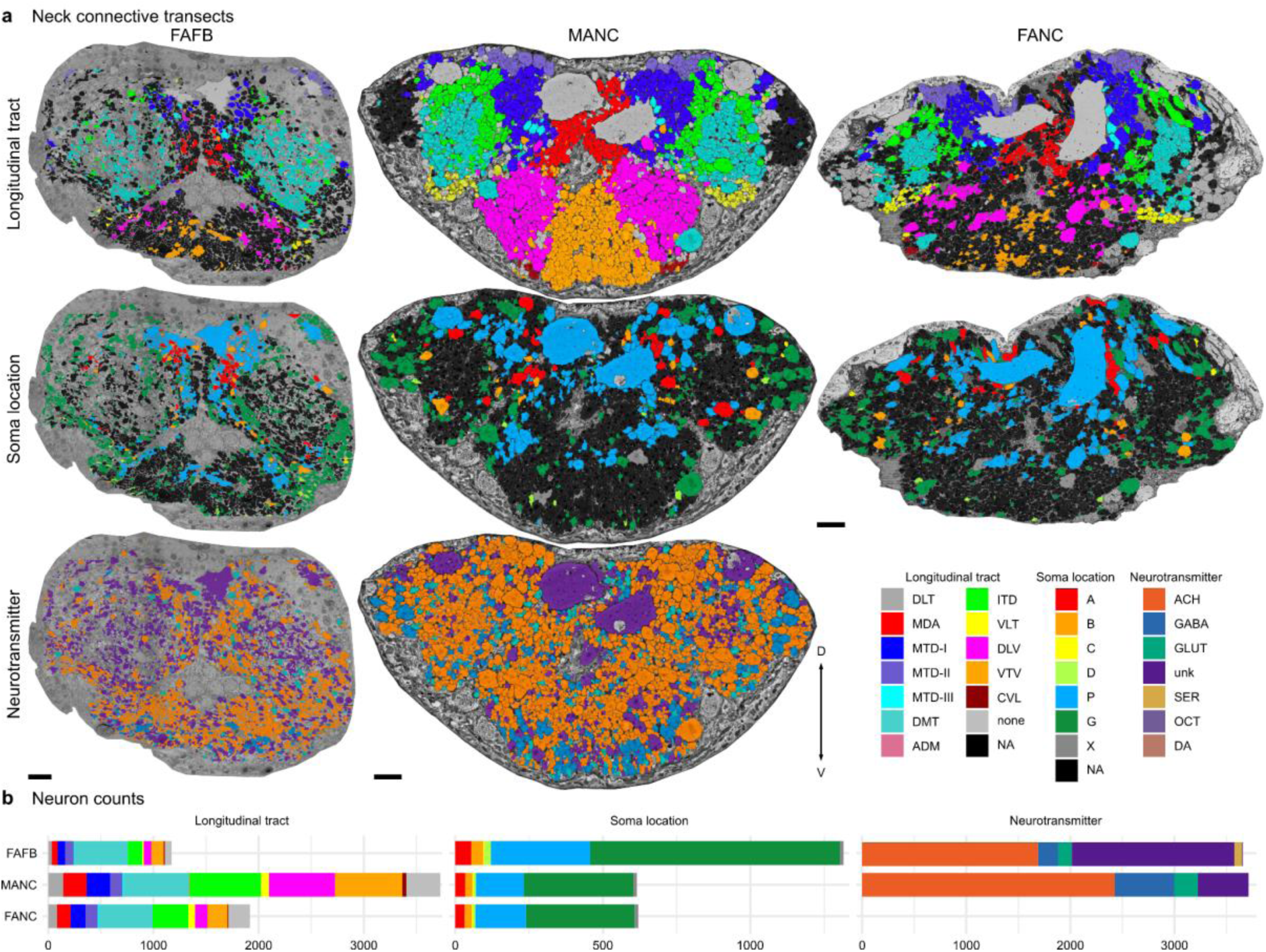
Cross section (frontal) of the neck connective in the three datasets. **a**, All neurons in the neck connective colour coded by their longitudinal tract, soma location or predicted neurotransmitter (Eckstein et al., 2020). **b**, Number of neurons by longitudinal tract, soma location or neurotransmitter in the three datasets. Neurotransmitter predictions are not yet available in FANC.

**Extended Data Fig. 2.**
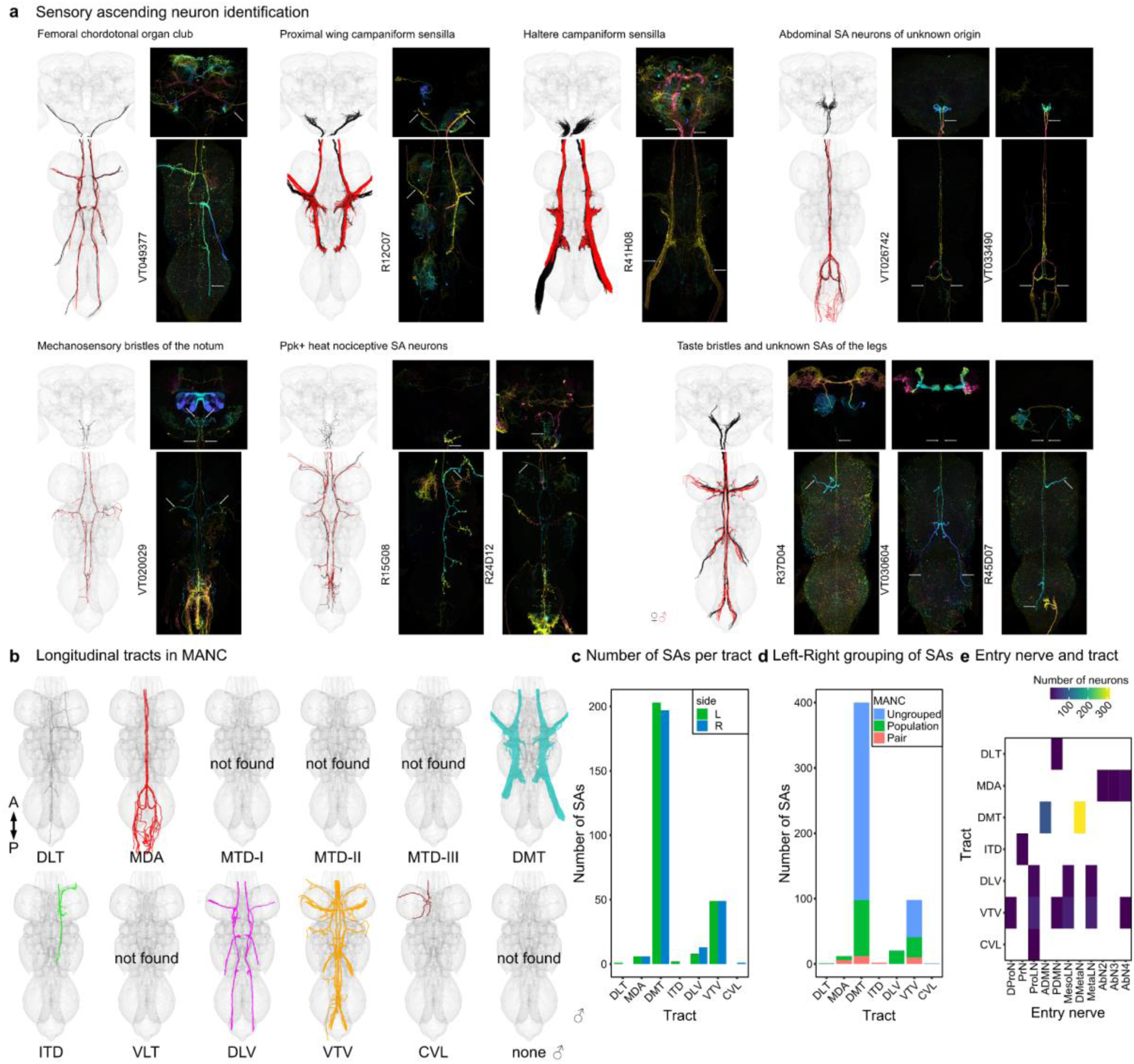
Sensory ascending neurons. **a**, Morphology of sensory ascending neurons identified in the three EM volumes. In black the EM morphology of DNs from female datasets (FAFB, FANC), in red from the male dataset (MANC). Next to them the LM images that allowed a grouping into sensory subclasses. **b**, Tract-based analysis of sensory ascending neurons in MANC. None of the SAs project along the MTD, or VLT tract. **c**, Number of SAs in each tract. **d**, Number of SA grouped into pairs or populations. **e**, Correlation of entry nerve to tract membership for MANC SAs (Marin et al. 2023).

**Extended Data Fig. 3.**
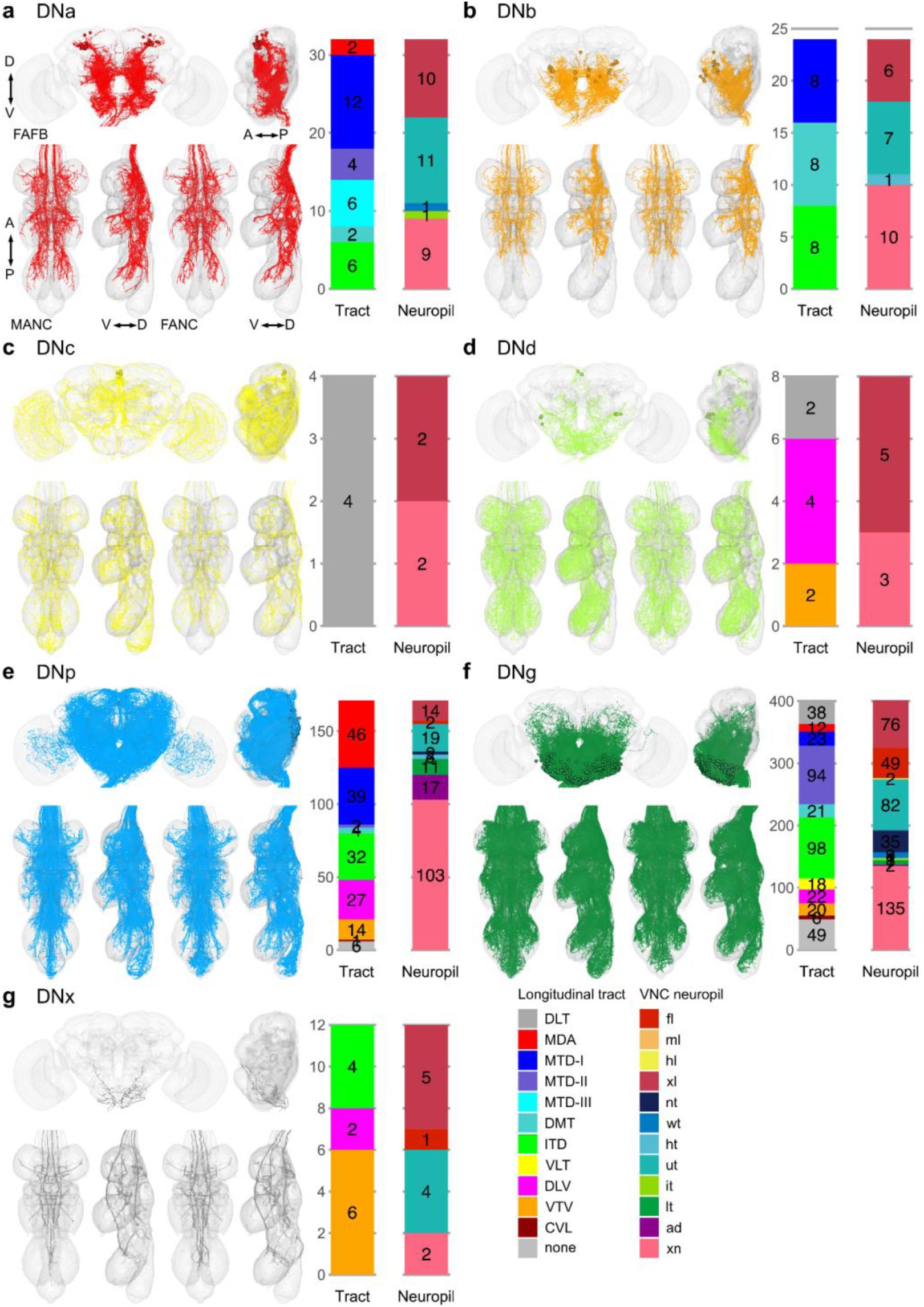
Morphology matched across the neck - soma. Morphology of LM matched DNs across the three datasets colour coded by cell body location according to (Namiki et al. 2018). **a,** DNa neurons have an anterior dorsal soma; **b**, DNb an anterior ventral soma; **c**, DNc a soma in the pars intercerebralis; **d**, DNd a soma in an anterior outside cell cluster; **e**, DNp are on the posterior surface; **f**, DNg are located in the GNG and **g**, DNx are outside the brain. In each panel the top images show reconstruction in FAFB in anterior and lateral view; the two bottom left images show MANC and two bottom right FANC in ventral and lateral view, respectively. The bar charts represent the distribution of the VNC characteristics longitudinal tract and neuropil innervation for the neurons in each category - see colour legend.

**Extended Data Fig. 4.**
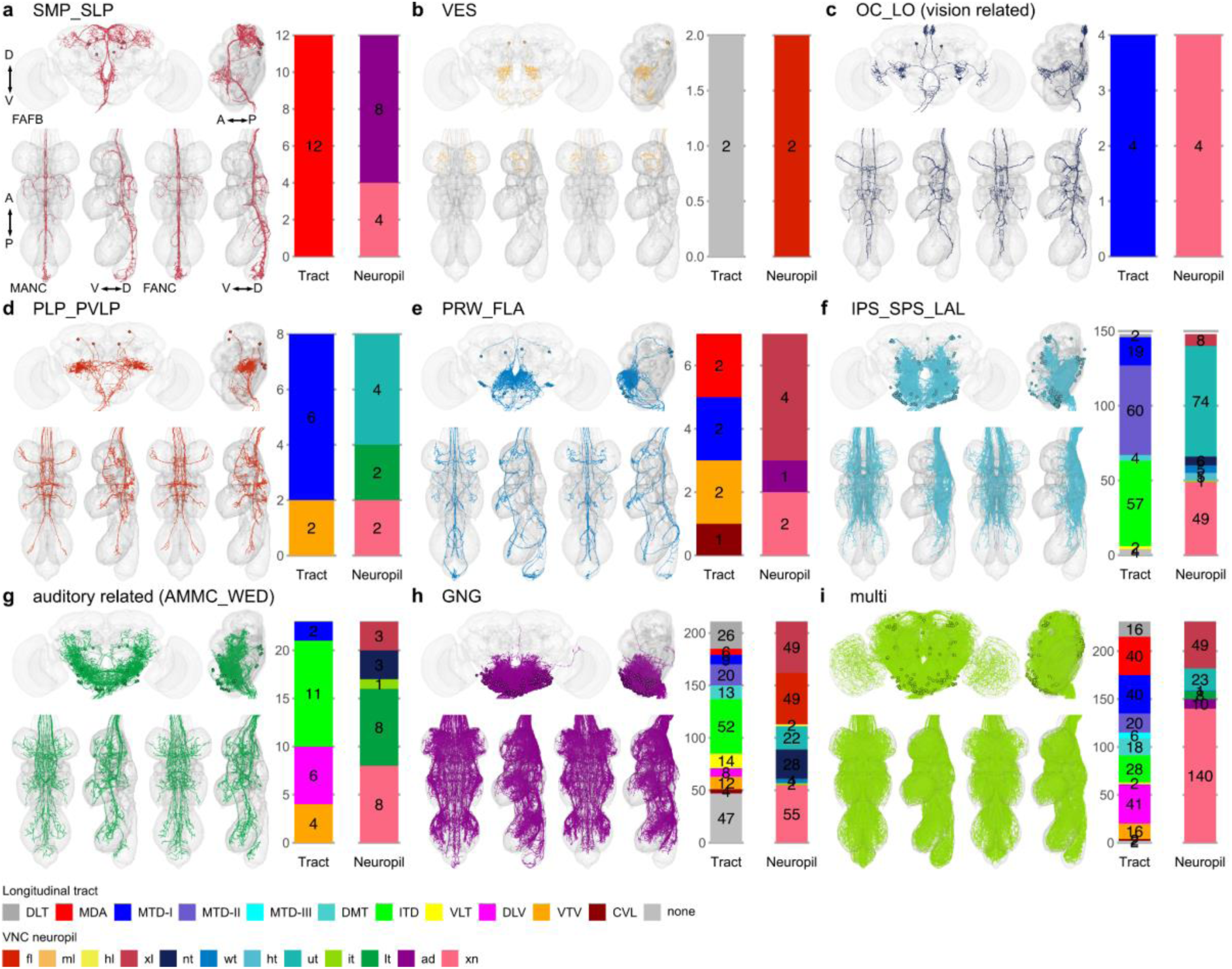
Morphology matched across the neck - brain neuropil. Morphology of LM matched DNs across the three datasets colour coded by their brain neuropil innervation. DNs with input neuropil. **a,** superior medial protocerebrum and superior lateral protocerebrum (SMP_SLP); **b**, vest (VES); **c**, ocellar ganglion and lobular (OC_LO, vision related); **d**, posterior lateral protocerebrum (PLP); **e**, prow and flange (PRW_FLA); **f**, posterior slope and lateral accessory lobe (IPS_SPS_LAL); **g**, antennal mechanosensory and motor centre and wedge (AMMC_WED, auditory related); **h**, gnathal ganglia (GNG) and **i**, multiple innervations of neuropils across the brain (multi). In each panel the top images show reconstruction in FAFB in anterior and lateral view; the two bottom left images show MANC and two bottom right FANC in ventral and lateral view, respectively. The bar charts represent the distribution of the VNC characteristics longitudinal tract and neuropil innervation for the neurons in each category - see colour legend.

**Extended Data Fig. 5.**
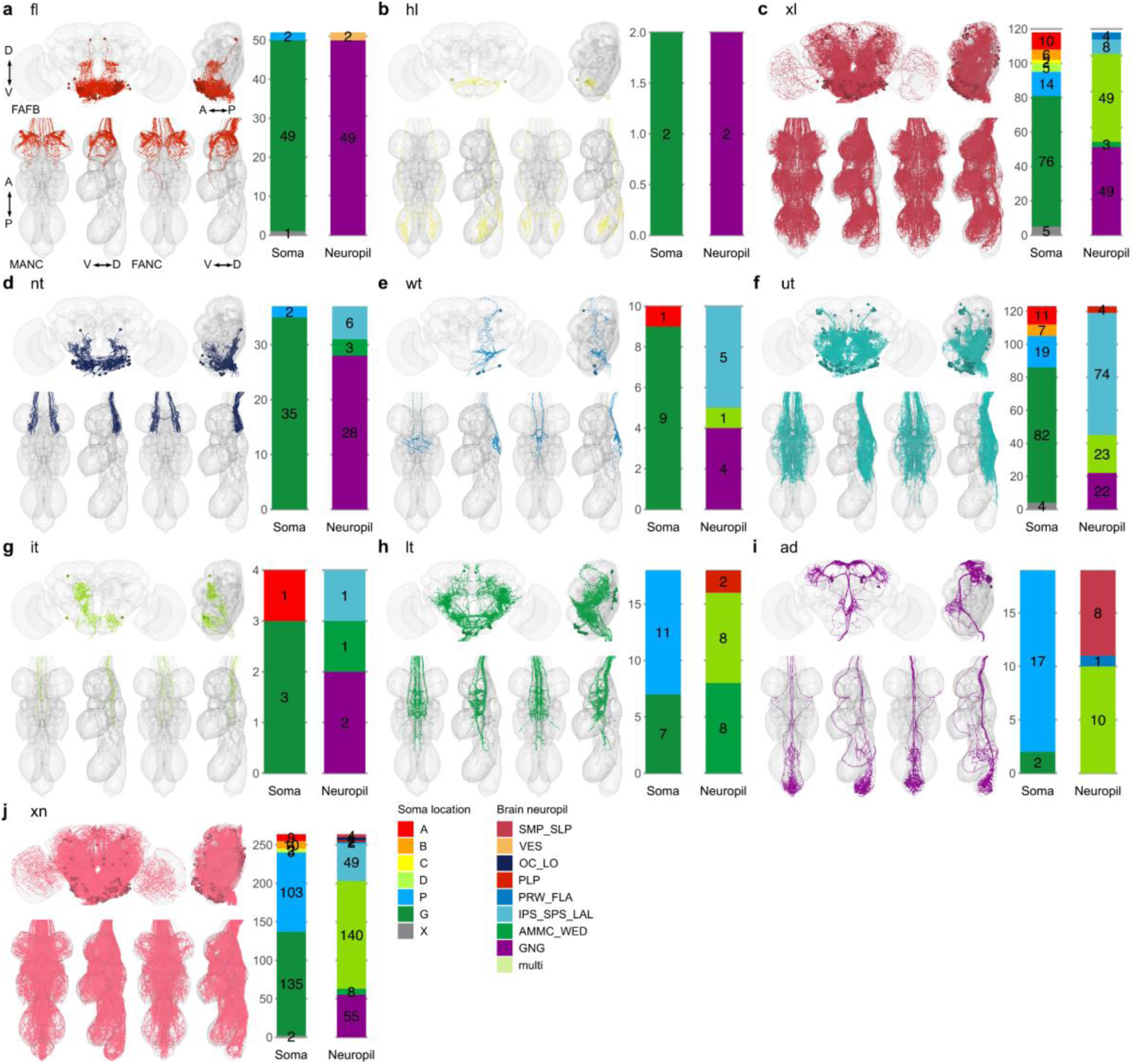
Morphology matched across the neck - VNC neuropil. Morphology of LM matched DNs across the three datasets colour coded by VNC neuropil innervation. DNs with output neuropil. **a,** front leg (fl); **b**, hind leg (hl); **c**, multiple innervation in leg compartments (xl); **d**, neck tectulum (nt); **e**, wing tectulum (wt); **f**, multiple innerveration into upper tectulum neuropils of the neck, wings and halteres (ut); **g**, intermediate tectulum (it); **h**, lower tectulum (lt); **i**, abdomen (ad) and **j**, multiple innervations of neuropils across the VNC (xn). In each panel the top images show reconstruction in FAFB in anterior and lateral view; the two bottom left images show MANC and two bottom right FANC in ventral and lateral view, respectively. The bar charts represent the distribution of the brain characteristics soma location and neuropil innervation for the neurons in each category - see colour legend.

**Extended Data Fig. 6.**
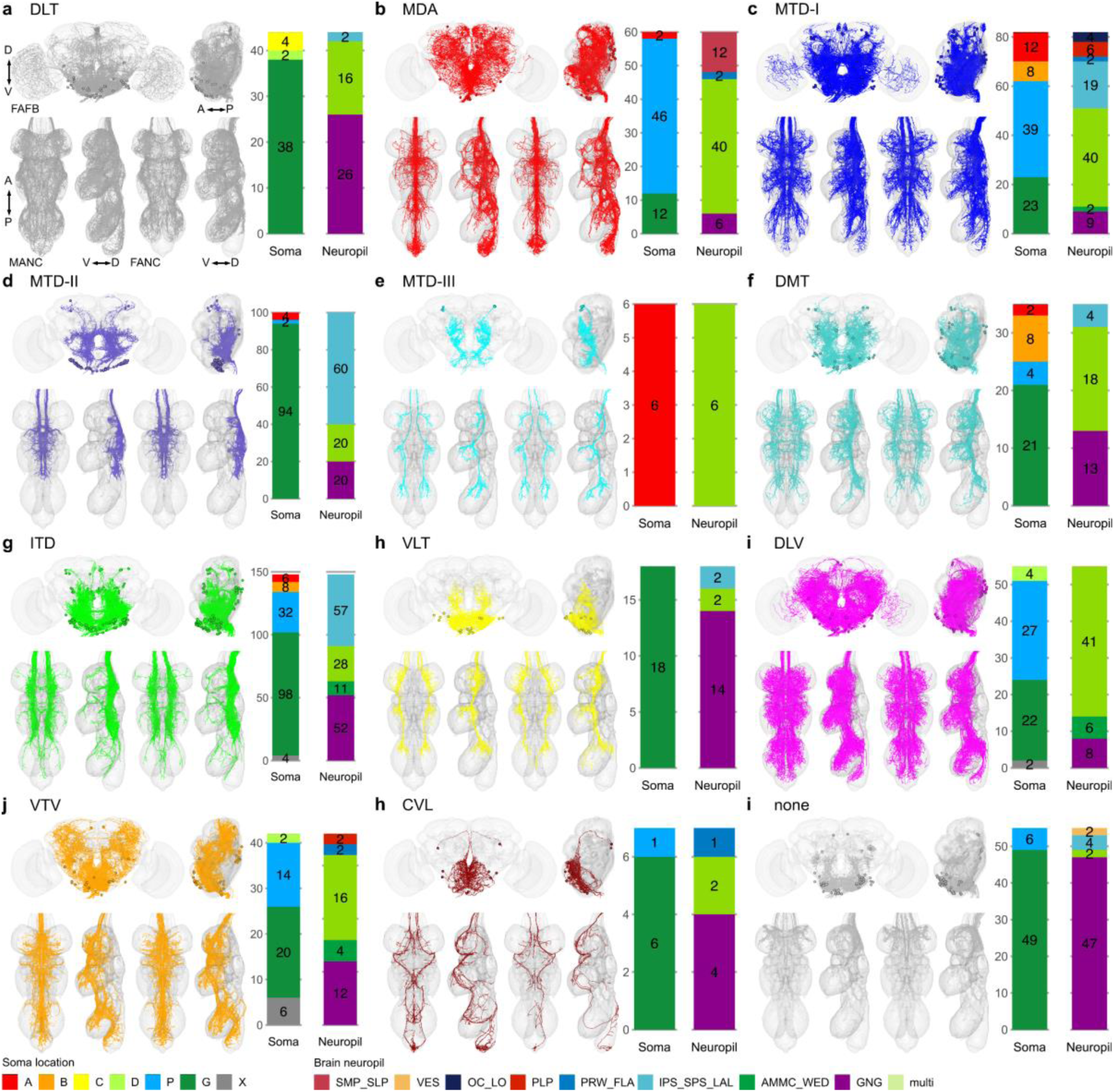
Morphology matched across the neck - tract. Morphology of LM matched DNs across the three datasets colour coded by longitudinal tract membership in the VNC. DNs in the tract. **a**, dorsal lateral tract (DLT); **b**, median dorsal abdominal tract (MDA); **c**, ventral route of the mediate tract of dorsal cervical fasciculus (MTD-I); **d**, dorsal route of the mediate tract of dorsal cervical fasciculus (MTD-II); **e**, lateral route of the mediate tract of dorsal cervical fasciculus (MTD-III); **f**, dorsal median tract (DMT); **g**, intermediate tract of dorsal cervical fasciculus (ITD); **h**, ventral lateral tract (VLT); **i**, dorsal lateral tract of ventral cervical fasciculus (DLV); **j**, ventral median tract of ventral cervical fasciculus (VTV); **h**, curved ventral lateral tract (CVL) and **i**, no tract membership (none). In each panel the top images show reconstruction in FAFB in anterior and lateral view; the two bottom left images show MANC and two bottom right FANC in ventral and lateral view, respectively. The bar charts represent the distribution of the brain characteristics soma location and neuropil innervation for the neurons in each category - see colour legend.

**Extended Data Fig. 7.**
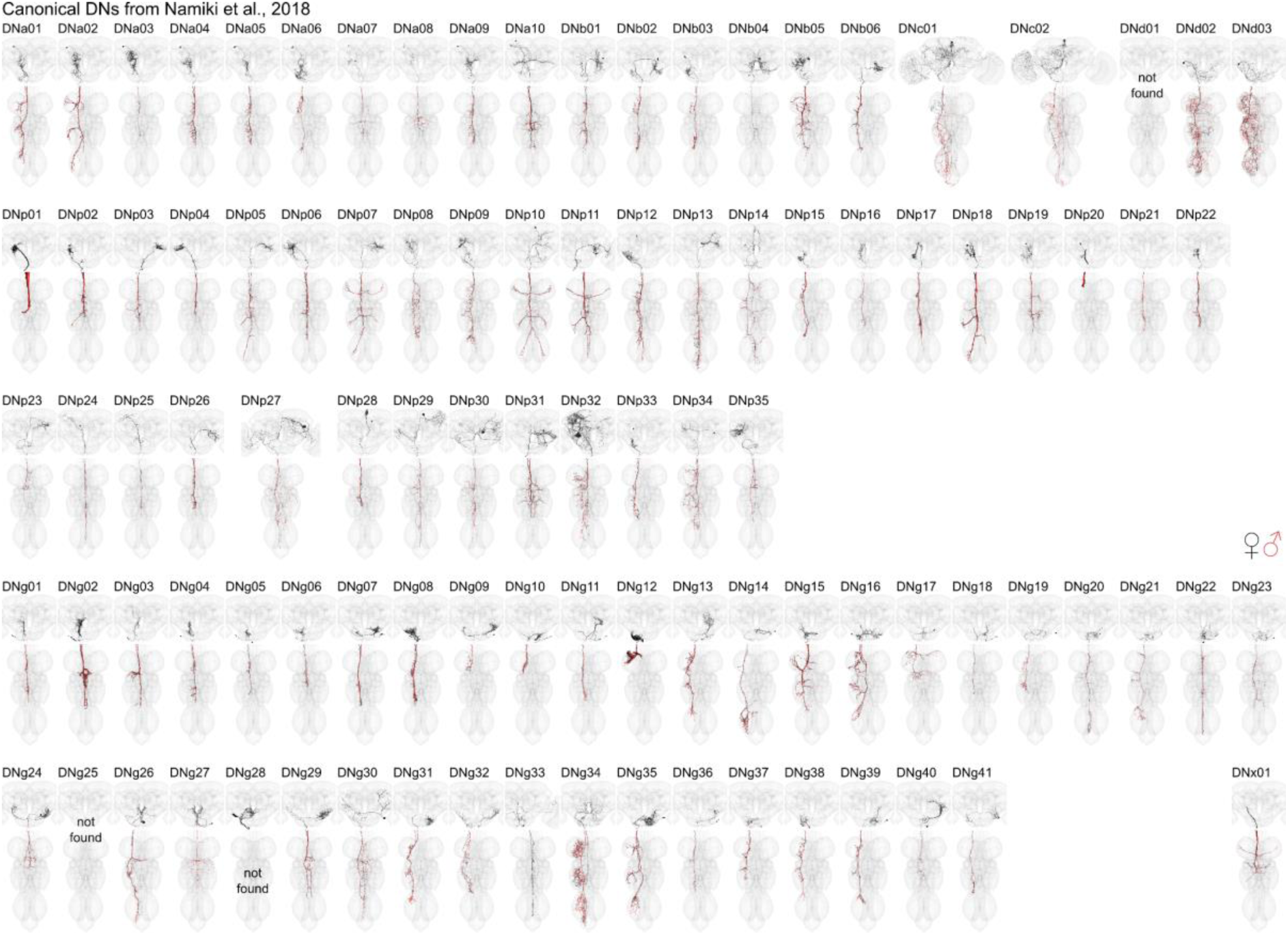
DN matching to Namiki et al. 2018. Morphology of identified DNs across all three datasets with nomenclature as described in (Namiki et al. 2018). Two DN types could not be identified (DNd01, DNg25) in any of the three EM datasets and one DN type (DNg28) is only identifiable in the brain. See supplementary table DN_identification for slide codes and for DN synonyms from the literature. In black the morphology of DNs from the female datasets (FAFB, FANC) in red from the male dataset (MANC). This figure is also provided in high resolution and DNs can be viewed in 3D at https://tinyurl.com/NeckConnective.

**Extended Data Fig. 8.**
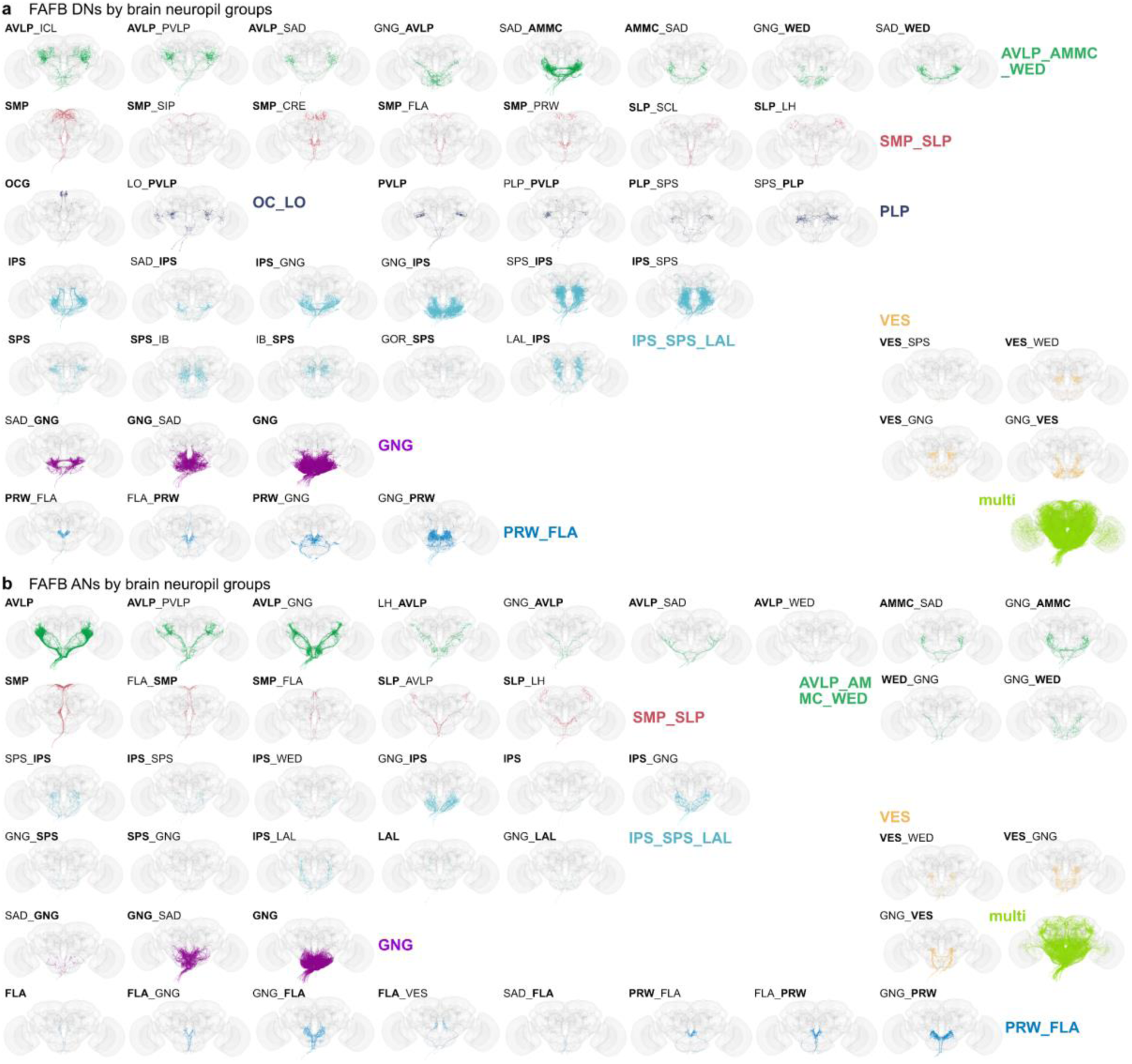
Brain neuropil groups. Morphology of DNs and ANs by primary brain neuropil. **a**, All DNs in FAFB were assigned one or two input neuropils. DNs that receive input from more than two neuropils are referred to as multi. **b**, All ANs in FAFB were assigned one or two output neuropils. ANs that output to more than two neuropils are referred to as multi. Morphologies are coloured by broader neuropil groups: auditory related neuropils (AVLP_AMMC_WED), higher order multimodal sensory integration 1 (SMP_SLP), vision related (OC_LO), multimodal sensory integration 2 (PVLP, PLP), sensory modalities from the GNG (GNG, SAD), multimodal sensory integration and steering (IPS_SPS_LAL), vest related (VES) and flange/prow (PRW_FLA).

**Extended Data Fig. 9.**
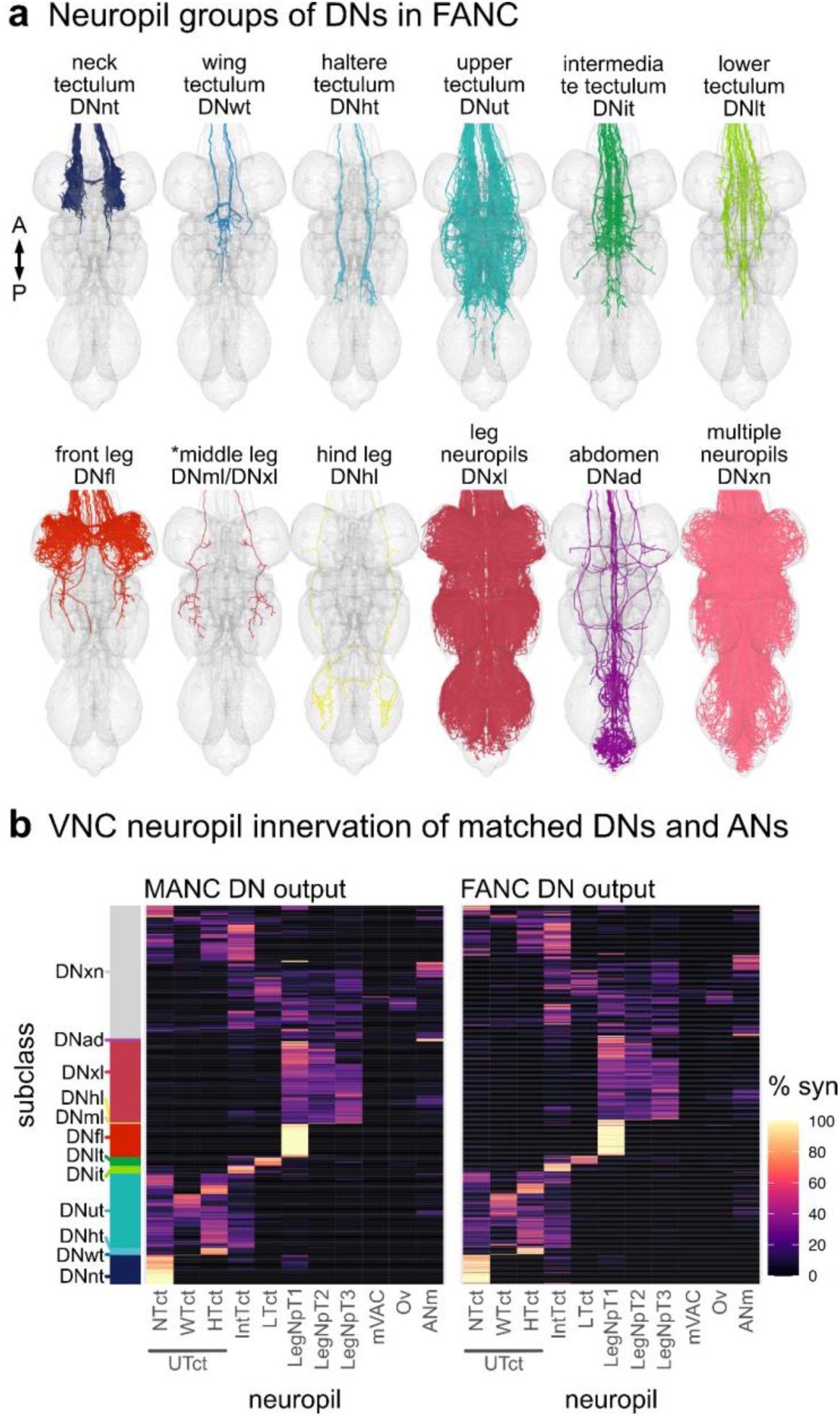
Neuropil-based analysis of descending neurons in FANC. **a,** Primary neuropil assignment of DNs in the FANC dataset to compare to previously published one in the MANC dataset (H. S. J. *. Cheong et al., 2024). **b**, Synaptic output in % by VNC neuropil of matched DNs in MANC and FANC. Each row represents one DN type, order is conserved between the two datasets. Left bar indicates the previously assigned neuropil based subclasses from the MANC dataset (H. S. J. *. Cheong et al., 2024).

**Extended Data Fig. 10.**
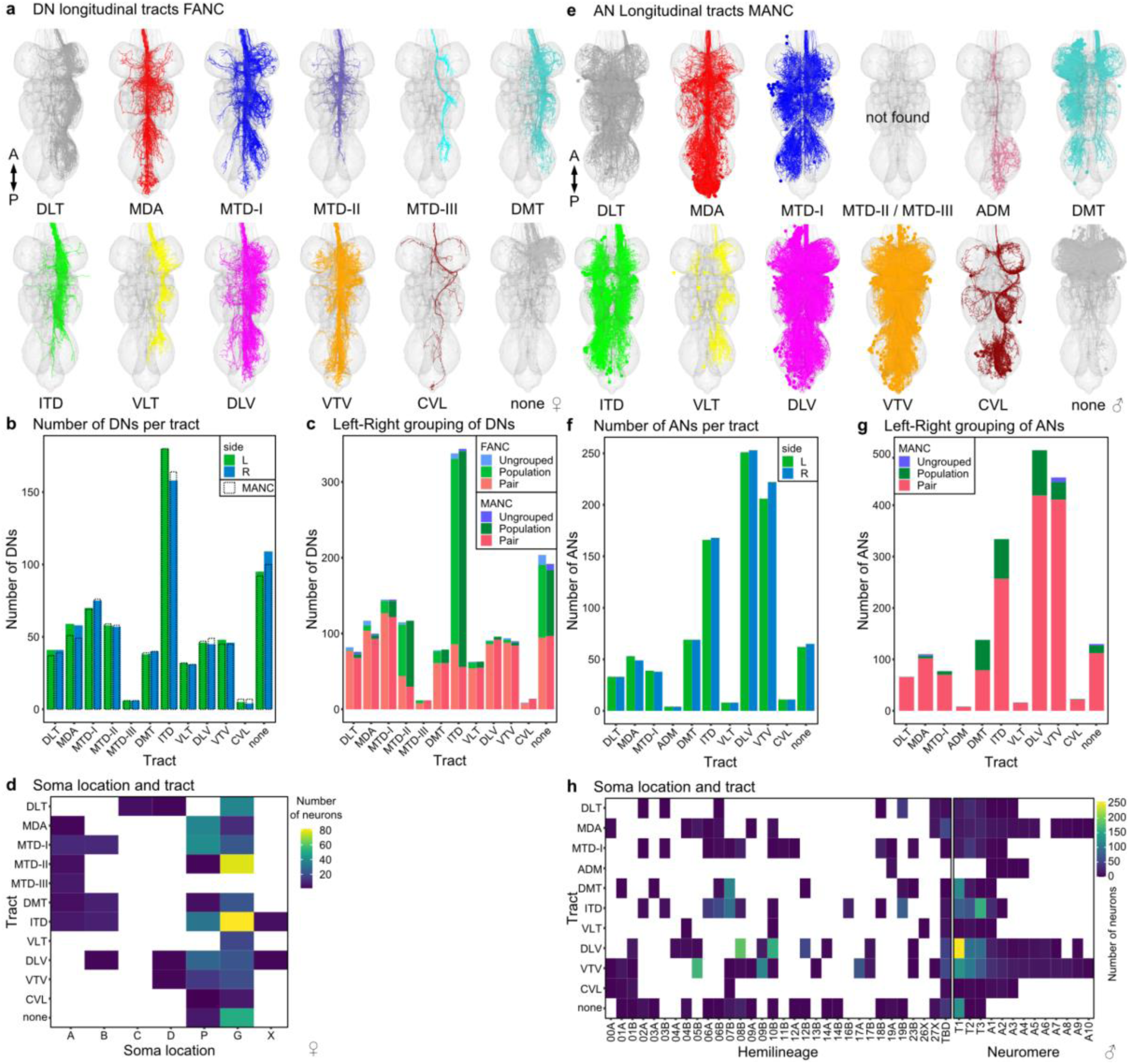
Tract-based analysis of descending neurons in FANC and ascending neurons in MANC. **a,** Tract assignment of all left side DNs in the FANC dataset to compare to previously published tract assignment in the MANC dataset (H. S. J. *. Cheong et al., 2024). **b**, Number of DNs for each tract in comparison to MANC DNs (dotted line). **c**, DNs grouped into pairs or populations comparing FANC to MANC. **d**, Correlation of soma location and tract membership for identified FANC DN types based on LM data (Namiki et al. 2018). **e**, Tract assignment of all left side ANs in the MANC dataset. None of the ANs project along the MTD-II or MTD-III tract. A small additional tract was observed for ANs, referred to as AN-specific dorsal medial tract (ADM). **f**, Number of ANs in each tract. **g**, ANs grouped into pairs or populations comparing MANC to FANC. **h**, Correlation of hemilineage and neuromere to tract membership for MANC ANs (Marin et al., 2023).

**Extended Data Fig. 11.**
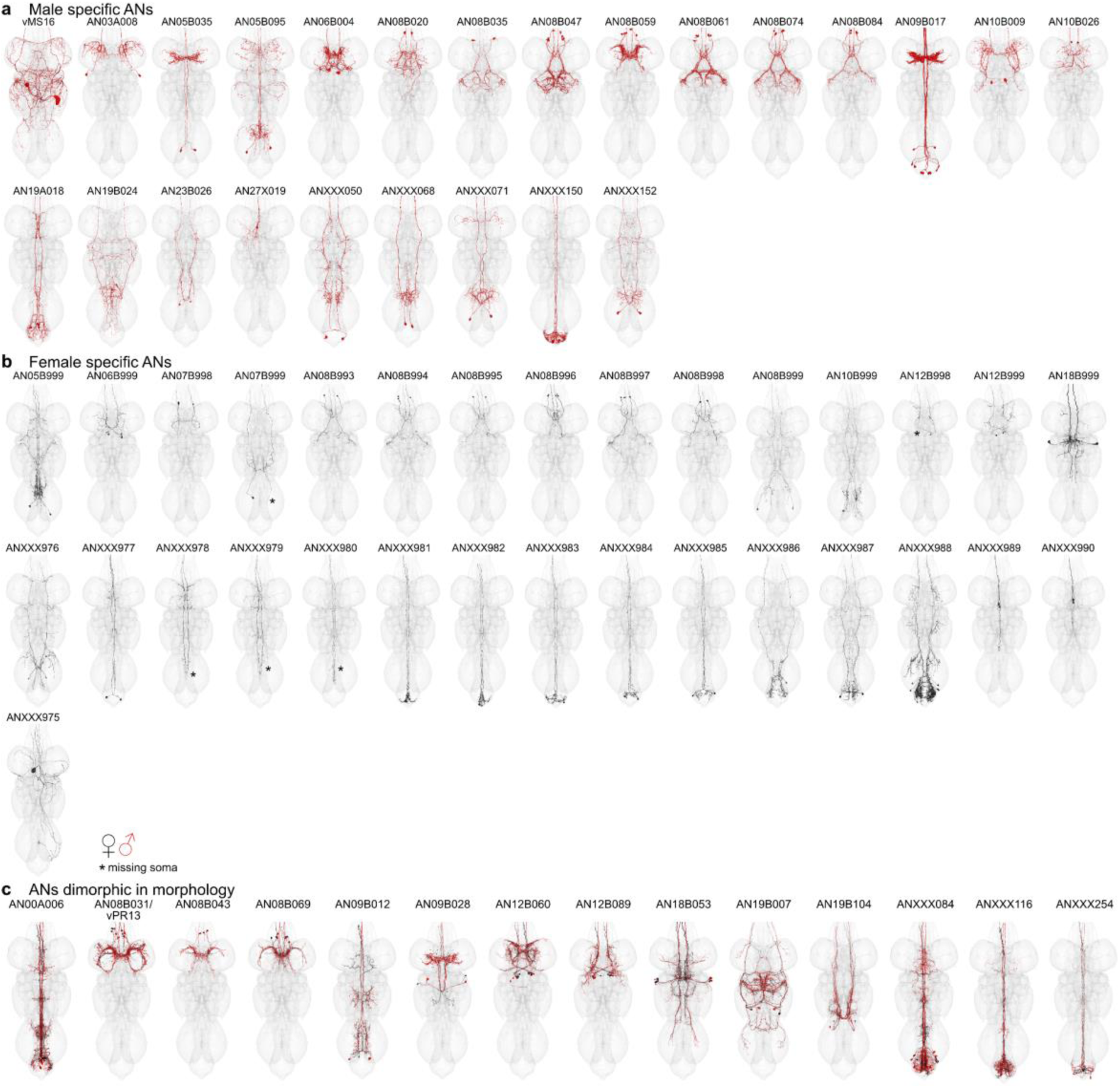
Potentially sexually dimorphic or sex-specific ANs in the VNC. **a**, Morphology of all the potentially male specific ANs by type. **b**, Morphology of the potentially female specific ANs by newly assigned types. **c**, Morphology of the potentially sexually dimorphic ANs by MANC type names. In black the EM morphology from the female dataset (FANC) in red from the male dataset (MANC). Stars indicate ANs with missing soma in FANC due to missing EM image data.

## References

1. Ache, J. M., Namiki, S., Lee, A., Branson, K., & Card, G. M. (2019). State-dependent decoupling of sensory and motor circuits underlies behavioral flexibility in Drosophila. Nature Neuroscience, 22(7), 1132–1139.

2. Auer, T. O., & Benton, R. (2016). Sexual circuitry in Drosophila. Current Opinion in Neurobiology, 38, 18–26.

3. Aymanns, F., Chen, C.-L., & Ramdya, P. (2022). Descending neuron population dynamics during odor-evoked and spontaneous limb-dependent behaviors. eLife, 11. 10.7554/eLife.81527

4. Azevedo, A., Lesser, E., Mark, B., Phelps, J., Elabbady, L., Kuroda, S., Sustar, A., Moussa, A., Kandelwal, A., Dallmann, C. J., Agrawal, S., Lee, S.-Y. J., Pratt, B., Cook, A., Skutt-Kakaria, K., Gerhard, S., Lu, R., Kemnitz, N., Lee, K., … Tuthill, J. C. (2022). Tools for comprehensive reconstruction and analysis of Drosophila motor circuits. In bioRxiv. 10.1101/2022.12.15.520299

5. Bates, A. S., Manton, J. D., Jagannathan, S. R., Costa, M., Schlegel, P., Rohlfing, T., & Jefferis, G. S. (2020). The natverse, a versatile toolbox for combining and analysing neuroanatomical data. eLife, 9. 10.7554/eLife.53350

6. Bidaye, S. S., Machacek, C., Wu, Y., & Dickson, B. J. (2014). Neuronal control of Drosophila walking direction. Science, 344(6179), 97–101.

7. Braun, J., Hurtak, F., Wang-Chen, S., & Ramdya, P. (2024). Descending networks transform command signals into population motor control. Nature. 10.1038/s41586-024-07523-9

8. Cachero, S., Ostrovsky, A. D., Yu, J. Y., Dickson, B. J., & Jefferis, G. S. X. E. (2010). Sexual dimorphism in the fly brain. Current Biology: CB, 20(18), 1589–1601.

9. Cande, J., Namiki, S., Qiu, J., Korff, W., Card, G. M., Shaevitz, J. W., Stern, D. L., & Berman, G. J. (2018). Optogenetic dissection of descending behavioral control in Drosophila. eLife, 7. 10.7554/eLife.34275

10. Chen, C.-L., Aymanns, F., Minegishi, R., Matsuda, V. D. V., Talabot, N., Günel, S., Dickson, B. J., & Ramdya, P. (2023). Ascending neurons convey behavioral state to integrative sensory and action selection brain regions. Nature Neuroscience, 26(4), 682–695.

11. Cheong, H. S. J., Boone, K. N., Bennett, M. M., Salman, F., Ralston, J. D., Hatch, K., Allen, R. F., Phelps, A. M., Cook, A. P., Phelps, J. S., Erginkaya, M., Lee, W.-C. A., Card, G. M., Daly, K. C., & Dacks, A. M. (2024). Organization of an ascending circuit that conveys flight motor state in Drosophila. Current Biology: CB, 34(5), 1059–1075.e5.

12. Cheong, H. S. J. *., Eichler, K. *., Stuerner, T. *., Asinof, S., Champion, A. S., Marin, E. C., Oram, T. B., Namiki, S., Siwanowicz, I., Costa, M., Berg, S., Janelia FlyEM Project Team, Jefferis, G. S. X. E., & Card, G. M. (2024). Transforming descending input into behavior: The organization of premotor circuits in the Drosophila Male Adult Nerve Cord connectome. eLife. 10.7554/eLife.96084.1

13. Clements, J., Goina, C., Hubbard, P. M., Kawase, T., Olbris, D. J., Otsuna, H., Svirskas, R., & Rokicki, K. (2022). NeuronBridge: an intuitive web application for neuronal morphology search across large data sets. In bioRxiv (p. 2022.07.20.500311). 10.1101/2022.07.20.500311

14. Clowney, J. E., Iguchi, S., Bussell, J. J., Scheer, E., & Ruta, V. (2015). Multimodal Chemosensory Circuits Controlling Male Courtship in Drosophila. Neuron, 87(5), 1036– 1049.

15. Cook, S. J., Jarrell, T. A., Brittin, C. A., Wang, Y., Bloniarz, A. E., Yakovlev, M. A., Nguyen, K. C. Q., Tang, L. T.-H., Bayer, E. A., Duerr, J. S., Bülow, H. E., Hobert, O., Hall, D. H., & Emmons, S. W. (2019). Whole-animal connectomes of both Caenorhabditis elegans sexes. Nature, 571(7763), 63–71.

16. Costa, M., Manton, J. D., Ostrovsky, A. D., Prohaska, S., & Jefferis, G. S. X. E. (2016). NBLAST: Rapid, Sensitive Comparison of Neuronal Structure and Construction of Neuron Family Databases. Neuron, 91(2), 293–311.

17. Court, R., Namiki, S., Armstrong, J. D., Börner, J., Card, G., Costa, M., Dickinson, M., Duch, C., Korff, W., Mann, R., Merritt, D., Murphey, R. K., Seeds, A. M., Shirangi, T., Simpson, J. H., Truman, J. W., Tuthill, J. C., Williams, D. W., & Shepherd, D. (2020). A Systematic Nomenclature for the Drosophila Ventral Nerve Cord. Neuron, 107(6), 1071–1079.e2.

18. Currier, T. A., & Nagel, K. I. (2020). Multisensory control of navigation in the fruit fly. Current Opinion in Neurobiology, 64, 10–16.

19. Dallmann, C. J., Luo, Y., Agrawal, S., Chou, G. M., Cook, A., Brunton, B. W., & Tuthill, J. C. (2024). Presynaptic inhibition selectively suppresses leg proprioception in behaving. bioRxiv : The Preprint Server for Biology. 10.1101/2023.10.20.563322

20. Dorkenwald, S., Matsliah, A., Sterling, A. R., Schlegel, P., Yu, S.-C., McKellar, C. E., Lin, A., Costa, M., Eichler, K., Yin, Y., Silversmith, W., Schneider-Mizell, C., Jordan, C. S., Brittain, D., Halageri, A., Kuehner, K., Ogedengbe, O., Morey, R., Gager, J., … FlyWire Consortium. (2023). Neuronal wiring diagram of an adult brain. bioRxiv : The Preprint Server for Biology. 10.1101/2023.06.27.546656

21. Eckstein, N., Bates, A. S., Du, M., Hartenstein, V., Jefferis, G. S. X., & Funke, J. (2020). Neurotransmitter Classification from Electron Microscopy Images at Synaptic Sites in Drosophila. In bioRxiv (p. 2020.06.12.148775). 10.1101/2020.06.12.148775

22. Eckstein, N., Bates, A. S., Du, M., Hartenstein, V., Jefferis, G. S. X., & Funke, J. (2024). Neurotransmitter classification from electron microscopy images at synaptic sites in Drosophila melanogaster. Cell, 187(10), 2574–2594.e23.

23. Eichler, K., Hampel, S., Alejandro-García, A., Calle-Schuler, S. A., Santana-Cruz, A., Kmecova, L., Blagburn, J. M., Hoopfer, E. D., & Seeds, A. M. (2024). Somatotopic organization among parallel sensory pathways that promote a grooming sequence in. eLife, 12. 10.7554/eLife.87602

24. Fujiwara, T., Brotas, M., & Chiappe, M. E. (2022). Walking strides direct rapid and flexible recruitment of visual circuits for course control in Drosophila. Neuron, 110(13), 2124– 2138.e8.

25. Gerhard, S., Andrade, I., Fetter, R. D., Cardona, A., & Schneider-Mizell, C. M. (2017). Conserved neural circuit structure across larval development revealed by comparative connectomics. eLife, 6. 10.7554/eLife.29089

26. Golgi, C. (1873). Sulla struttura della grigia del cervello. Gaz. Med. Intalianna Lomb, 6, 244– 246.

27. Guo, L., Zhang, N., & Simpson, J. H. (2022). Descending neurons coordinate anterior grooming behavior in Drosophila. Current Biology: CB, 32(4), 823–833.e4.

28. Hampel, S., Franconville, R., Simpson, J. H., & Seeds, A. M. (2015). A neural command circuit for grooming movement control. eLife, 4, e08758.

29. Hoopfer, E. D., Jung, Y., Inagaki, H. K., Rubin, G. M., & Anderson, D. J. (2015). P1 interneurons promote a persistent internal state that enhances inter-male aggression in Drosophila. eLife, 4. 10.7554/eLife.11346

30. Ito, K., Shinomiya, K., Ito, M., Armstrong, J. D., Boyan, G., Hartenstein, V., Harzsch, S., Heisenberg, M., Homberg, U., Jenett, A., Keshishian, H., Restifo, L. L., Rössler, W., Simpson, J. H., Strausfeld, N. J., Strauss, R., Vosshall, L. B., & Insect Brain Name Working Group. (2014). A systematic nomenclature for the insect brain. Neuron, 81(4), 755–765.

31. Jenett, A., Rubin, G. M., Ngo, T.-T. B., Shepherd, D., Murphy, C., Dionne, H., Pfeiffer, B. D., Cavallaro, A., Hall, D., Jeter, J., Iyer, N., Fetter, D., Hausenfluck, J. H., Peng, H., Trautman, E. T., Svirskas, R. R., Myers, E. W., Iwinski, Z. R., Aso, Y., … Zugates, C. T. (2012). A GAL4-driver line resource for Drosophila neurobiology. Cell Reports, 2(4), 991–1001.

32. Kauer, I., Borst, A., & Haag, J. (2015). Complementary motion tuning in frontal nerve motor neurons of the blowfly. Journal of Comparative Physiology. A, Neuroethology, Sensory, Neural, and Behavioral Physiology, 201(4), 411–426.

33. Kohatsu, S., Koganezawa, M., & Yamamoto, D. (2011). Female contact activates male- specific interneurons that trigger stereotypic courtship behavior in Drosophila. Neuron, 69(3), 498–508.

34. Lappalainen, J. K., Tschopp, F. D., Prakhya, S., McGill, M., Nern, A., Shinomiya, K., Takemura, S.-Y., Gruntman, E., Macke, J. H., & Turaga, S. C. (2023). Connectome- constrained deep mechanistic networks predict neural responses across the fly visual system at single-neuron resolution. In bioRxiv (p. 2023.03.11.532232). 10.1101/2023.03.11.532232

35. Lee, S.-Y. J., Dallmann, C. J., Cook, A. P., Tuthill, J. C., & Agrawal, S. (2024). Divergent neural circuits for proprioceptive and exteroceptive sensing of the leg. bioRxiv : The Preprint Server for Biology. 10.1101/2024.04.23.590808

36. Lesser, E., Azevedo, A. W., Phelps, J. S., Elabbady, L., Cook, A., Sakeena Syed, D., Mark, B., Kuroda, S., Sustar, A., Moussa, A., Dallmann, C. J., Agrawal, S., Lee, S.-Y. J., Pratt, B., Skutt-Kakaria, K., Gerhard, S., Lu, R., Kemnitz, N., Lee, K., … Tuthill, J. C. (2024). Synaptic architecture of leg and wing premotor control networks in. bioRxiv : The Preprint Server for Biology. 10.1101/2023.05.30.542725

36a. Lillvis, J. L., Wang, K., Shiozaki, H. M., Xu, M., Stern, D. L., & Dickson, B. J. (n.d.). The neural basis of Drosophila courtship song. Manuscript in Preparation.

37. Lillvis, J. L., Wang, K., Shiozaki, H. M., Xu, M., Stern, D. L., & Dickson, B. J. (2024). Nested neural circuits generate distinct acoustic signals during Drosophila courtship. Current Biology: CB, 34(4), 808–824.e6.

38. Lima, S. Q., & Miesenböck, G. (2005). Remote control of behavior through genetically targeted photostimulation of neurons. Cell, 121(1), 141–152.

39. Li, M., Chen, D. S., Junker, I. P., Szorenyi, F., Chen, G. H., Berger, A. J., Comeault, A. A., Matute, D. R., & Ding, Y. (2023). Ancestral neural circuits potentiate the origin of a female sexual behavior. bioRxiv : The Preprint Server for Biology. 10.1101/2023.12.05.570174

40. Maitin-Shepard, J., Baden, A., Silversmith, W., Perlman, E., F., C., Blakely, T., Funke, J., Jordan, C., Falk, B., Kemnitz, N., C., T. R., Castro, M., Jagannathan, S., J., M. C., Hoag, A., Katz, B., Parsons, D., Wu, J., Kamentsky, L., … Roeder L, Li P. (2021). google/neuroglancer: Github. Com/google/neuroglancer, Retrieved, 04–30.

41. Mann, K., Gordon, M. D., & Scott, K. (2013). A pair of interneurons influences the choice between feeding and locomotion in Drosophila. Neuron, 79(4), 754–765.

42. Marin, E. C., Büld, L., Theiss, M., Sarkissian, T., Roberts, R. J. V., Turnbull, R., Tamimi, I. F. M., Pleijzier, M. W., Laursen, W. J., Drummond, N., Schlegel, P., Bates, A. S., Li, F., Landgraf, M., Costa, M., Bock, D. D., Garrity, P. A., & Jefferis, G. S. X. E. (2020). Connectomics Analysis Reveals First-, Second-, and Third-Order Thermosensory and Hygrosensory Neurons in the Adult Drosophila Brain. Current Biology: CB, 30(16), 3167–3182.e4.

43. Marin, E. C., Morris, B. J., Stuerner, T., Champion, A. S., Krzeminski, D., Badalamente, G., Gkantia, M., Dunne, C. R., Eichler, K., Shin-Takemura, Y., Tamimi, I. F. M., Fang, S., Moon, S. S., Cheong, H. S. J., Li, F., Schlegel, P., Berg, S., FlyEM Project Team, Card, G. M., … Jefferis, G. S. X. E. (2023). Systematic annotation of a complete adult male Drosophila nerve cord connectome reveals principles of functional organisation. bioRxiv.

44. McKellar, C. E., Lillvis, J. L., Bath, D. E., Fitzgerald, J. E., Cannon, J. G., Simpson, J. H., & Dickson, B. J. (2019). Threshold-Based Ordering of Sequential Actions during Drosophila Courtship. Current Biology: CB, 29(3), 426–434.e6.

45. Meissner, G. W., Nern, A., Dorman, Z., DePasquale, G. M., Forster, K., Gibney, T., Hausenfluck, J. H., He, Y., Iyer, N. A., Jeter, J., Johnson, L., Johnston, R. M., Lee, K., Melton, B., Yarbrough, B., Zugates, C. T., Clements, J., Goina, C., Otsuna, H., … FlyLight Project Team. (2023). A searchable image resource of GAL4 driver expression patterns with single neuron resolution. eLife, 12. 10.7554/eLife.80660

46. Meissner, G. W., Vannan, A., Jeter, J., Close, K., DePasquale, G. M., Dorman, Z., Forster, K., Beringer, J. A., Gibney, T. V., Hausenfluck, J. H., He, Y., Henderson, K., Johnson, L., Johnston, R. M., Ihrke, G., Iyer, N., Lazarus, R., Lee, K., Li, H.-H., … Rubin, G. M. (2024). A split-GAL4 driver line resource for Drosophila CNS cell types. In bioRxiv (p. 2024.01.09.574419). 10.1101/2024.01.09.574419

47. Mezzera, C., Brotas, M., Gaspar, M., Pavlou, H. J., Goodwin, S. F., & Vasconcelos, M. L. (2023). Ovipositor Extrusion Promotes the Transition from Courtship to Copulation and Signals Female Acceptance in Drosophila melanogaster. Current Biology: CB, 33(22), 5034.

48. Namiki, S., Dickinson, M. H., Wong, A. M., Korff, W., & Card, G. M. (2018). The functional organization of descending sensory-motor pathways in. eLife, 7. 10.7554/eLife.34272

49. Namiki, S., & Kanzaki, R. (2016). Comparative Neuroanatomy of the Lateral Accessory Lobe in the Insect Brain. Frontiers in Physiology, 7, 244.

50. Nern, A., Lösche, F., Takemura, S.-Y., Burnett, L. E., Dreher, M., Gruntman, E., Hoeller, J., Huang, G. B., Januszewski, M., Klapoetke, N. C., Koskela, S., Longden, K. D., Lu, Z., Preibisch, S., Qiu, W., Rogers, E. M., Seenivasan, P., Zhao, A., Bogovic, J., … Reiser, M. B. (2024). Connectome-driven neural inventory of a complete visual system. bioRxiv : The Preprint Server for Biology. 10.1101/2024.04.16.589741

51. Pavlou, H. J., & Goodwin, S. F. (2013). Courtship behavior in Drosophila melanogaster: towards a “courtship connectome.” Current Opinion in Neurobiology, 23(1), 76–83.

52. Phelps, J. S., Hildebrand, D. G. C., Graham, B. J., Kuan, A. T., Thomas, L. A., Nguyen, T. M., Buhmann, J., Azevedo, A. W., Sustar, A., Agrawal, S., Liu, M., Shanny, B. L., Funke, J., Tuthill, J. C., & Lee, W.-C. A. (2021). Reconstruction of motor control circuits in adult Drosophila using automated transmission electron microscopy. Cell, 184(3), 759–774.e18.

53. Possidente, D. R., & Murphey, R. K. (1989). Genetic control of sexually dimorphic axon morphology in Drosophila sensory neurons. Developmental Biology, 132(2), 448–457.

54. Rayshubskiy, A., Holtz, S. L., D’Alessandro, I., Li, A. A., Vanderbeck, Q. X., Haber, I. S., Gibb, P. W., & Wilson, R. I. (2020). Neural circuit mechanisms for steering control in walking Drosophila. In bioRxiv (p. 2020.04.04.024703). 10.1101/2020.04.04.024703

55. Sapkal, N., Mancini, N., Kumar, D. S., Spiller, N., Murakami, K., Vitelli, G., Bargeron, B., Maier, K., Eichler, K., Jefferis, G. S. X., Shiu, P. K., Sterne, G. R., & Bidaye, S. S. (2023). Neural circuit mechanisms underlying context-specific halting in Drosophila. In bioRxiv (p. 2023.09.25.559438). 10.1101/2023.09.25.559438

56. Scheffer, L. K., Xu, C. S., Januszewski, M., Lu, Z., Takemura, S.-Y., Hayworth, K. J., Huang, G. B., Shinomiya, K., Maitlin-Shepard, J., Berg, S., Clements, J., Hubbard, P. M., Katz, W. T., Umayam, L., Zhao, T., Ackerman, D., Blakely, T., Bogovic, J., Dolafi, T., … Plaza, S. M. (2020). A connectome and analysis of the adult Drosophila central brain. eLife, 9. 10.7554/eLife.57443

57. Schlegel, P., Bates, A. S., Stürner, T., Jagannathan, S. R., Drummond, N., Hsu, J., Serratosa Capdevila, L., Javier, A., Marin, E. C., Barth-Maron, A., Tamimi, I. F., Li, F., Rubin, G. M., Plaza, S. M., Costa, M., & Jefferis, G. S. X. E. (2021). Information flow, cell types and stereotypy in a full olfactory connectome. eLife, 10. 10.7554/eLife.66018

58. Schlegel, P., Yin, Y., Bates, A. S., Dorkenwald, S., Eichler, K., Brooks, P., Han, D. S., Gkantia, M., Dos Santos, M., Munnelly, E. J., Badalamente, G., Capdevila, L. S., Sane, V. A., Pleijzier, M. W., Tamimi, I. F. M., Dunne, C. R., Salgarella, I., Javier, A., Fang, S., … Jefferis, G. S. X. E. (2023). Whole-brain annotation and multi-connectome cell typing quantifies circuit stereotypy in. bioRxiv : The Preprint Server for Biology. 10.1101/2023.06.27.546055

59. Seeds, A. M., Ravbar, P., Chung, P., Hampel, S., Midgley, F. M., Jr, Mensh, B. D., & Simpson, J. H. (2014). A suppression hierarchy among competing motor programs drives sequential grooming in Drosophila. eLife, 3, e02951.

60. Shiozaki, H. M., Wang, K., Lillvis, J. L., Xu, M., Dickson, B., & Stern, D. (2023). Combinatorial circuit dynamics orchestrate flexible motor patterns in Drosophila. bioRxiv. 10.1101/2022.12.14.520499

61. Shirangi, T. R., Wong, A. M., Truman, J. W., & Stern, D. L. (2016). Doublesex Regulates the Connectivity of a Neural Circuit Controlling Drosophila Male Courtship Song. Developmental Cell, 37(6), 533–544.

62. Simpson, J. H. (2024). Descending control of motor sequences in Drosophila. Current Opinion in Neurobiology, 84, 102822.

63. Steinbeck, F., Adden, A., & Graham, P. (2020). Connecting brain to behaviour: a role for general purpose steering circuits in insect orientation? The Journal of Experimental Biology, 223(Pt 5). 10.1242/jeb.212332

64. Strausfeld, N. J., Seyan, H. S., & Milde, J. J. (1987). The neck motor system of the flyCalliphora erythrocephala. Journal of Comparative Physiology A, 160(2), 205–224.

65. Suver, M. P., Huda, A., Iwasaki, N., Safarik, S., & Dickinson, M. H. (2016). An Array of Descending Visual Interneurons Encoding Self-Motion in Drosophila. The Journal of Neuroscience: The Official Journal of the Society for Neuroscience, 36(46), 11768– 11780.

66. Takemura, S.-Y., Huang, G. B., Januszewski, M., Lu, Z., Marin, E. C., Preibisch, S., Xu, C. S., Champion, A., Cheong, H. S. J., Costa, M., Eichler, K., Knecht, C. J., Li, F., Morris, B. J., Schlegel, P., Stuerner, T., Badalamente, G., Bogovic, J., Brooks, P., … Berg, S. (2023). A Connectome of the Male Drosophila Ventral Nerve Cord. bioRxiv.

67. Tirian, L., & Dickson, B. J. (2017). The VT GAL4, LexA, and split-GAL4 driver line collections for targeted expression in the Drosophila nervous system. In bioRxiv (p. 198648). 10.1101/198648

68. Tsubouchi, A., Yano, T., Yokoyama, T. K., Murtin, C., Otsuna, H., & Ito, K. (2017). Topological and modality-specific representation of somatosensory information in the fly brain. Science, 358(6363), 615–623.

69. Valdes-Aleman, J., Fetter, R. D., Sales, E. C., Heckman, E. L., Venkatasubramanian, L., Doe, C. Q., Landgraf, M., Cardona, A., & Zlatic, M. (2021). Comparative Connectomics Reveals How Partner Identity, Location, and Activity Specify Synaptic Connectivity in Drosophila. Neuron, 109(1), 105–122.e7.

70. von Philipsborn, A. C., Liu, T., Yu, J. Y., Masser, C., Bidaye, S. S., & Dickson, B. J. (2011). Neuronal control of Drosophila courtship song. Neuron, 69(3), 509–522.

71. von Reyn, H. C., Jang, H., Eichler, K., Stuerner, T., Costa, M., Ausborn, J., & Catherine, R. (2024 unpublished). A circuit underlying evasive flight maneuvers centered around the descending neuron DNp03.

72. Wang, F., Wang, K., Forknall, N., Parekh, R., & Dickson, B. J. (2020b). Circuit and Behavioral Mechanisms of Sexual Rejection by Drosophila Females. Current Biology: CB, 30(19), 3749–3760.e3.

73. Wang, F., Wang, K., Forknall, N., Patrick, C., Yang, T., Parekh, R., Bock, D., & Dickson, B. J. (2020a). Neural circuitry linking mating and egg laying in Drosophila females. Nature, 579(7797), 101–105.

74. Wang, K., Wang, F., Forknall, N., Yang, T., Patrick, C., Parekh, R., & Dickson, B. J. (2021). Neural circuit mechanisms of sexual receptivity in Drosophila females. Nature, 589(7843), 577–581.

75. Winding, M., Pedigo, B. D., Barnes, C. L., Patsolic, H. G., Park, Y., Kazimiers, T., Fushiki, A., Andrade, I. V., Khandelwal, A., Valdes-Aleman, J., Li, F., Randel, N., Barsotti, E., Correia, A., Fetter, R. D., Hartenstein, V., Priebe, C. E., Vogelstein, J. T., Cardona, A., & Zlatic, M. (2023). The connectome of an insect brain. Science, 379(6636), eadd9330.

76. Witvliet, D., Mulcahy, B., Mitchell, J. K., Meirovitch, Y., Berger, D. R., Wu, Y., Liu, Y., Koh, W. X., Parvathala, R., Holmyard, D., Schalek, R. L., Shavit, N., Chisholm, A. D., Lichtman, J. W., Samuel, A. D. T., & Zhen, M. (2021). Connectomes across development reveal principles of brain maturation. Nature, 596(7871), 257–261.

77. Yang, H. H., Brezovec, L. E., Capdevila, L. S., Vanderbeck, Q. X., Adachi, A., Mann, R. S., & Wilson, R. I. (2023). Fine-grained descending control of steering in walking Drosophila. bioRxiv : The Preprint Server for Biology. 10.1101/2023.10.15.562426

78. Yu, J. Y., Kanai, M. I., Demir, E., Jefferis, G. S. X., & Dickson, B. J. (2010). Cellular Organization of the Neural Circuit that Drives Drosophila Courtship Behavior. In Current Biology (Vol. 20, Issue 18, pp. 1602–1614). 10.1016/j.cub.2010.08.025

79. Zheng, Z., Lauritzen, J. S., Perlman, E., Robinson, C. G., Nichols, M., Milkie, D., Torrens, O., Price, J., Fisher, C. B., Sharifi, N., Calle-Schuler, S. A., Kmecova, L., Ali, I. J., Karsh, B., Trautman, E. T., Bogovic, J. A., Hanslovsky, P., Jefferis, G. S. X. E., Kazhdan, M., … Bock, D. D. (2018). A Complete Electron Microscopy Volume of the Brain of Adult Drosophila melanogaster. Cell, 174(3), 730–743.e22.

