## Supplementary figures and images for "Comparative connectomics of the descending and ascending neurons of the *Drosophila* nervous system: stereotypy and sexual dimorphism"

### Extended_Data_Fig1_formatted600.pdf

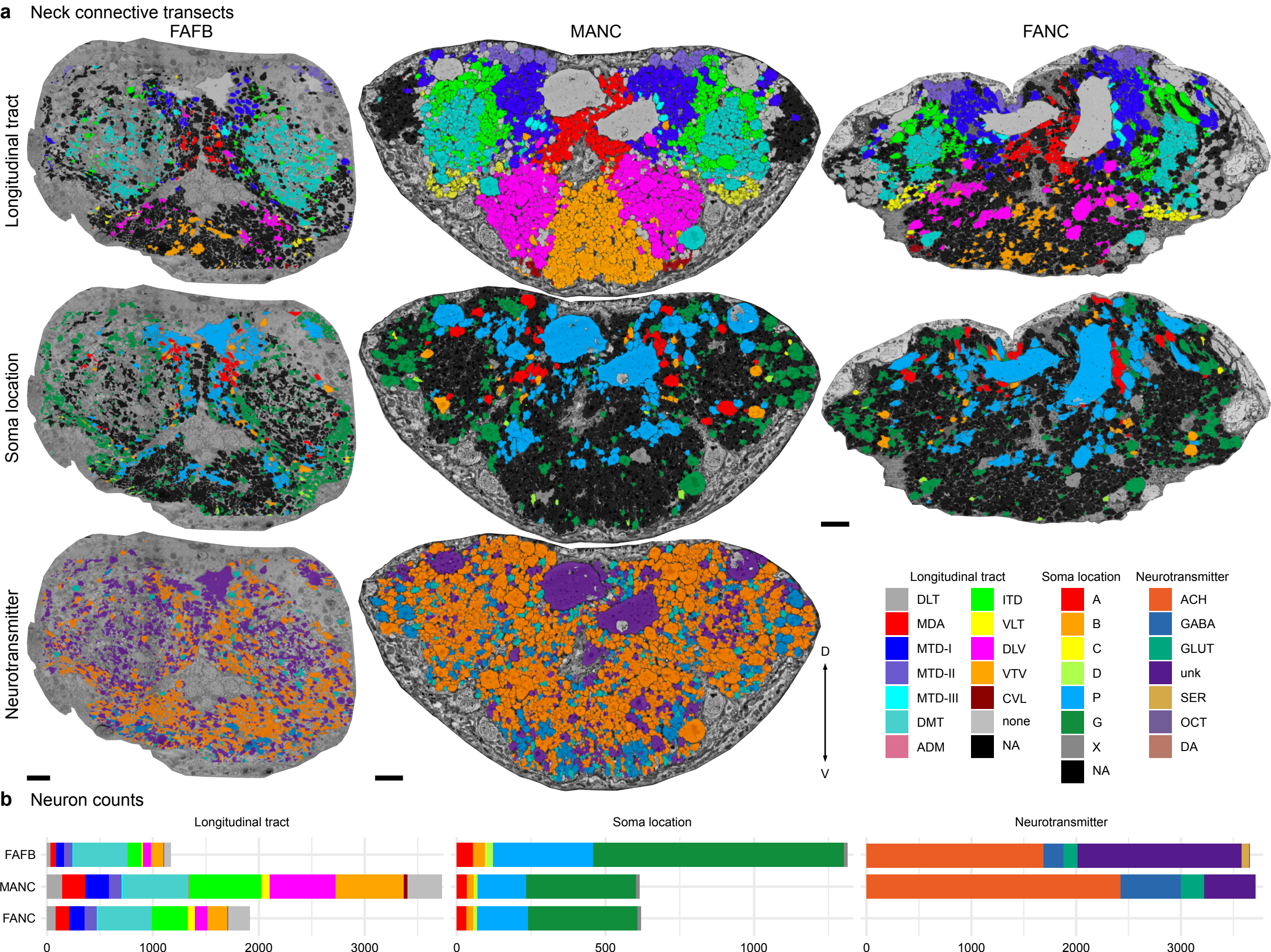

### Extended_Data_Fig2_formatted600.pdf

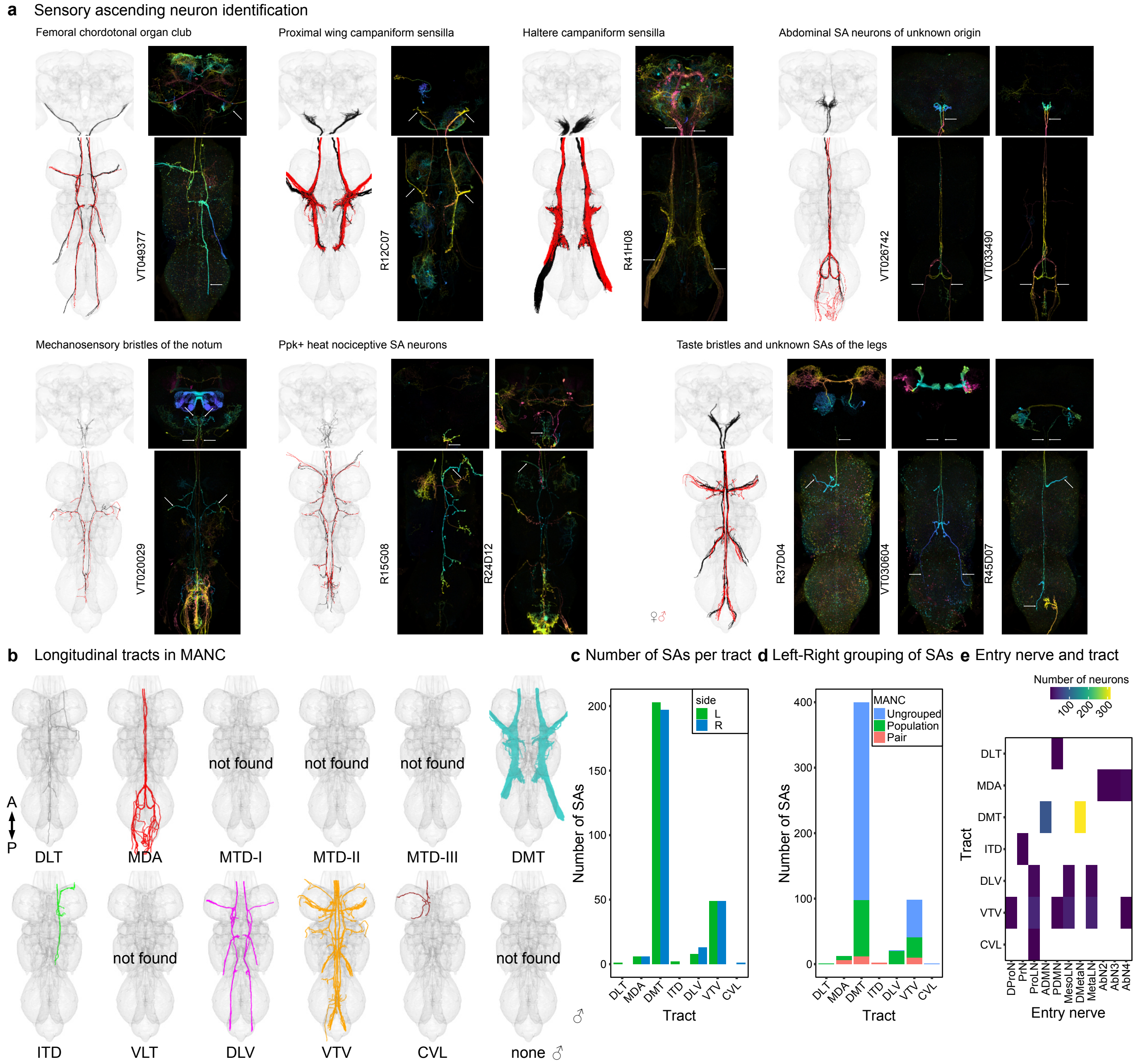

### Extended_Data_Fig3_formatted600.pdf

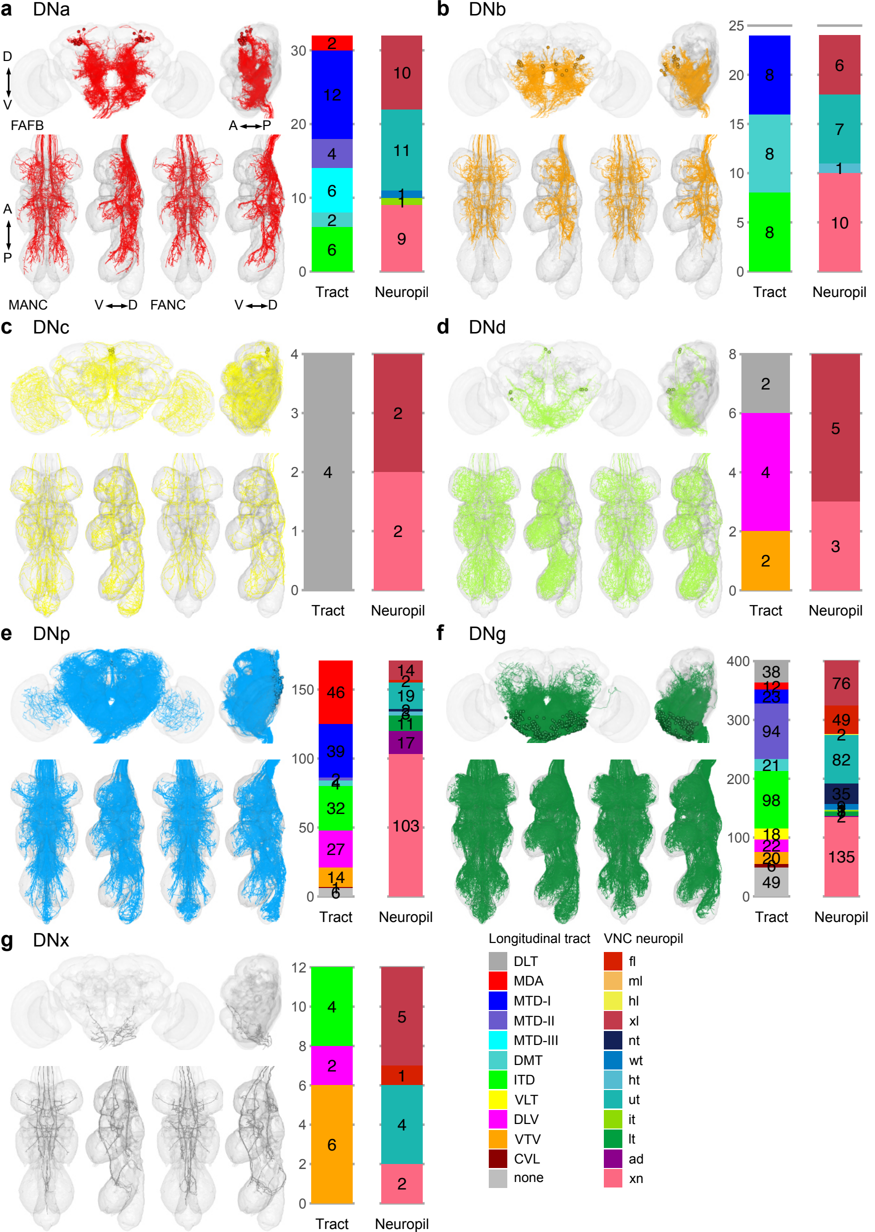

### Extended_Data_Fig4_formatted600.pdf

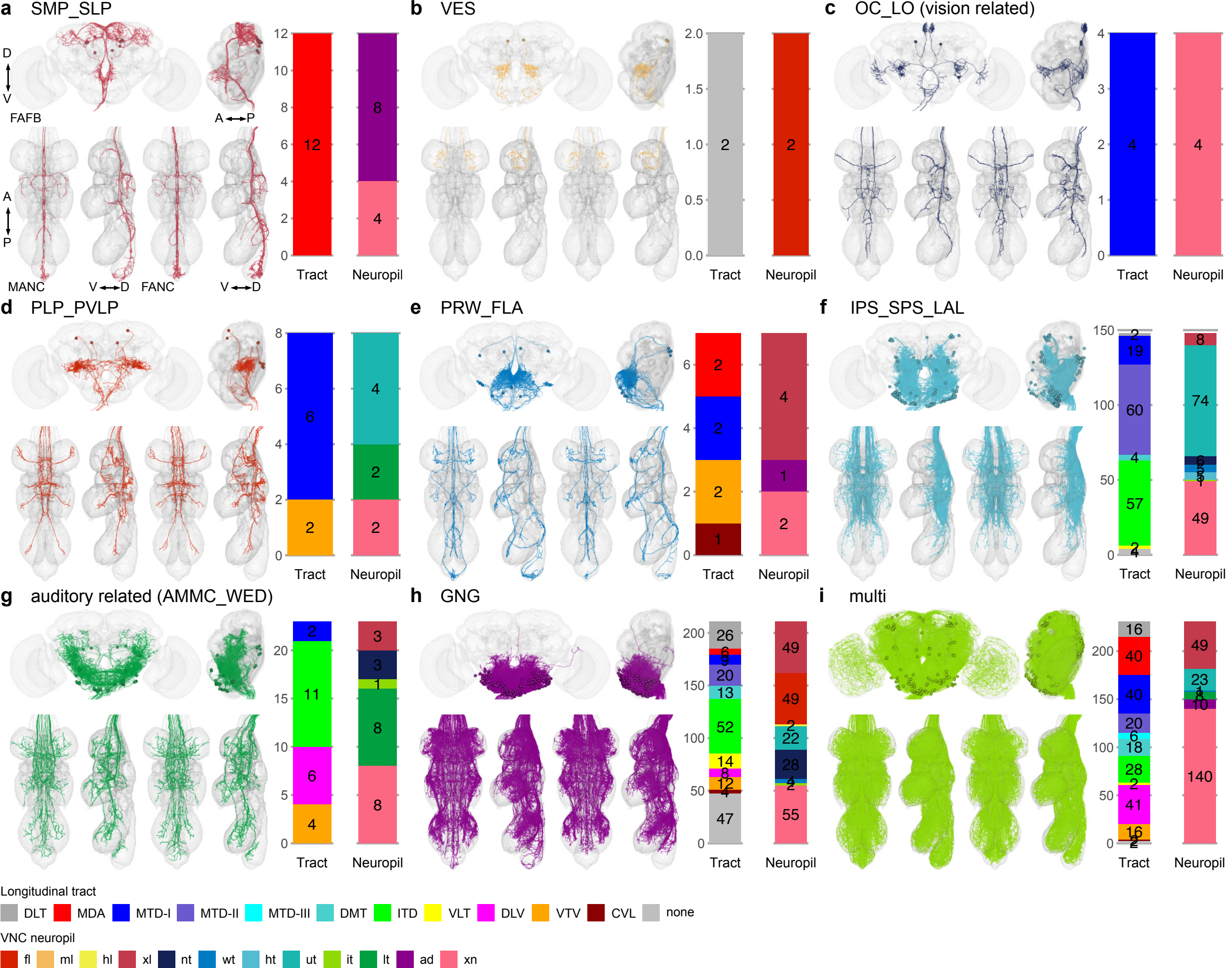

### Extended_Data_Fig5_formatted600.pdf

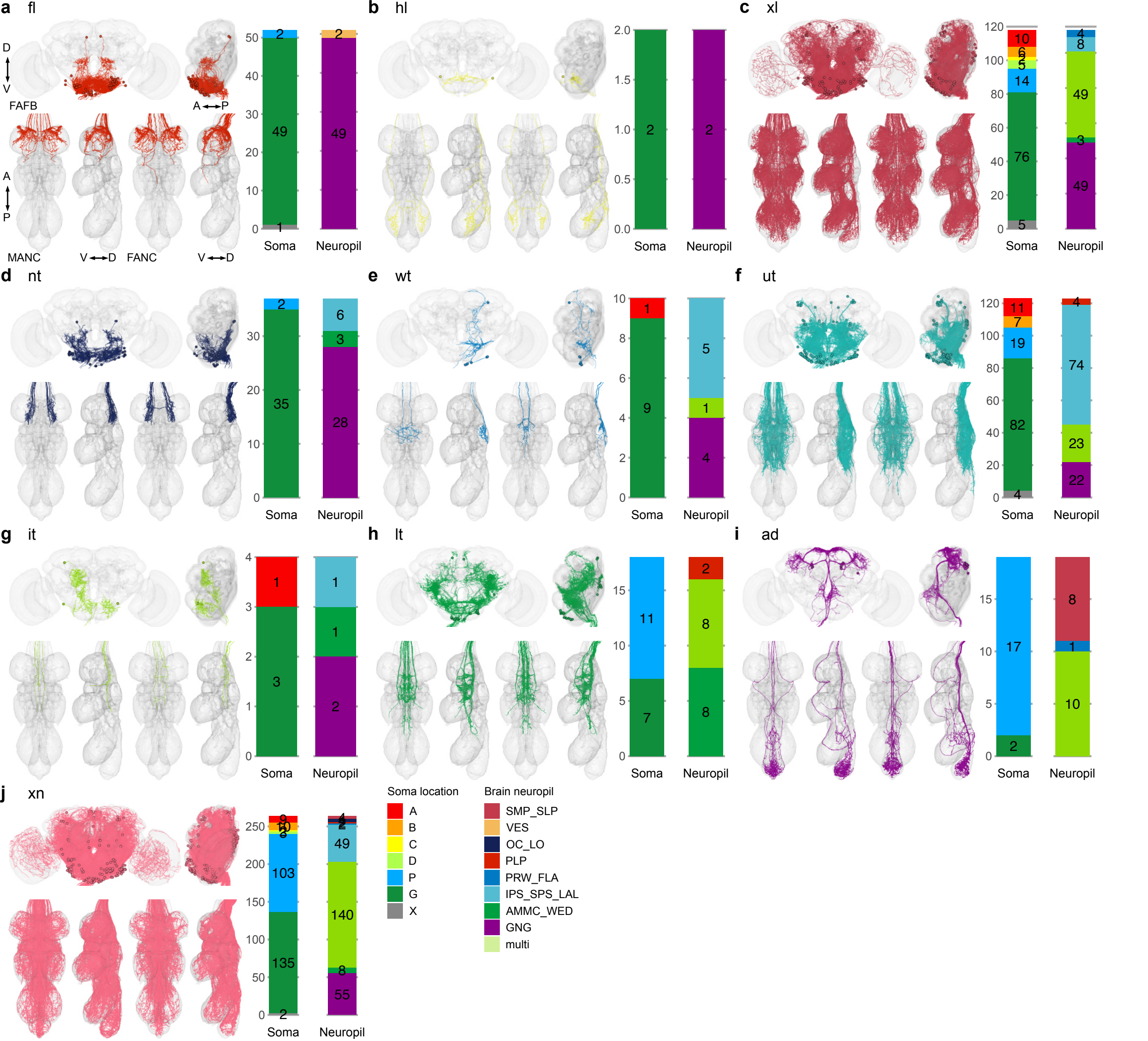

### Extended_Data_Fig6_formatted600.pdf

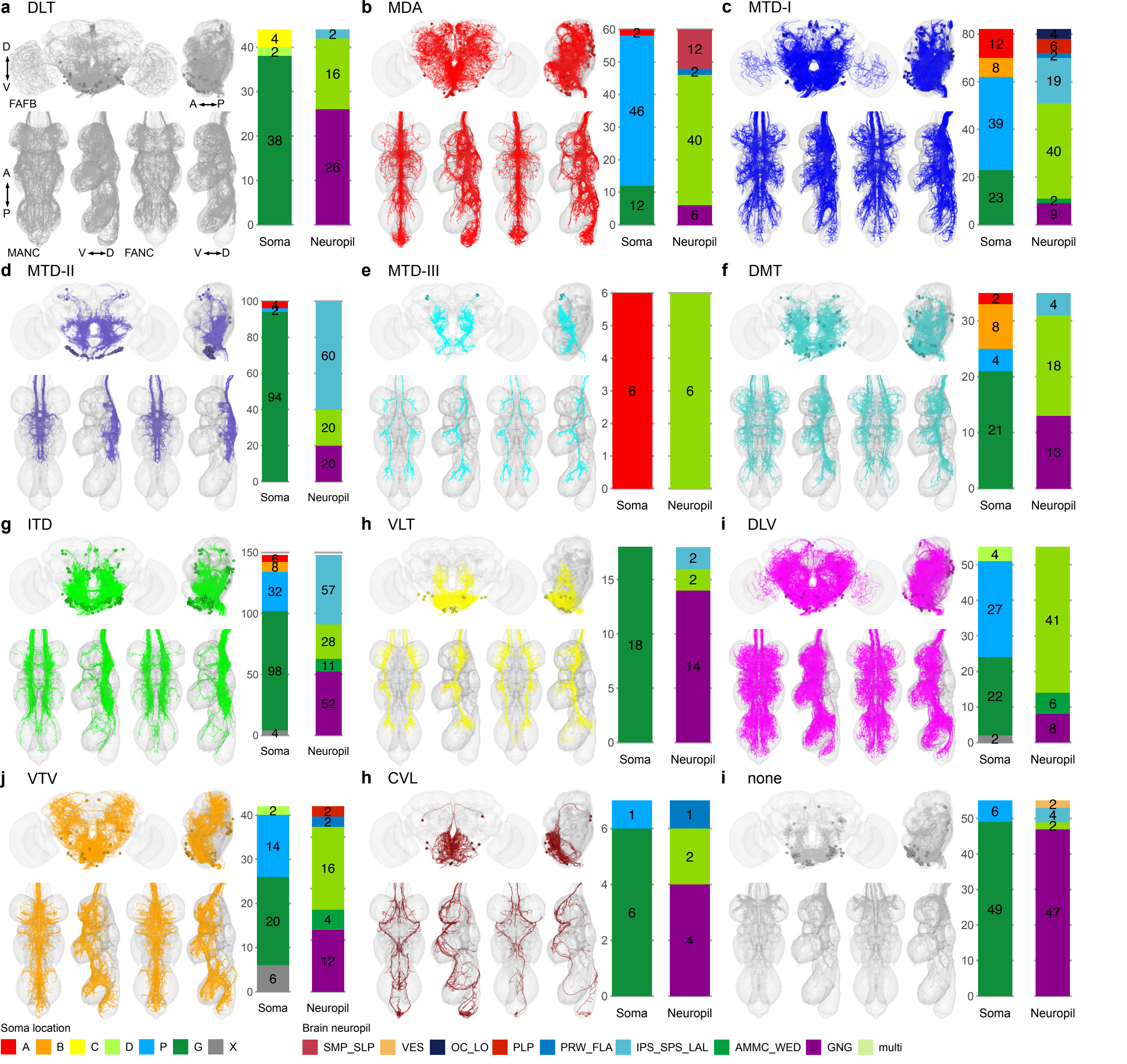

### Extended_Data_Fig7_Namiki_DNs_formatted1200.pdf

#### Canonical DNs from Namiki et al., 2018

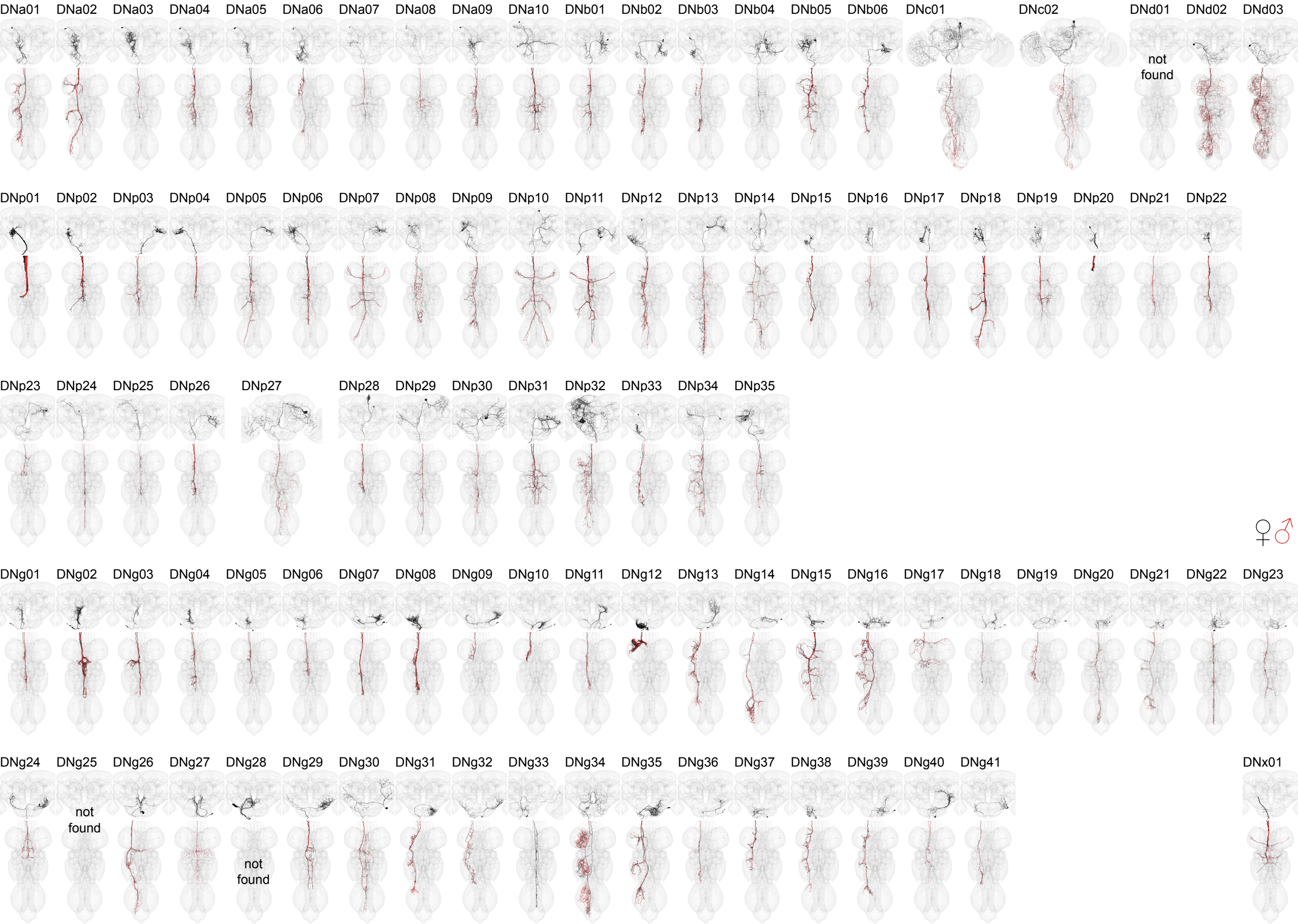

### Extended_Data_Fig8_formatted1200.pdf

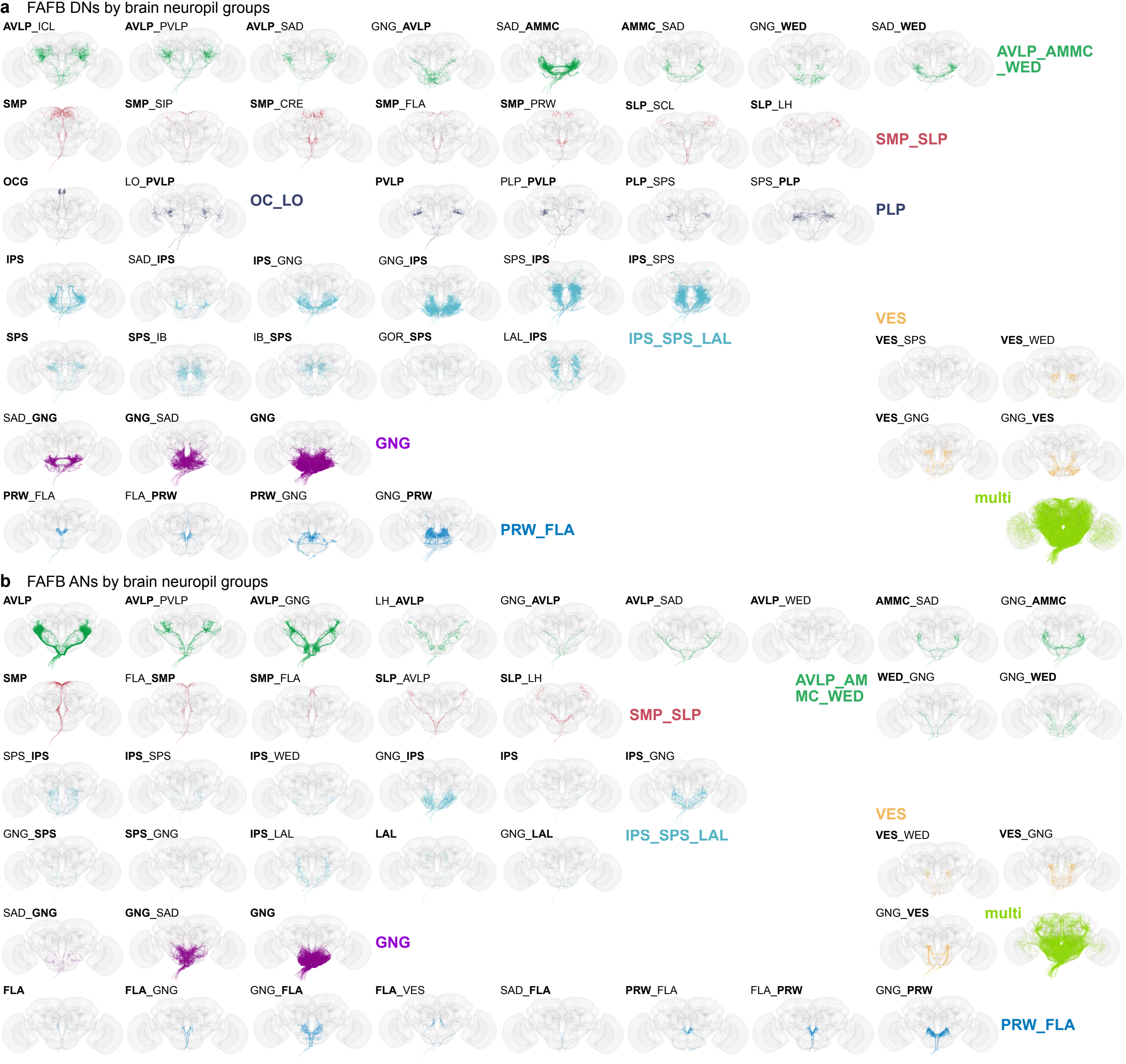

### Extended_Data_Fig9_formatted600.pdf

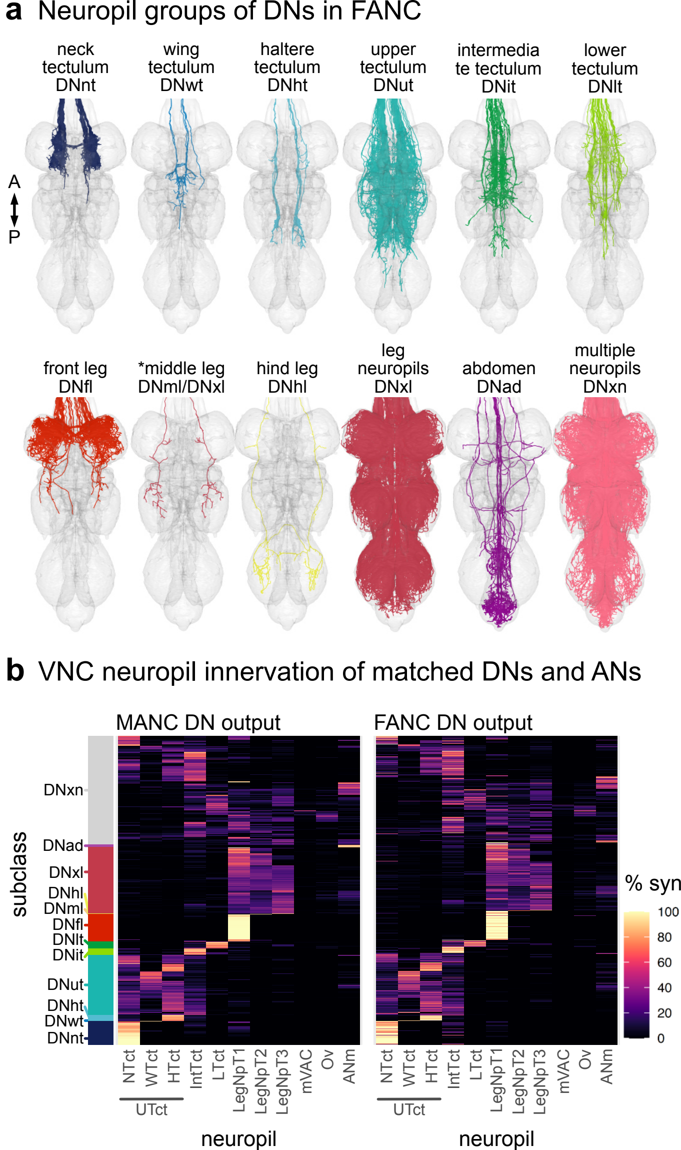

### Extended_Data_Fig10_formatted600.pdf

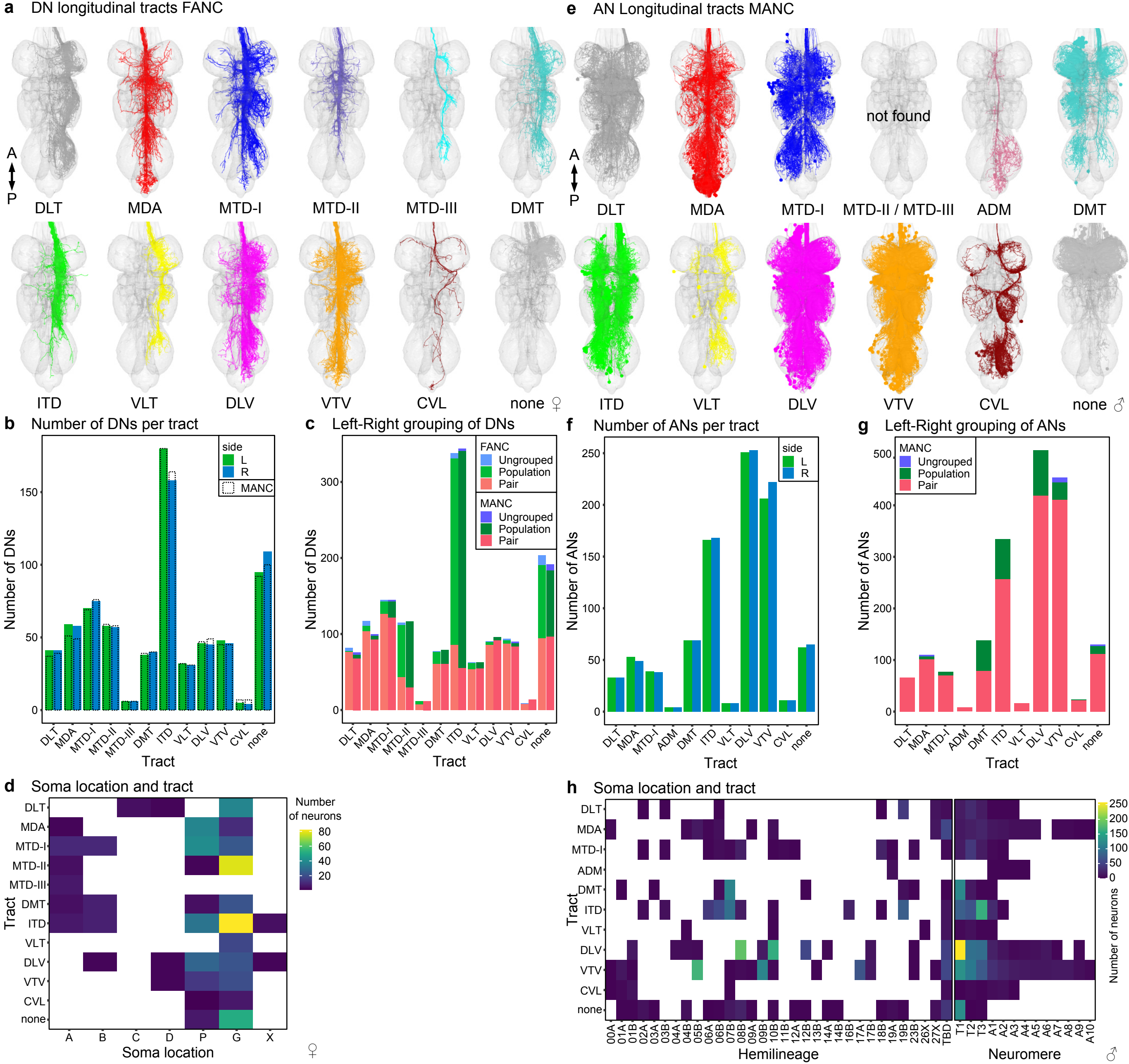

### Extended_Data_Fig11_formatted1200.pdf

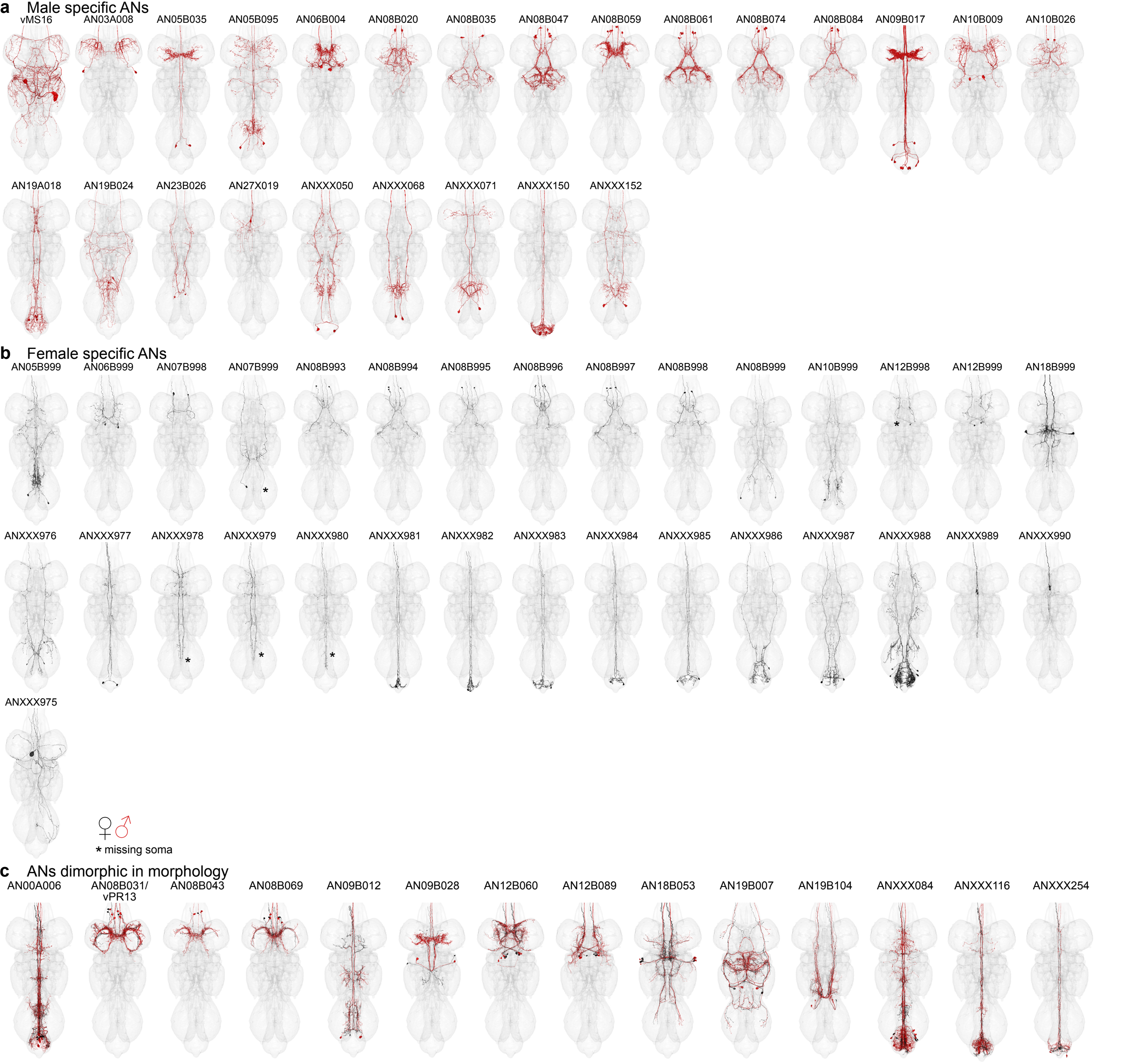

### Fig1-Overview_formatted600.pdf

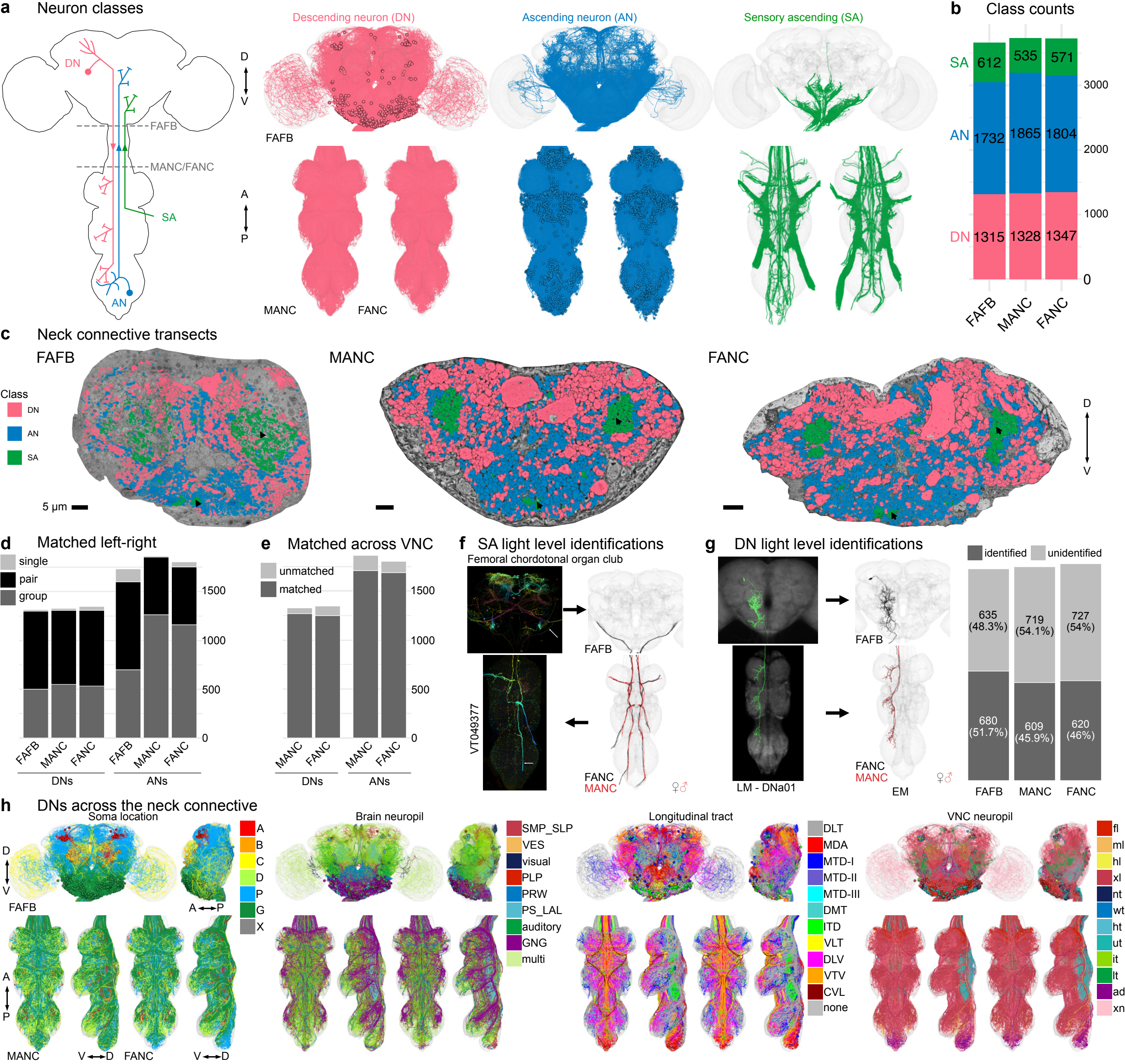

### Fig2-Namiki2_DNs_formatted_1200.pdf

Canonical DNs from Namiki et al., 2024

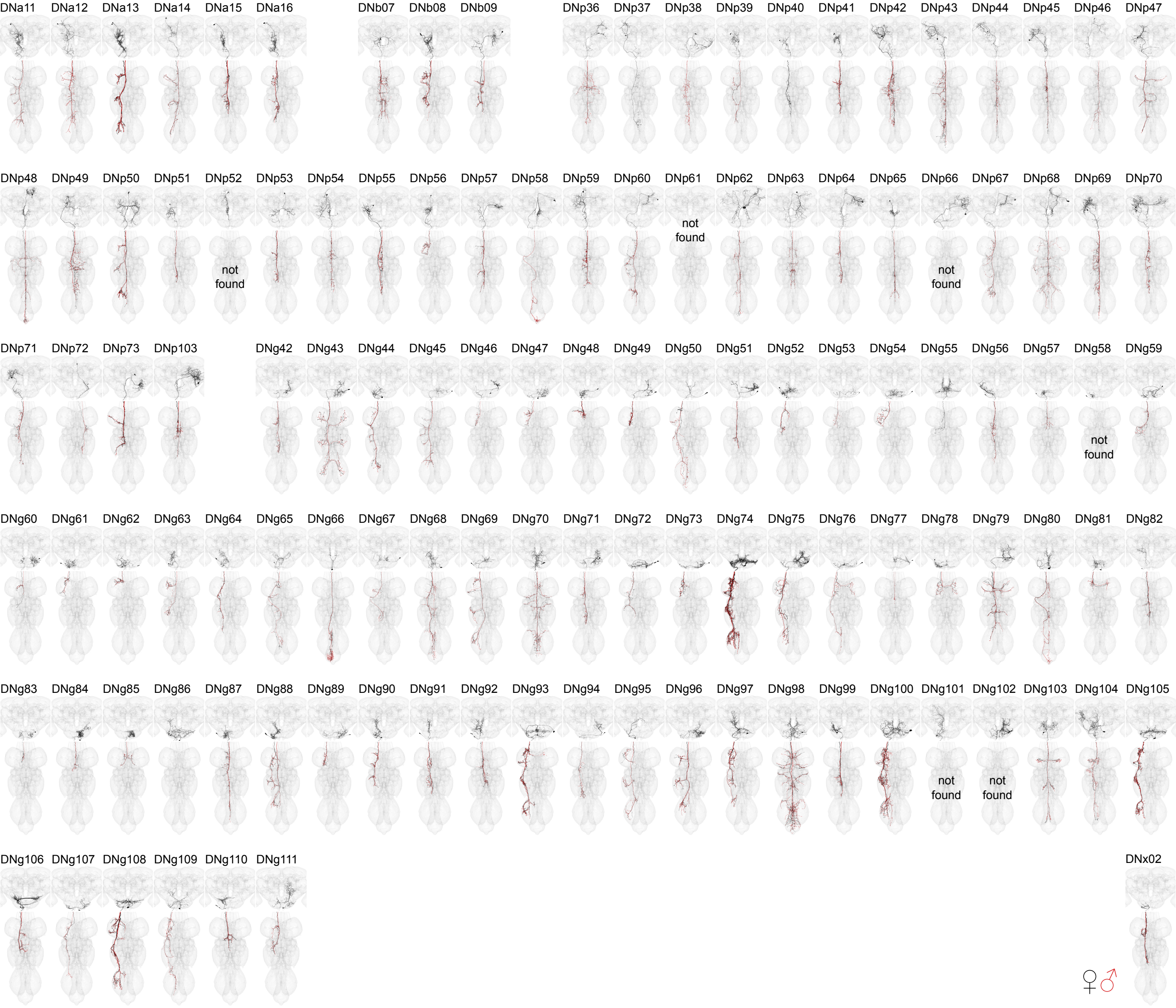

### Fig3-sensory_ranking_formatted600.pdf

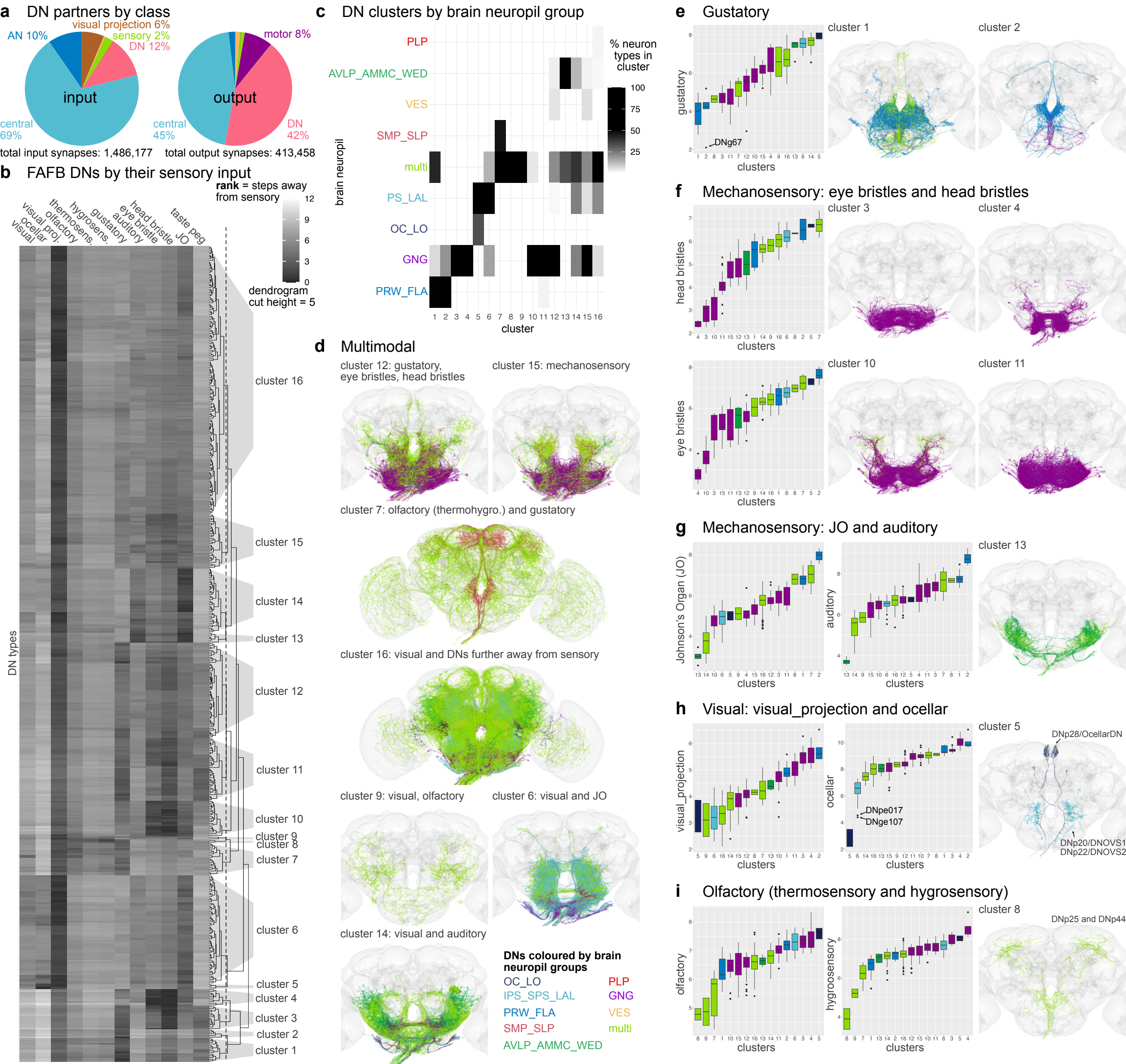

### Fig4-DNx02_formatted600.pdf

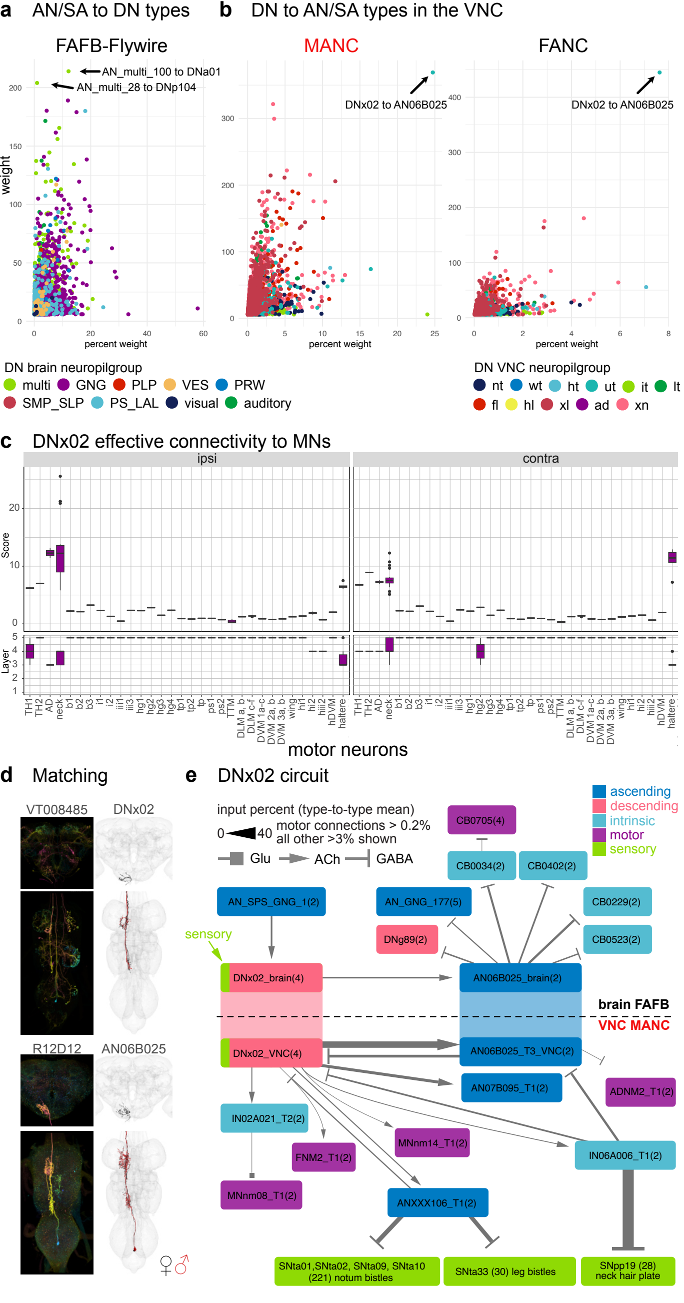

### Fig5-stereotypy_formatted600.pdf

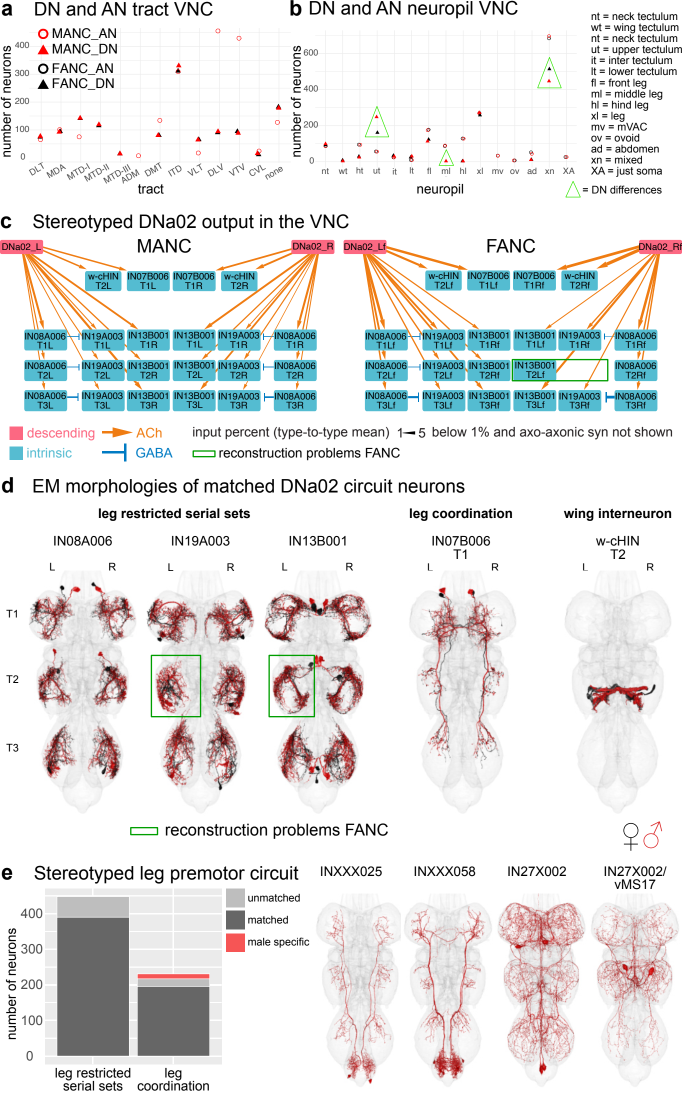

### Fig6-dimorphism_formatted600.pdf

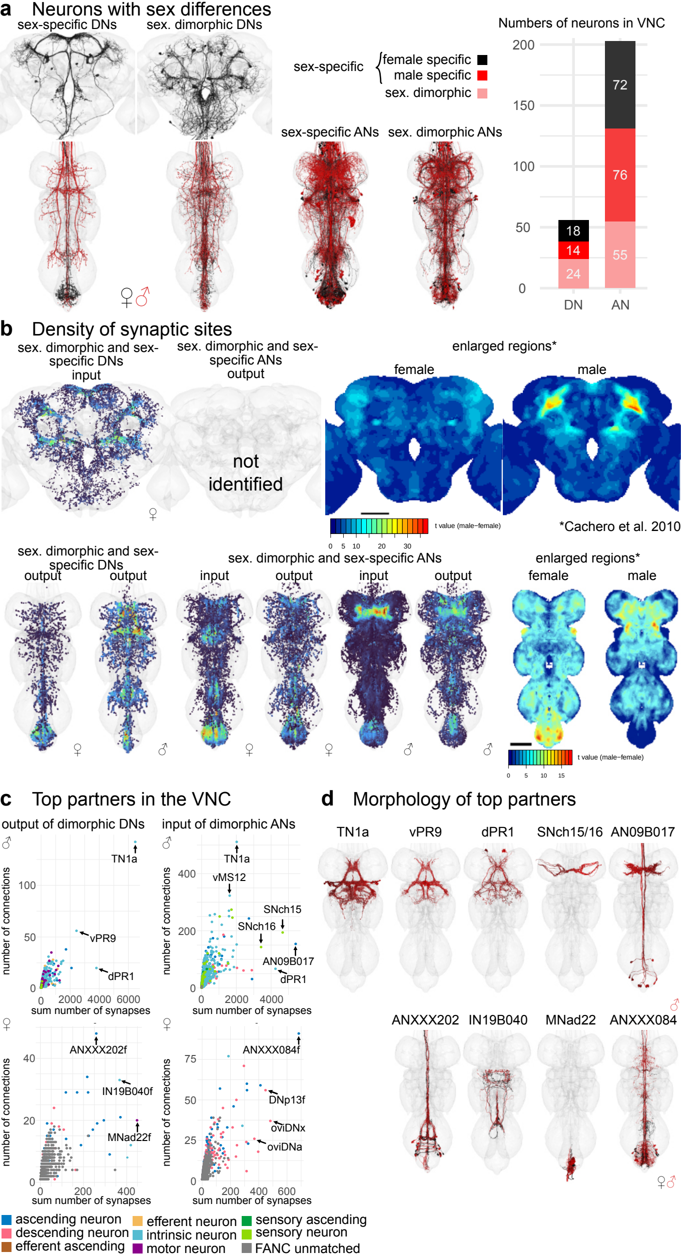

### Fig7-dimorphic_DNs_formatted1200.pdf

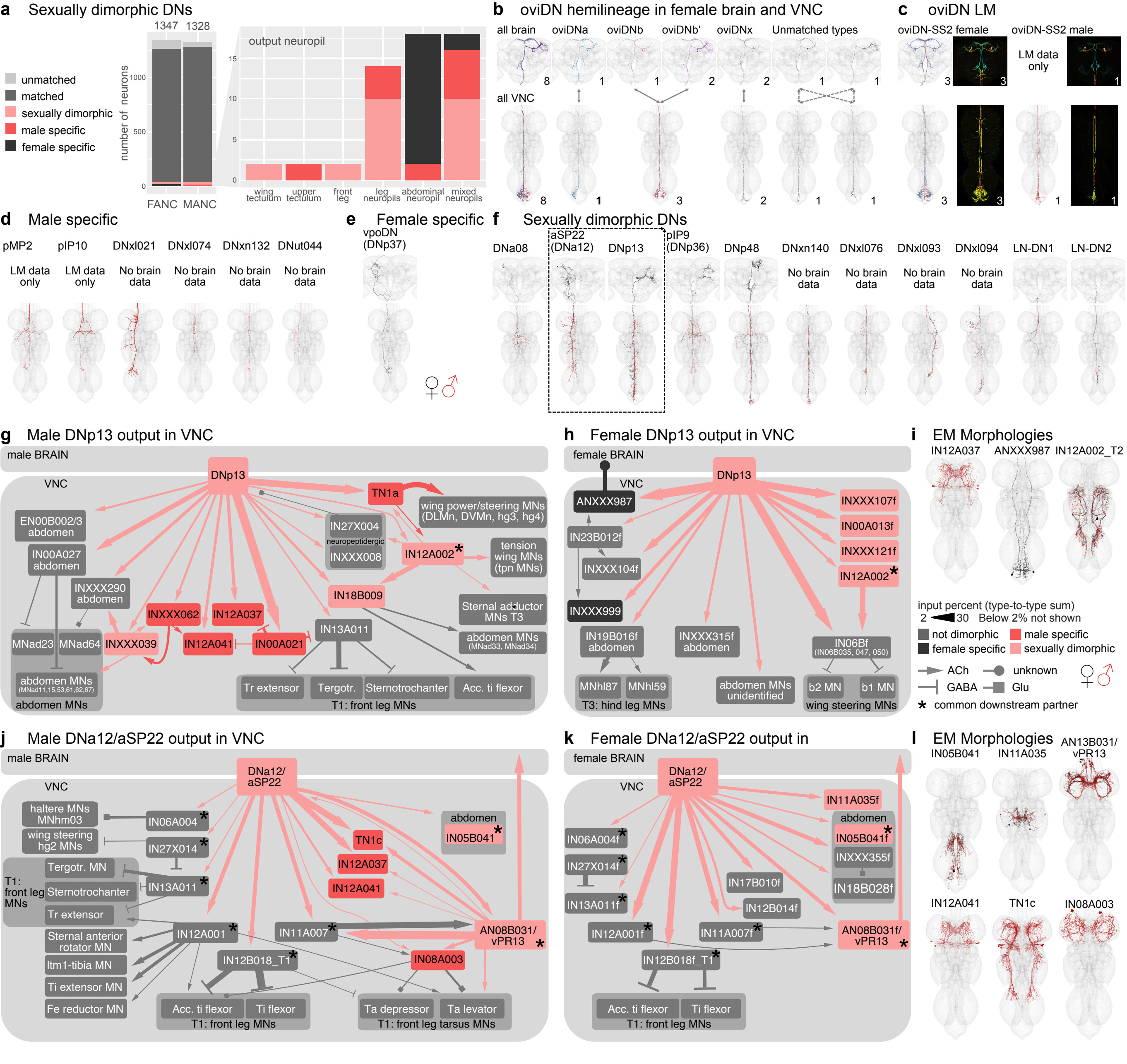

### Fig8-dimorphic_ANs_formatted1200.pdf

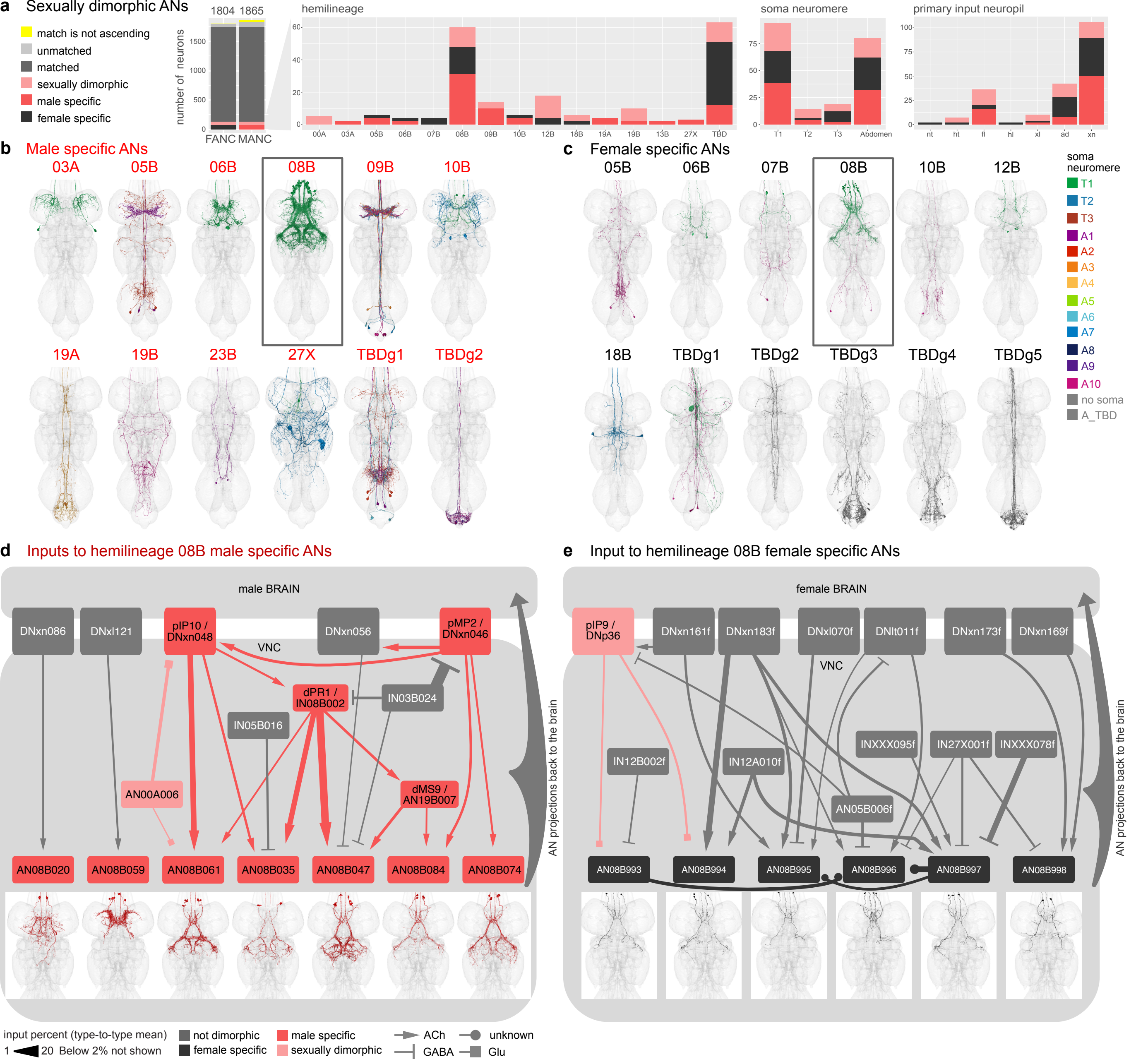
